# Individual dopaminergic neurons induce unique, yet overlapping combinations of behavioural modulations including safety learning, memory retrieval and acute locomotion

**DOI:** 10.1101/2025.01.23.634646

**Authors:** Naoko Toshima, Arman Behrad, Franziska Behnke, Gauri Kaushik, Aliće Weiglein, Martin Strauch, Juliane Thoener, Oliver Kobler, Maia Lisandra M. Wang, Ayaka Fukushima, Markus Dörr, Michael Schleyer

## Abstract

Two evolutionary highly conserved functions of dopamine are to carry “teaching” signals during associative learning and to control movement. In mammals and humans, these functions are generally thought to be produced by different populations of neurons. Here, we investigated in the larva of *Drosophila melanogaster* whether both these functions can be induced by the same individual dopaminergic neurons in the central brain. Focusing on the dopaminergic neurons of the DL1-cluster, we asked whether the optogenetic activation of individual neurons established associative punishment and/or safety memories, controlled the retrieval of the established memories, and acutely modulated locomotion. We found that each neuron had a unique, yet overlapping set of behavioural effects. Several individual neurons both established a memory and modulated acute locomotion by increasing the animals’ bending and decreasing its velocity. Our results demonstrate that individual dopaminergic neurons can fulfil a surprisingly broad range of functions in different behavioural contexts. Given the highly conserved roles of the dopaminergic system across the animal kingdom, this study raises the question whether a similarly diverse functionality can be found also in other animals, including humans.

## Introduction

Dopamine has many functions in the brain. Among them, two turned out to be highly conserved throughout the animal kingdom: to carry “teaching” signals during associative learning and to initiate and control movement^1–5^. In mammals and humans, it is widely accepted that these two functions are brought about by different populations of dopaminergic neurons (DANs): effects regarding learning are thought to be elicited by the mesolimbic pathway, effects regarding movement by the nigro-stratiatal pathway^4^. Here, we study in the larva of the fruit fly *Drosophila melanogaster* whether in contrast the same individual central brain-DANs can induce both functions regarding memory and movement.

The fruit fly larva is an established model organism to study the neuronal and genetic mechanisms underlying associative learning^6–7^. Due to the numerical simplicity of its brain and the available toolbox for transgene expression, the larva allows investigating these mechanisms at tractable levels of complexity. In *D. melanogaster*, distinct sets of DANs convey appetitive and aversive unconditioned stimuli (US), respectively (larvae^6–7^; adults^2,8^) (a similar scenario may be emerging in vertebrates, too^1,9–13^). Each of the DANs synapses onto a different compartment of the parallel fibres of Kenyon cells (KCs) that form a higher-order brain structure called mushroom body (larvae^7^; adults^14^). Odours are combinatorically coded across the array of these KCs which in turn give output to mushroom body output neurons (MBONs). The MBONs collect information across the KCs and send it towards efferent circuitry, promoting either approach or avoidance (larvae^7^; adults^14^). Within the KCs, the coincidence of odour presentation and activity in DANs is detected and enables a memory trace to be formed. The respective memory engram is thought to come by synaptic depression between the KCs and the MBONs of the same mushroom body compartment, such that the net effect of the activity across the MBONs is tilted towards approach or avoidance, respectively (larvae^15–16^; adults^17–18^). Electron microscope reconstructions further revealed a direct within-compartment connection from the DANs onto the MBONs (larvae^19^; adults^20^). Thus, each DAN has two main targets: the KCs and the MBONs of its own compartment. The general organization of this DAN-KC-MBON matrix is likely shared with other insects^2,21–23^.

Three pairs of identified DANs of the dorso-lateral cluster 1 (DL1) have been shown to support aversive learning upon optogenetic activation during Pavlovian conditioning^15,24–25^. In this study, we ask whether those same DANs have additional functions. Specifically, we ask for their role for safety learning – that is, learning that a cue predicts the absence of a threat; memory retrieval – that is, whether a memory will be retrieved and affect behaviour; and modulations of acute, innate locomotion. Little is known about how the dopaminergic system controls any of these functions – if at all. By systematically testing the functions of the same DANs across diverse behavioural tasks, we investigate the fundamental organization of the dopaminergic system: do different individual neurons specialize in each function, or does a single neuron serve multiple functions? Given the highly conserved functionality of the dopaminergic system across the animal kingdom, this study has the potential to provide valuable insights into the dopaminergic organisation of behaviour in any brain.

## Results

We focus our study on the DANs that support aversive learning when optogenetically activated, called DAN-d1, DAN-f1 and DAN-g1. We investigated the transgenic expression pattern of the driver strains covering each DAN, and the effects of activating each DAN on learning, memory retrieval and locomotion.

### The used driver strains drive transgene expression in identified DANs

Previous studies had described the expression patterns of the used split-driver strains in the central nervous system in detail^15,25–27^. Here, we extended the view on the expression within the whole body. Towards this end, we applied a recent protocol for whole-body visualizations^28^. We confirmed the highly specific transgene expression in the brain of the four split-Gal4 driver strains without any detectable expression in the rest of the body (Fig. 1A-D, Supplementary material S1). *TH-Gal4* is known to cover almost all DANs in the brain, with the notable exception of the neurons of the primary protocerebral anterior medial (pPAM) cluster^26^. We confirm this pattern of transgene expression and in addition found strong expression outside of the brain, in particular in the proventriculus and in hypodermal melanin-producing cells (Fig. 1E, Supplementary Movie 1). This pattern fits to the reported function of dopamine in the hardening and darkening of the insect cuticle^29–30^ and a recent report of *TH-Gal4* expression in the adult proventriculus^31^. This result means that any genetic manipulation using *TH-Gal4* may affect cells outside of the central nervous system. Finally, we confirm that *R58E02-Gal4* covers three of the four pPAM DANs that are missing in the *TH-Gal4* expression pattern^27^, with no expression outside of the central nervous system (Fig. 1F).

**Figure 1:**
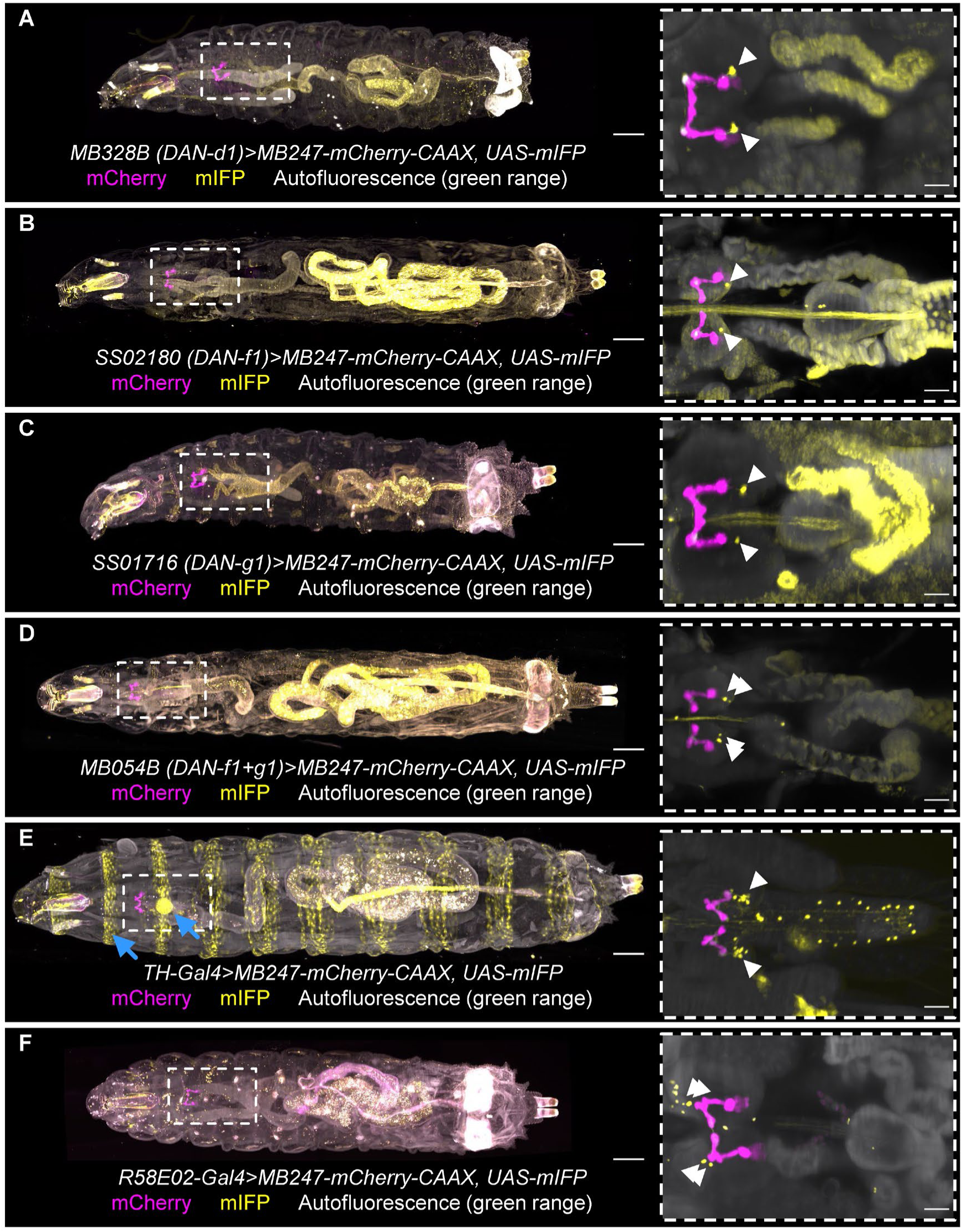
Whole-body expression patterns of DAN-driver strains. All driver strains were crossed to a *MB247-mCherry-CAAX, UAS-mIFP* double effector that will show a mCherry signal (magenta) in the mushroom body and a mIFP signal (yellow) controlled by the driver strain. In addition, autofluorescence in the green range is shown in white. The insert shows a volume-restricted magnification of the CNS region. For images of each separate channel, as well as for genetic controls, see supplementary material S1. (A) *MB328B*, (B) *SS02180*, (C) *SS01716* and (D) *MB054B* each drive expression in one to three cells per hemisphere (arrowheads) innervating the mushroom body as described previously^15,25^. No additional cells were found in the body. (E) *TH-Gal4* drives additionally to the expression in the CNS (arrowheads indicate the DL1 cluster) also outside of the brain, most prominently in the melanin-producing cells at the segment barriers and the proventriculus (blue arrows). (F) *R58E02-Gal4* drives expression in three cells per hemisphere (arrowheads) innervating the medial lobe of the mushroom body, as previously described^27,47^. Scale bars represent 250 µm for the whole body and 50 µm for the inserts.

### Some individual DANs establish and retrieve associative memories

In larvae, for many kinds of aversive US learned behaviour is only observed if the US is present at the moment of memory retrieval^32–36^. For example, we trained groups of larvae with an odour as the conditioned stimulus (CS) and quinine as US and observed an aversive associative memory when we tested the animals in presence of quinine, but not in its absence (Fig. 2A). Learned aversive behaviour therefore can be viewed as an escape from the US, guided by the respective memory – an escape that is pointless in absence of anything to escape from^37–40^.

**Figure 2:**
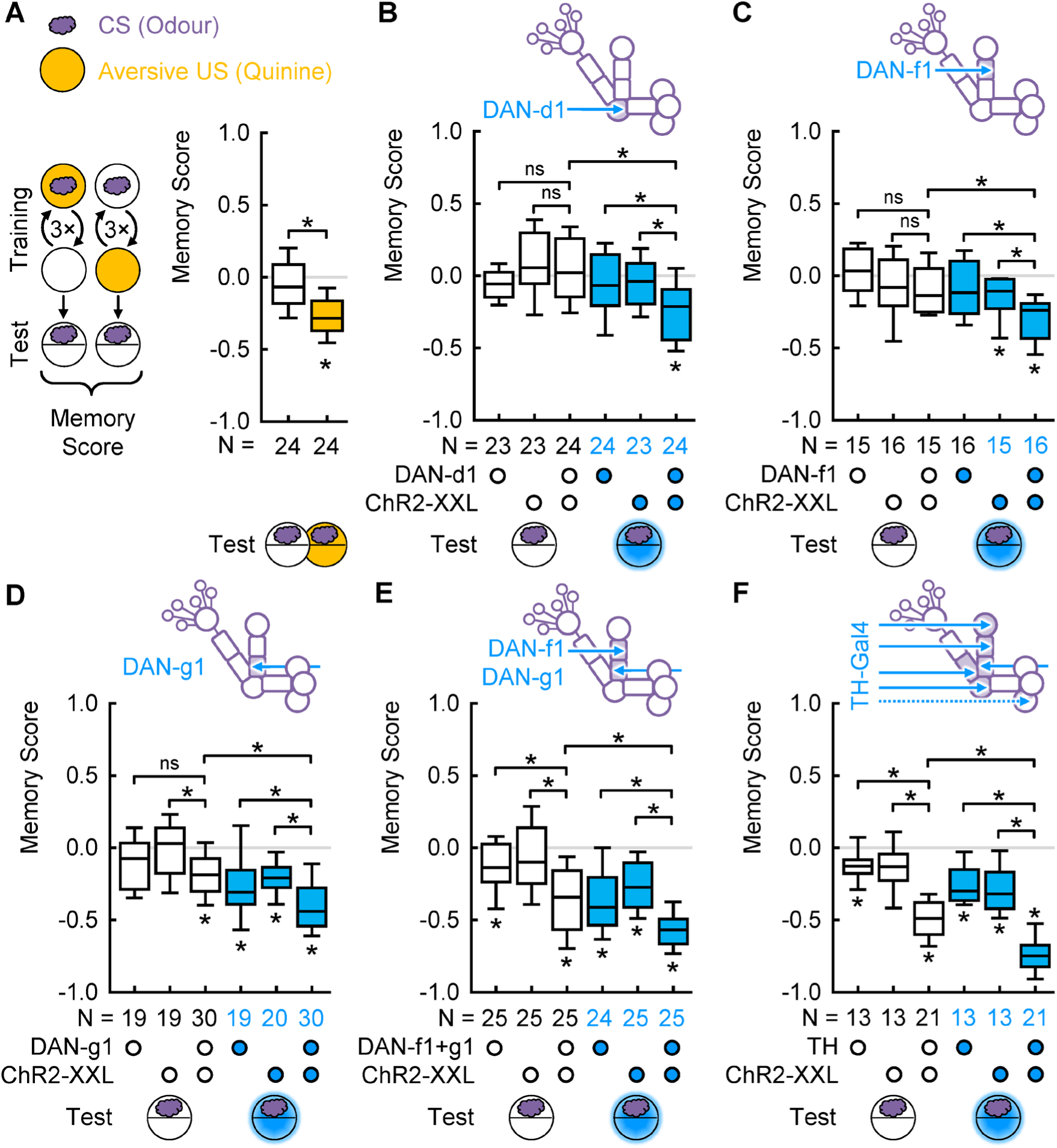
Individual DANs are sufficient to establish and retrieve aversive memory. (A) After training wild-type larvae with quinine as US, associative memory is observed only in presence of the US but not in its absence. (B) When animals were trained with DAN-d1 activation as US and tested in darkness, memory was retrieved in neither genotype. When tested in presence of blue light, negative memory scores only in the experimental genotype indicated aversive associative memory. For independent repetitions, see Figure supplement 2A-D. (C) As in (B), but with DAN-f1 activation. For an independent repetition, see Figure supplement 2E. (D) In the case of DAN-g1, aversive memory scores were observed in the experimental genotype even in darkness. When tested in blue light, aversive memory scores were observed in all genotypes, but significantly stronger in the experimental genotype. (E) As in (D), but with a combination of DAN-f1 and DAN-g1. For independent repetitions, see Figure supplement 2F-G. (F) As in (D), but with the broad *TH-Gal4* pattern. * and ns above brackets indicate pairwise significance or non-significance, respectively (MW), * below boxes indicate memory scores significantly different from zero (WSR). For the underlying source data and the results of the statistical tests, see Supplementary Data S1.

We then asked for the roles of the DL1-DANs for both establishing and retrieving associative memories by employing the same learning experiment, but instead of quinine we used optogenetic DAN activation by blue light as a virtual US. We also tested the innate odour preference of all genotypes, both in darkness and in blue light (Fig. 2 – Figure supplement 1) and observed no evidence that the innate odour preference was modified by activating the respective DANs.

After training with DAN-d1 as virtual US, zero memory scores were observed when the animals were tested in darkness. When tested in blue light, however, the animals of the experimental genotype, but not the controls, showed aversive associative memory (Fig. 2B, Fig. 2 – Figure supplement 2A-B). Interestingly, a recent study observed no significant memory with DAN-d1 as virtual US in a similar kind of experiment^25^. We wondered whether this difference could be attributed to the strength of DAN activation in different experimental setups. We therefore repeated the experiment with two different light intensities and observed an aversive memory only when using the low intensity that we use in all other learning experiments in this study, but not with a 10-fold higher intensity (Fig. 2 – Figure supplement 2C). Moreover, when raising the larvae in food supplemented with all-trans-retinal which had been shown to increase the effectiveness of our effector ChR2-XXL^41^, no aversive memory could be detected (Fig. 2 – Figure supplement 2D). Thus, it seems that a too strong activation of DAN-d1 prevents the establishment and/or retrieval of an aversive memory.

For DAN-f1, we likewise found an aversive memory that was retrieved only when DAN-f1 was activated also during the test (Fig. 2C, Fig. 2 – Figure supplement 2E). Our results indicate that DAN-d1 and DAN-f1 each are sufficient to establish an aversive memory which requires the respective DAN to be activated for retrieval.

A previous study used the same driver strains crossed to *UAS-CsChrimson* as the optogenetic effector in a similar learning experiment and observed learned behaviour also in darkness^15^. Using the same effector and the same intensity of red-light stimulation we were able to replicate these results (Fig. 2 – Figure supplement 3). Thus, the memories induced by either effector differed in the conditions of how they could be retrieved. These results call for caution against using different optogenetic effectors interchangeably.

### Some DANs establish memories that can be retrieved independently of the test conditions

Activation of DAN-g1, too, established an aversive memory (Fig. 2D). However, in contrast to DAN-d1 and DAN-f1, we found aversive associative memory scores in the experimental genotype independently of the testing condition (Fig. 2D).

Next, we employed two driver strains that each covers a combination of the DANs and activated them using ChR2-XXL. One of them covers DAN-f1 and DAN-g1, the other one, *TH-Gal4*, covers all three of them, plus many additional cells (Fig. 1E). In both cases, we observed associative memories in the experimental genotype both in absence and in presence of the US during the test (Fig. 2E-F, Fig. 2 – Figure supplement 2F-G).

Notably, in the experiments with these driver strains, we also observed significantly aversive memory scores in the genetic controls when tested in light (Fig. 2D-F). Given that this result was found both in driver and effector controls, it was most likely caused by the light itself. Light is known as an aversive US in larvae^42^ and has been seen to affect the results of optogenetic experiments before^24,43^. It is likely that the light naturally activated some aversive DANs through the visual system in addition to those we activated optogenetically in each experiment. Why this aversive effect of the light remained a non-significant trend in the previous experiments (Fig. 2B-C) is not known. In any case, this observation suggests that any effect of optogenetically activating DANs occurred on top of the light-induced effects. Therefore, we base our conclusions on the comparisons between the genotypes tested under the same light conditions and understand any effect of optogenetically activating a DAN as being additional to potential light-induced DAN activations.

Taken together, activating DAN-g1, alone or with additional DANs, established aversive memories that were retrieved independent of whether the respective DANs were activated during the test.

### Individual DANs are sufficient for establishing safety, but not punishment memories

Animals can associate stimuli not only with the presence of a threat, but also with its absence^44–45^. Such safety memory has been found in *D. melanogaster* but received little attention by the community until recently (larvae^40,46–48^; adults^49–50^). That means a US, such as bitter taste, can establish two different memories: punishment memory after paired training and safety memory after unpaired training^37,46^ (Fig. 3A). Critically, our results so far demonstrate that each of the individual DANs was sufficient to establish an associative memory – but not, whether this was a punishment memory, a safety memory, or both. Therefore, we next focussed on the animals’ odour preference after paired and unpaired training, respectively, compared to the odour preference of the pooled genetic controls as a baseline. A punishment memory would reveal itself by decreased odour preferences after paired training, a safety memory by increased preferences after unpaired training, compared to the baseline. Due to the rather small effect sizes and high variability of odour preferences, we pooled the data of all independent replications of the experiments (Fig. 2 including Figure supplements 2 and 3) as well as the genetic controls of each experiment for a post-hoc analysis of the odour preference.

**Figure 3:**
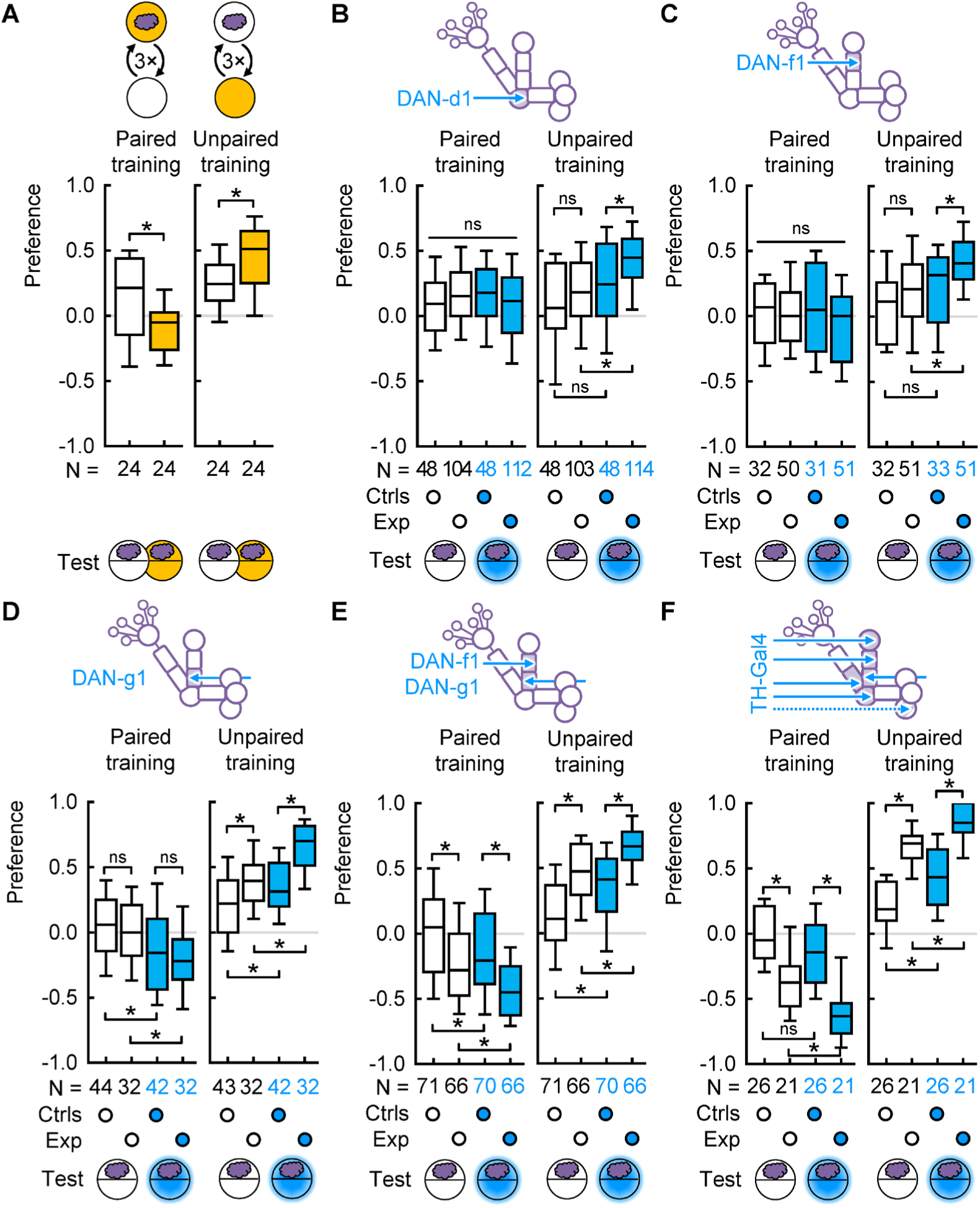
Individual DANs are sufficient to establish safety memory. (A) With quinine as US, odour preferences after paired training were more negative in presence than in absence of the US (left, indicating punishment memory), and after unpaired training were more positive (right, indicating safety memory). (B) Left: after paired training, the animals of the pooled genetic controls and the experimental genotype (*DAN-d1>ChR2-XXL*) showed the same odour preference, both when tested in darkness or light. Right: after unpaired training, the experimental genotype showed a more positive odour preference than the genetic controls only when tested in light, indicating safety memory. (C) As in (B), but with DAN-f1 activation. (D) Left: for DAN-g1, after paired training the animals of both the controls and the experimental genotype showed lower odour preferences when tested in presence than in absence of light. Right: after unpaired training, animals of the experimental genotype showed higher odour preferences than the controls, irrespective of the testing condition (right). (E) For a combination of DAN-f1 and DAN-g1, the animals of the experimental genotype showed more negative odour preferences than the controls after paired training (left, indicating punishment memory) and more positive preferences after unpaired training (right, indicating safety memory), irrespective of the testing condition. (F) For the *TH-Gal4*-DANs, similar results as in (E) were observed. ns above a line indicates non-significance across the whole data set (KW), * and ns above brackets indicate pairwise significance or non-significance, respectively (MW). For the underlying source data and the results of the statistical tests, see Supplementary Data S1.

For the case of DAN-d1, after paired training there was no difference in odour preference between the genetic controls and the experimental genotype, nor between the two testing conditions (Fig. 3B, left). After unpaired training, in contrast, the larvae of the experimental genotype exerted increased odour preferences when tested under blue light compared to all other groups (Fig. 3B, right). This suggests that the activation of DAN-d1 established no punishment memory, but a safety memory. For DAN-f1, we found analogous results (Fig. 3C). In the case of DAN-g1, after paired training we found more negative odour preferences when tested in presence than in absence of light, but no difference between the controls and the experimental genotype (Fig. 3D, left). This suggests a potential punishment memory by the light, but not by the activation of DAN-g1. After unpaired training, the animals of the experimental genotype showed higher odour preferences than the controls, both in darkness and light (Fig. 3D, right).

Taken together, we found that all three DANs induced only a safety memory after unpaired training, but no punishment memory after paired training (confirming similar results using CsChrimson^15^). The safety memory established by DAN-d1 and DAN-f1 were retrieved only if the respective neuron was activated during the test, whereas the safety memory established by DAN-g1 was, at least partially, also retrieved without neuronal activation.

### Combinations of DANs are sufficient for establishing both safety and punishment memories

We next wondered whether the activation of a combination of DANs would be sufficient to induce a punishment memory. For the combination of DAN-f1 and DAN-g1, after paired training we indeed found more negative odour preferences in the experimental group than in the genetic controls, irrespective of the testing condition (Fig. 3E, left). After unpaired training, we found higher odour preferences in the experimental genotype than in the genetic controls (Fig. 3E, right). With *TH-Gal4* we observed analogous results (Fig. 3F).

Taken together, activating DAN-f1 and DAN-g1 together, or activating most DANs in the brain, did establish both a punishment and a safety memory. Both kinds of memory were retrieved irrespective of whether the DANs were activated in the test or not.

### Activating the DANs covered by TH-Gal4 modulates the animals’ bending and velocity

Next, we investigated which effect activating the DL1-DANs would have on locomotion.

Using the novel IMBA (Individual Maggot Behaviour Analysis) software for behavioural analyses, we recently had found strong locomotor effects upon optogenetic activation of the DANs covered by *TH-Gal4*^51^. Within the current study, we repeated this experiment in total five times, in two different laboratories and with different parameters (Fig. 4 – Figure supplement 1A). This gave us the opportunity to probe the behavioural effects upon DAN activation for their robustness across experimental repetitions. Groups of around 8 experimentally naïve larvae of either the experimental genotype or the driver or effector controls were allowed to crawl freely on an agar-filled Petri dish in darkness. After 30 s, blue light was activated for 30 s, followed by darkness. We video-recorded the animals’ behaviour and analysed it afterwards with the IMBA software (Fig. 4A).

**Figure 4:**
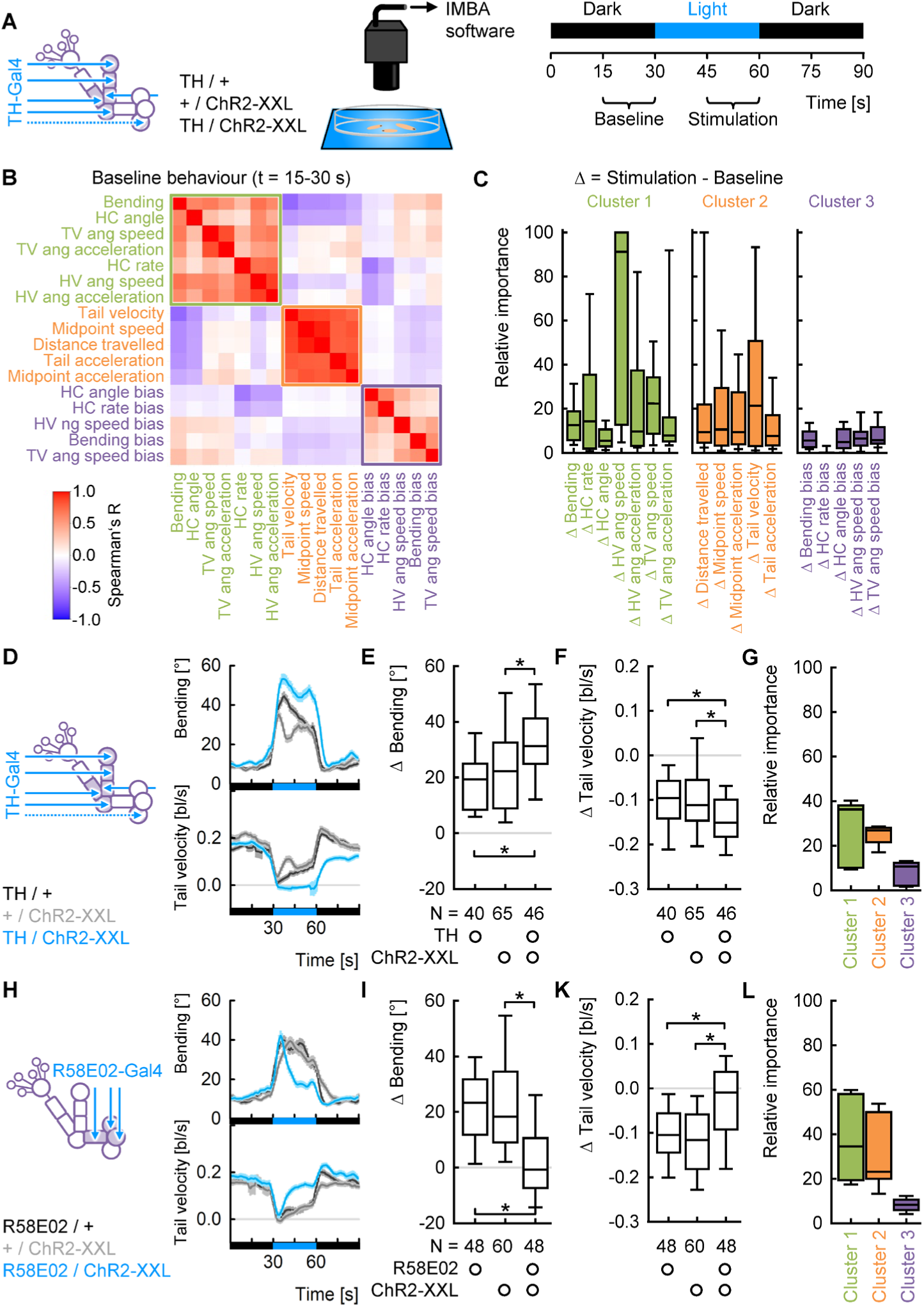
Two sets of DANs induce opposite effects on locomotion. (A) Behaviour of cohorts of experimentally naïve larvae was recorded. After 30 s of darkness, blue light was activated for 30 s. The second halves of the dark and light periods are henceforth referred to as “baseline” and “stimulation”, respectively. (B) Based on data from 5 experiments involving *TH>ChR2-XXL* and genetic controls within this study, the cross-correlation of 17 behavioural attributes during the baseline period revealed three clusters of inter-correlated attributes. Displayed is the Spearman’s correlation coefficient R for each pair of attributes. Values of 1 (red) indicate perfect positive, values of −1 (blue) perfect negative correlations. (C) A random forest algorithm was trained to classify individuals of the experimental genotype and either genetic control, using their change in behaviour between the baseline and stimulation. Displayed is the relative importance of each behavioural attribute, scaled from 0 (least important) to 100 (most important). (D) Average bending and tail velocity over time. The black and blue bars at the bottom indicate the light regimen. (E) The change in bending upon light stimulation. (F) As in (E), but for the tail velocity. (G) The average relative importance of each attribute cluster, as provided by the random forest. (H-L) As in (D-G), but using *R58E02-Gal4* as driver. * and ns above brackets indicate pairwise significance or non-significance, respectively (MW). For the underlying source data and the results of the statistical tests, see Supplementary Data S1.

The IMBA software provides a rich variety of behavioural attributes for individual animals. We selected 17 attributes (for a list and descriptions, see table 1) and first asked how these attributes relate to each other. Towards this end, we performed a Spearman correlation analysis of all 17 attributes based on the animals’ baseline behaviour in darkness. We found three clusters of inter-correlated attributes (Fig. 4B). Cluster 1 consists of attributes measuring different aspects of directional changes, e.g. the animals’ bending, or the rate of lateral head movements, the so-called head casts (HC). All attributes of cluster 2 relate to the speed of the larvae’s peristaltic movement. Finally, cluster 3 consists of attributes dealing with the directional bias of the animals’ behaviour. For example, the bending bias measures to which extent an animal was bent more towards one side than another.

**Table 1:** A list and description of all behavioural attributes used for locomotion analysis in this study.

| Name | Unit | Description | Cluster |
| --- | --- | --- | --- |
| Bending | ° | The bending angle (BA) is the angle between the larva's tail vector and a vector from the middle of the larva to its head. Positive values indicate the larva bending to the left, negative values bending to the right.<br>"Bending" in this study refers to the absolute, i.e. direction-independent BA. | 1 |
| Bending bias | | The absolute average of all BA values of a larva, divided by the average of all absolute BA values: $\text{abs}(\overline{BA})/(\overline{\text{abs}(BA)})$. The Bending bias is constrained between 0 and 1, with 0 indicating a larva was bending equally to left and right, and 1 indicating a larva was only bending into one direction (no matter which direction). | 3 |
| HV angular speed | °<br>—<br>s | The head vector angular speed (HVAS) is the angular speed of the head vector (HV) through the head of the larva. Positive values indicate movement of the head to the right, negative values to the left.<br>"HV angular speed" in this study refers to the absolute, i.e. direction-independent HVAS. | 1 |
| HV angular speed bias |  | Calculated analogous to the Bending bias. Constrained between 0 and 1, with 0 indicating a larva was moving its head equally fast to left and right, and 1 indicating a larva was moving its head only into one direction. | 3 |
| HV angular acceleration | °<br>—<br>s<sup>2</sup> | The absolute angular acceleration of the head vector (HV). | 1 |
| TV angular speed | °<br>—<br>s | The tail vector angular speed (TVAS) is the angular speed of the tail vector (TV) through the rear half of the larva. Positive values indicate movement to the right, negative values to the left.<br>"TV angular speed" in this study refers to the absolute, i.e. direction-independent TVAS. | 1 |
| TV angular speed bias |  | Calculated analogous to the Bending bias. Constrained between 0 and 1, with 0 indicating a larva was moving its tail equally fast to left and right, and 1 indicating a larva was moving its tail only into one direction. | 3 |
| TV angular acceleration | °<br>—<br>s<sup>2</sup> | The absolute angular acceleration of the tail vector (TV). | 1 |
| HC rate | $\frac{HC}{s}$ | The number of head casts (HC) per second. HCs were defined as events of a HVAS larger than 35 °/s or smaller than -35 °/s, respectively. | 1 |
| HC rate bias |  | The HC rate of HCs to the left minus the HC rate of HCs to the right, divided by the total HC rate. Constrained between 0 and 1, with 0 indicating a larva had equal HC rates to left and right, and 1 indicating a larva was making HCs in only one direction. | 3 |
| HC angle | ° | The head cast angle (HCA) indicates how much the orientation of the animal changes upon each HC. It is defined as the difference between the bending angle at the end and the beginning of the HC. Positive values indicate that the larva changes its orientation to the left, negative values that it changes its orientation to the right.<br>"HC angle" in this study refers to the absolute, i.e. direction-independent HCA. | 1 |
| HC angle bias |  | Calculated analogous to the Bending bias. Constrained between 0 and 1, with 0 indicating a larva was making equally large HCs to left and right, and 1 indicating a larva was making HCs in only into one direction. | 3 |
| Distance travelled | bl | The distance of the complete path a larva travels during the observation period (measured in body lengths [bl]). | 2 |
| Midpoint speed | $\frac{bl}{s}$ | The speed of the larva's midpoint, irrespective of direction. | 2 |
| Midpoint acceleration | $\frac{bl}{s^2}$ | The absolute acceleration of the larva's midpoint. | 2 |
| Tail velocity | $\frac{bl}{s}$ | The velocity of the larva's tail point in a forward direction, that is in the direction of the tail vector. | 2 |
| Tail acceleration | $\frac{bl}{s^2}$ | The absolute acceleration of the larva's tail in the direction of the tail vector. | 2 |

Then, we wondered how much each attribute would change upon light stimulation. Towards this end, we calculated a Δ-value of each attribute as an individual’s behaviour during the second half of the light stimulation minus its baseline behaviour during darkness (Fig. 4A). We used only the second half of the stimulation to exclude any potential startle-like response to the sudden light. Then, we used a random forest classification algorithm to classify individual larvae between the genotypes based on the 17 Δ-attributes. Crucially, the random forest provides the relative importance of each attribute for the classification. In other words, it can be used to obtain an unbiased measure of how relevant each attribute is for discriminating the behaviour of different genotypes^51^. We found consistently the attributes of cluster 1 and 2 to be much more important than those of cluster 3 (Fig. 4C). Within each cluster, most attributes were on average similarly important, with strong variability both within and across data sets (Fig. 4 – Figure supplement 1).

We concluded that there does not appear to be a single “most important” behavioural attribute that accounts for the differences between the experimental genotype and the genetic controls. Rather, the differences are best described by clusters 1 and 2, representing the amount of bending and directional changes and the peristaltic speed, respectively. Within each cluster, the attributes were strongly correlated (Fig. 4B) and varied in importance (Fig. 4 – Figure supplement 1). Therefore, we decided to present the relative importance of each attribute cluster along with each experiment. Additionally, we present one representative attribute of each cluster 1 and cluster 2. We picked bending and tail velocity as intuitive behavioural descriptors that have been used previously^51^.

### Activating different subsets of DANs modulates locomotion in opposite directions

In accordance with our previous results^51^, we found that the control genotypes increased their bending and decreased their tail velocity upon light stimulation (Fig. 4D, Supplementary Movie 2). Such responses to blue light have been reported before^52^ and were consistently seen throughout this study (Fig. 4 – Figure supplement 2). Both types of behavioural change were significantly enhanced in the experimental genotype in which the *TH-Gal4*-DANs were activated^51^ (Fig. 4D-F). More specifically, when the DANs were activated, the larvae kept high bending and low velocity throughout the 30 seconds of light stimulation whereas the controls gradually reverted towards their normal behaviour (Fig. 4D). These results were confirmed by a repetition of the experiment using larger Petri dishes and multiple light stimulations (Fig. 4 – Figure supplement 3). For the random forest, the attributes from cluster 1 and 2 were equally important (Fig. 4G).

As *TH-Gal4* covers all DANs except those of the pPAM cluster, we next asked whether activation of the pPAM DANs would have a similar effect on locomotion. Towards this end, we used *R58E02-Gal4* which covers 3 out of 4 pPAM DANs (DAN-h1, DAN-i1 and DAN-j1)^27^. To our surprise, activation of these DANs decreased bending and increased tail velocity compared to the controls (Fig. 4H-K, Supplementary Movie 3). More precisely, after an initial reaction to switching on the light, when the pPAM DANs were activated, the larvae reverted to their normal behaviour much quicker than the controls (Fig. 4H). Again, the attributes of cluster 1 and 2 were similarly important for the random forest classification (Fig. 4L).

Thus, the activation of two non-overlapping sets of DANs had opposite effects on behaviour: activating the *TH-Gal4*-DANs (including the aversive US-signalling DANs) increased bending and decreased velocity, whereas activating the *R58E02-Gal4*-DANs (which were shown to signal appetitive US in larvae^27,47,53^ and adults^54–55^) decreased bending and increased velocity.

### DANs modulate locomotion independently of the light responses

In the previous experiments, animals of all genotypes displayed a very strong response to the light stimulation. We therefore wondered whether the observed effects upon DAN activation were due to a modulation of this light response, or an independent modulation of the animals’ locomotion.

We compared the animals’ behavioural changes in three sets of experiments, activating the DANs with strong blue light, weak blue light, or red light, respectively (Fig. 5). When using the about 10-fold lower light intensity, we nevertheless observed rather strong light responses in the controls (Fig. 4 – Figure supplement 2), and robust differences in behaviour upon activation of either set of DANs (Fig. 5 – Figure supplement 1).

**Figure 5:**
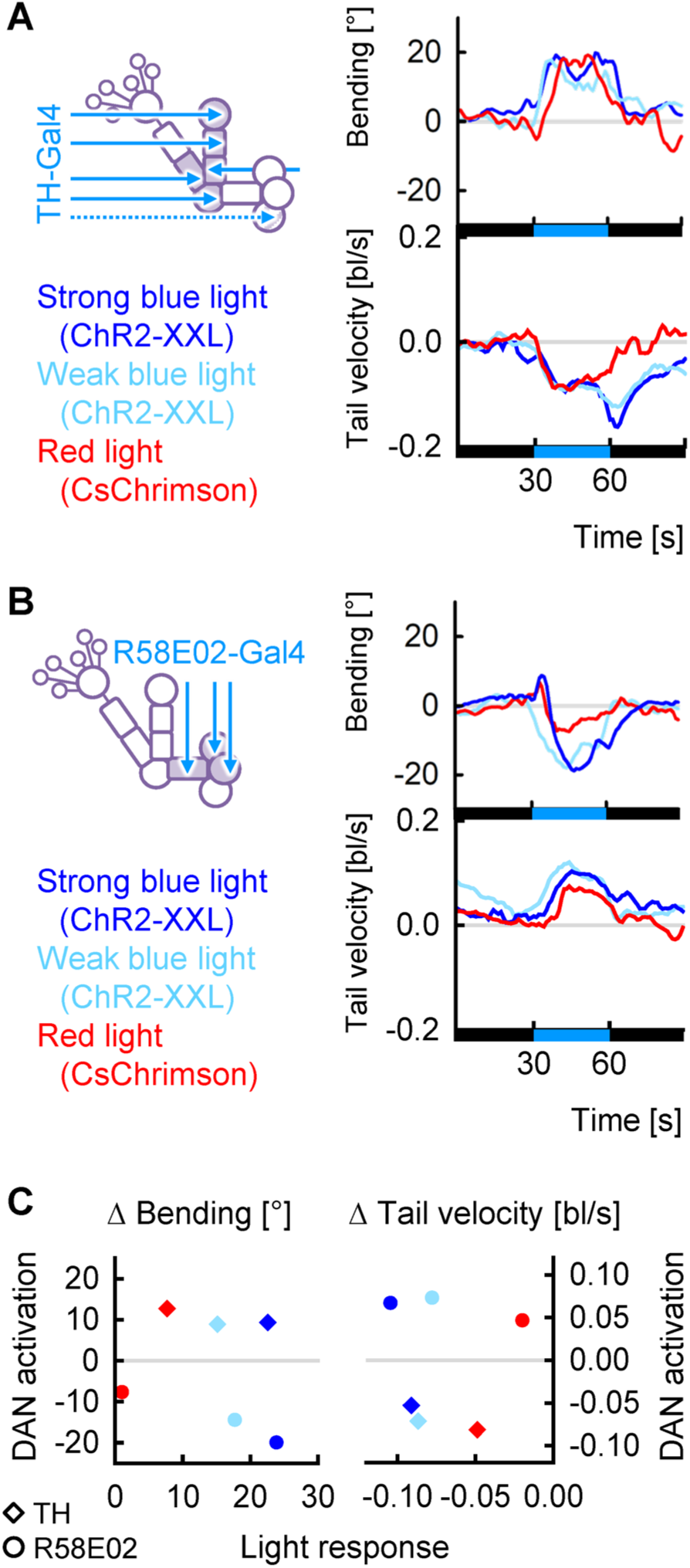
DAN activations modulate locomotion independently of light. (A) Average difference in bending and tail velocity of the experimental group from the control groups over time. Black and blue bars at the x-axis indicate the light regimen. Each curve represents the behaviour of the experimental group minus the pooled controls of a given experiment to indicate the effect of DAN activation, “purified” from the light responses. Compared are experiments using strong blue light (dark blue, Fig. 4D) and weak blue light (light blue, Figure supplement 1A) with ChR2-XXL as effector, and red light (red, Figure supplement 2A) with CsChrimson. (B) As in (A), but for the experiments shown in Figures 4H, Figure supplement 1E and 2E. (C) For each of the 6 experiments, the average Δ bending (left) or Δ tail velocity (right) of the pooled control groups was taken as the effect size of the light response (x-axis). The Δ bending or Δ tail velocity of the experimental group minus those of the pooled control groups was taken as the effect size of DAN activation (y-axis). Note that the left and right graphs use different axes and scaling. The colours match those of (A-B). All experiments using the same driver resulted in similar effect sizes of DAN activation, despite strong differences in effect sizes of the light response.

Red light reportedly triggers only very mild light responses^16,52^. We therefore next used the red-shifted effector CsChrimson^54^ and raised larvae either with or without all-trans-retinal which is necessary for the Chrimson channels to function^56^. When the larvae were raised with retinal, we observed similar effects as when using ChR2-XXL (Fig. 5 – Figure supplement 2). The control without retinal, in contrast, displayed only very weak responses to the red light (Fig. 5 – Figure supplement 3).

In order to compare the effects of DAN activation between the three experiments, we defined the behaviour of the pooled controls in each experiment as zero and displayed the difference of each experimental group from their respective controls. The behavioural effects of the DAN activations were strikingly similar in the three conditions, both in magnitude and time course (Fig. 5A-B). The only obvious difference was an extended reduction of the tail velocity beyond the activation of the *TH-Gal4*-DANs that could be seen only when using ChR2-XXL (Fig. 5A). This “after-effect” may be due to the reportedly slow closing times of the ChR2-XXL channel^41^.

We wanted to further quantify the relation between the light responses elicited in each experiment and the behavioural effects of the DAN activation. Therefore, we used the mean behavioural changes upon light stimulation of the pooled controls of each experiment as an indicator for the effect size of the light response, and the difference in behavioural changes between the experimental groups and the respective controls as an indicator for the effect size of the DAN activation. We found that the effect sizes of the DAN activation were all relatively similar and completely unrelated to the effect sizes of the light responses (Fig. 5C).

In summary, we concluded that the observed modulations of bending and tail velocity upon DAN activation occurred independently of the animals’ responses to the light stimulation itself.

### Modulations of locomotion do not determine olfactory behaviour

The experiments shown in Fig. 5 – Figure supplement 1 used the same light intensity and genotypes as our previous olfactory experiments with *TH-Gal4*-DANs (Fig. 2F, Fig. 2 – Figure supplement 1F, Fig. 3F). Considering the results of both kinds of experiments, two questions arise: how can larvae approach or avoid an odour while the *TH-Gal4*-DANs are activated (as seen e.g. in Fig. 3F) if activating *TH-Gal4*-DANs completely stops forward locomotion (as seen in Fig. 5 – Figure supplement 1)? And is it possible that the modulations upon DAN activation directly determine the odour preference? To address these questions, we analysed video recordings of the test phases of the experiments shown in Fig. 2 – Figure supplement 1F and Fig. 3F. During innate olfactory preference (Fig. 6A-C), we found increased bending and decreased tail velocity when the experimental genotype was tested in light rather than in darkness. Both effects were weaker than those during the locomotion experiments and declined further over time (Fig. 6B-C). In other words, in this experiment the animals’ locomotion was only moderately altered, in a way that still allowed them to reach their goal (Fig. 6A). A possible reason for the differing strengths of behavioural responses may be the different handling of the animals: in the olfactory experiments, the Petri dish was loaded with the larvae in dimmed room light and placed in the experimental setup, followed by an immediate start of the light stimulation and video recording. In the locomotion experiments, the larvae were kept in darkness for at least 40 s before the light stimulation started. The sudden light stimulus after a period of darkness may trigger stronger modulations of locomotion.

**Figure 6:**
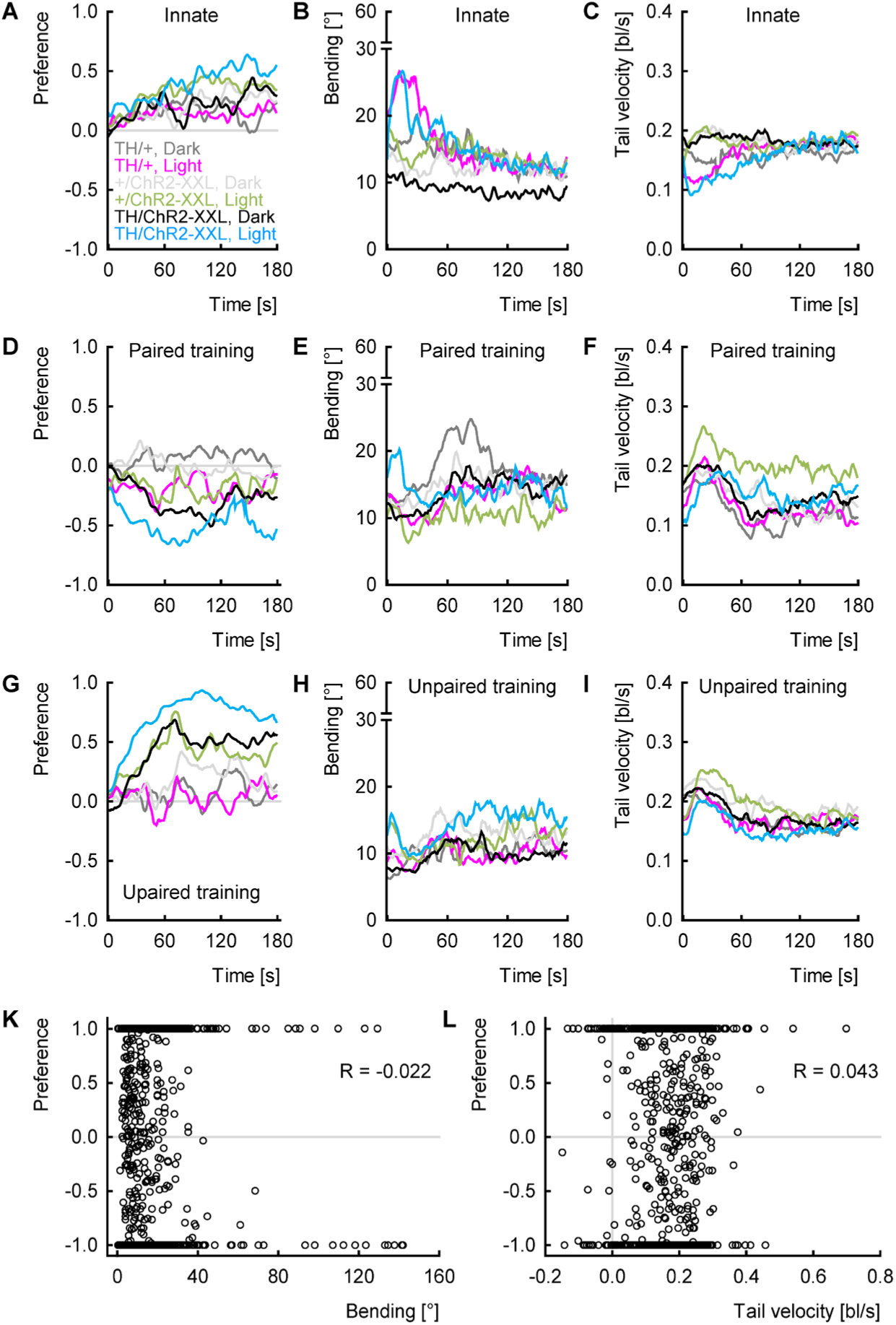
DAN activation during innate odour preference and learning experiments causes weak locomotion modulations. (A) Larvae were video-recorded and their behaviour analysed during the experiment displayed in (A-C) Fig. 2 – Figure supplement 1F (innate odour preference), (D-F) Fig. 3F (paired training) and (G-I) Fig. 3F (unpaired training). (A) Larvae of all genotypes and conditions approached the odour over time, with the experimental genotype (*TH>ChR2-XXL*) tested in light (blue) having the strongest preference (see Fig. 2 – Figure supplement 1F). (B) The larvae of the experimental genotype had a higher average bending in blue light (blue) than in darkness (black). The driver control, too, had a higher bending in light (pink) than in darkness (dark grey) for the first third of the experiment. (C) The larvae of the experimental genotype showed a lower tail velocity in light than in darkness during the first half of the experiment. (D) After unpaired training, the larvae avoided the odour over time, with the experimental genotype showing the strongest aversion, especially when tested in light (see Fig. 3F, left). (E) Only in the first 10 s of the experiment, the larvae of the experimental genotype showed stronger bending in light than in darkness. (F) The larvae of all genotypes and conditions had a similar tail velocity, with a small peak in the first 20 s. (G) After paired training, the larvae approached the odour, with the larvae of the experimental genotype having the highest preference, especially when tested in light (see Fig. 3F, right). (H-I) As in (E-F), but after unpaired training. (K) No correlation between the bending of individual animals during the first 60 s of all three experiments and their odour preference in the following 60 s. (L) As in (K), but for the tail velocity. Displayed is Spearman’s correlations coefficient R as an indicator of the correlation. For the underlying source data and the results of the statistical tests, see Supplementary Data S1.

During the testing phase after paired training (Fig. 6D-F) and unpaired training (Fig. 6G-I), the locomotion effects were even weaker: the experimental genotype showed only a small peak in bending within the first approximately 10 s of the test (Fig. 6E,H) and was not different from the other genotypes in terms of forward velocity (Fig. 6F,I). It appears that the repeated exposure to both light and DAN activation during the training further reduced the modulations of locomotion.

Notably, although bending and velocity were very similar after paired and unpaired training in the experimental genotype, the animals’ odour preference was drastically different, with strong avoidance after paired training (Fig. 6D) and approach after unpaired training (Fig. 6G). This suggests that the preference is not directly dependent on locomotion modulations. Next, we addressed this question more directly. We asked whether the bending or tail velocity of each individual larvae within the first 60 s would be correlated with its average preference (that is, its position on the dish) during the following 60 s, which would be indicative of a causal relationship. However, no such correlation was found (Fig. 6K-L), suggesting that the types of locomotion modulations we study here do not have a direct impact on odour preference. This is consistent with previous studies, which identified the timing and direction of reorientation events with respect to the odour as the major determinants of olfactory preference^40,51,57–58^.

### Different subsets of DANs convey opposite acute valence when activated

Next, we asked how to interpret the behavioural effects of activating DANs. Stopping and turning around upon a sudden activation of *TH-Gal4*-DANs can be interpreted as a reaction to an aversive signal^16,52^ – on the other hand, slowing down and turning on the spot could also be a behavioural strategy to stay in a pleasant situation. Likewise, going straight and speeding up upon *R58E02-Gal4*-DAN activation can be interpreted as a “go on”-reaction to an appetitive signal or a strategy to escape. In both larvae and adult flies, the artificial activation of *TH-Gal4*-DANs and *R58E02-Gal4*-DANs induce aversive and appetitive memories, respectively^27,47,53,54–55,59–60^. However, studies in adults suggest that acute valence (approach/avoidance) and associative valence (appetitive/aversive US) are often not correlated^60–62^. Therefore, to test for the valence of activation of either set of DANs in larvae we performed choice experiments between a dark and an illuminated half of a Petri dish (Fig. 7A). Larvae of all genotypes avoided the illuminated half as expected^42,52^. Relative to the genetic controls, larvae with *TH-Gal4-*DANs activated showed more avoidance, and larvae with *R58E02-Gal4-*DANs showed more approach (Fig. 7B-C, Supplementary Movies 4-5). With CsChrimson as effector and red instead of blue light, larvae had a positive preference for having their *R58E02-Gal4-*DANs activated (Fig. 7 – Figure supplement 1A). We next analysed the animals’ behaviour near the dark-light-border. When activating *TH-Gal4*-DANs, the animals’ bending was increased and their velocity decreased on the illuminated half compared to the controls (Fig. 7D-F), similar to our previous experiments (Fig. 4). When activating *R58E02-Gal4*-DANs, the animals’ bending was decreased and their velocity increased compared to the controls everywhere on the dish (Fig. 7G-I, Fig. 7 – Figure supplement 1B-D).

**Figure 7:**
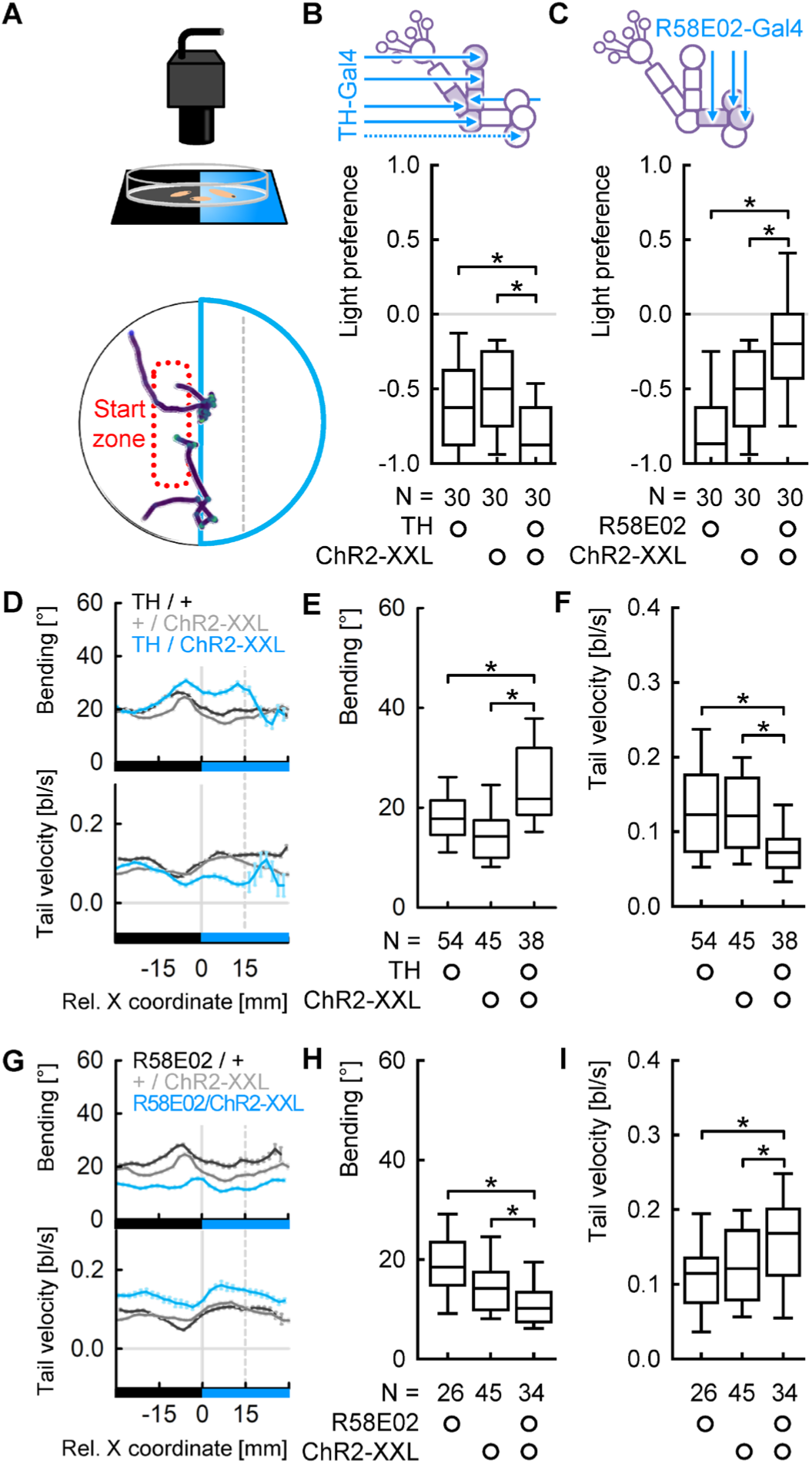
Different subsets of DANs convey opposite acute valence when activated. (A) Larvae were given the choice between a dark and an illuminated (weak blue light) half of a Petri dish and their behaviour was recorded for three minutes. Animals were placed into the dark half at the dish. Displayed are 2 sample tracks for animals of TH>ChR2-XXL. (B) The light preference after 3 minutes. All genotypes showed light avoidance which was enhanced in the experimental genotype. (C) As in (B), but for R58E02-Gal4 as driver. The experimental genotype showed less light avoidance than the controls. (D) Average bending and tail velocity of TH>ChR2-XXL and genetic controls over the X coordinate of the Petri dish, with 0 indicating the midline. The black and blue bars at the bottom indicate which half was illuminated. (E) The average bending on the illuminated side of the dish. We limited the analysis to an area up to 15 mm from the midline (indicated as stippled line in A) to capture the behaviour in the choice zone. (F) As in (E), but for the tail velocity. (G-I) As in (D-F), but for R58E02-Gal4 as driver. * and ns above brackets indicate pairwise significance or non-significance, respectively (MW). For the underlying source data and the results of the statistical tests, see Supplementary Data S1.

Taken together, we concluded that activating *TH-Gal4*-DANs conveys an acute aversive signal and activating *R58E02-Gal4*-DANs conveys an acute appetitive signal. The animals’ reaction to the sudden activation of these DANs therefore should be interpreted as aversive and appetitive behaviour, respectively.

### The modulations of bending and velocity are dopamine-dependent

Several DANs have been found to have additional neurotransmitters^19,63^. Therefore, we wondered whether the observed effects on locomotion were indeed dependent on dopamine signalling. Towards this end, we combined optogenetic activation by ChR2-XXL with TH-RNAi which knocks down the expression of tyrosine hydroxylase, the rate limiting enzyme of dopamine synthesis. Thus, when both effectors are expressed in a set of DANs, these DANs can get activated by light but have a reduced dopamine synthesis. We found that animals with the *ple-dsRNA* (TH-RNAi) construct have a higher baseline bending before stimulation than without, irrespective of the presence of *TH-Gal4* (Fig. 8A-B). In addition, the baseline tail velocity was increased in the animals expressing both effectors in the *TH-Gal4-*DANs (Fig. 8A,C). These observations suggest that the TH knockdown was effective in modulating behaviour. Upon stimulation, we found an increased bending in both genotypes that expressed ChR2-XXL compared to the genetic controls. This increase was slightly smaller in animals with the TH-RNAi (Fig. 8D, 3^rd^ and 5^th^ group from left). The change in tail velocity upon DAN activation was unaffected by the TH-RNAi (Fig. 8E, 3^rd^ and 5^th^ group from left). The attributes of cluster 1 and 2 were both important to discriminate between the genotypes, suggesting that the TH-RNAi did also affect attributes related to forward locomotion to some extent (Fig. 8F-G). Nevertheless, the effects of the TH-RNAi were overall rather weak. There are several possible reasons for this lack of effects, including a too weak knock-down of tyrosine hydroxylase by the *ple-dsRNA* construct used, or a developmental adaptation to the reduced dopamine levels. Therefore, we next sought to inhibit tyrosine hydroxylase by an independent and acute technique.

**Figure 8:**
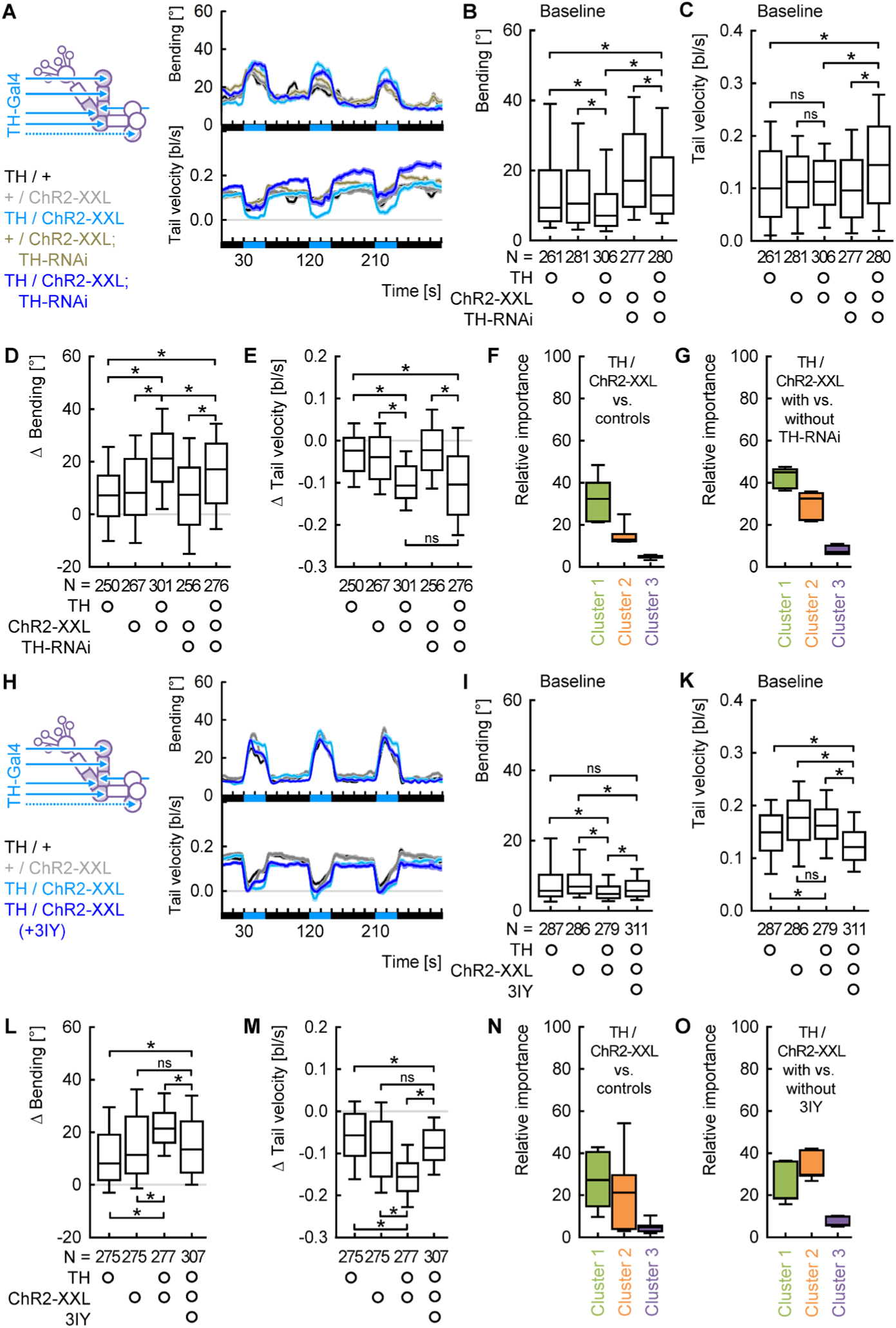
The effects on locomotion upon DAN activation are dopamine-dependent. (A) Average difference in bending and tail velocity of the experimental groups from the control groups over time. Black and blue bars at the x-axis indicate the light regimen. (B) The animals’ average bending during the baseline period (15 to 30 s) in darkness. Both genotypes including the *ple-dsRNA* (TH-RNAi) construct displayed increased bending, suggesting a genetic background effect. (C) As for (B), but for the tail velocity. Animals expressing *ple-dsRNA* had an increased velocity compared to all other genotypes. (D) The change in bending upon the first light stimulation. This change was significantly increased in the experimental genotype and slightly reduced when in addition expressing *ple-dsRNA*. (E) As in (D), but for the tail velocity. (F) The average relative importance of all behavioural attributes within each cluster, as provided by the random forest algorithm when comparing *TH>ChR2-XXL* with the genetic controls. (G) The average relative importance of each attribute cluster for a comparison between animals with and without *TH-RNAi*. (H-O) As for (A-G) but feeding the TH inhibitor 3IY instead of inducing *TH-RNAi*. The changes in both bending and tail velocity upon light stimulation for animals fed the TH inhibitor 3IY were statistically different from animals without 3IY feeding and not statistically different from the genetic controls. * and ns above brackets indicate pairwise significance or non-significance, respectively (MW). For the underlying source data and the results of the statistical tests, see Supplementary Data S1.

Larvae were fed the tyrosine hydroxylase inhibitor 3IY for four hours before the experiment, following established protocols^64–65^. 3IY-fed larvae showed different baseline behaviour compared to animals without 3IY, but overall, the differences were less pronounced than in the case of TH-RNAi (Fig. 8H-K). Interestingly, the baseline tail velocity was decreased rather than increased compared to animals without 3IY feeding (Fig. 8K). This means inhibiting the activity of tyrosine hydroxylase acutely or during the whole development had opposite effects on the baseline tail velocity. Upon DAN activation, 3IY-fed animals increased their bending less, and reduced their tail velocity less, than animals without 3IY, in both cases to levels comparable to the genetic controls (Fig. 8L-O). These effects, both regarding the changes in bending and tail velocity as well as the shifted baseline velocity, were replicated in two independent experiments (Fig. 8 – Figure supplement 1).

Taken together, we concluded that the locomotor effects upon activation of the *TH-Gal4*-DANs were largely dopamine-dependent, although we note that this conclusion is based mostly on inhibiting the tyrosine hydroxylase acutely with 3IY.

### Individual DANs are sufficient to increase bending and decrease velocity

Given the broad expression pattern of *TH-Gal4*, we wondered which cells were responsible for the observed modulations of bending and velocity. Therefore, we probed the DANs of the DL1 cluster individually with experiments that included two cycles of light stimulation (Fig. 9 – Figure supplement 1). Overall, we found similar differences between the experimental genotype and the controls upon both stimulations, but more enhanced during the second stimulation.

**Figure 9:**
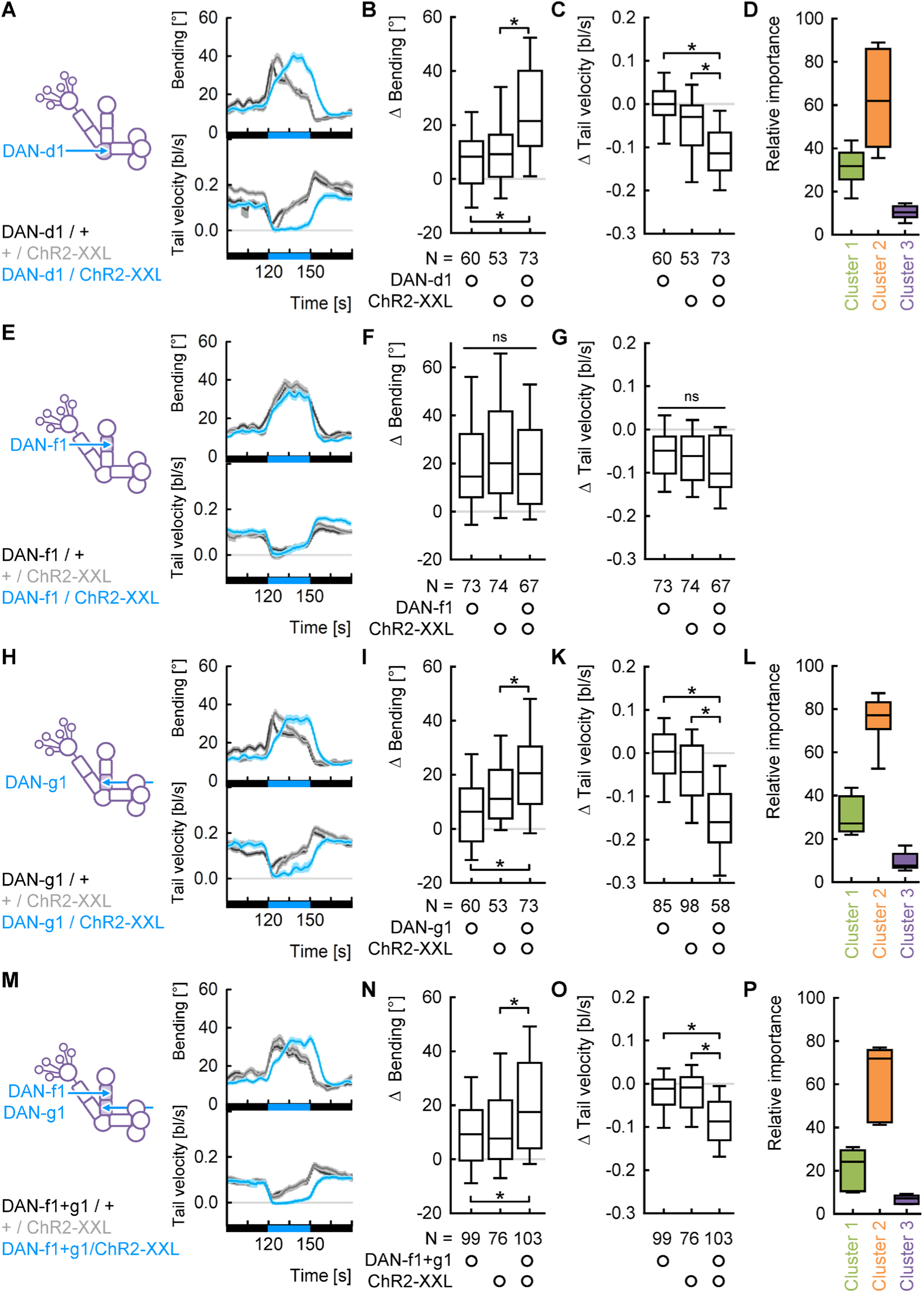
Individual DANs are sufficient to modulate bending and tail velocity. (A) Average bending and tail velocity over time. The black and blue bars at the bottom indicate the light regimen. Note that the second light stimulation is shown. For the full time course and a quantification of the first stimulation, see Figure supplement 1. (B) The change in bending upon the second light stimulation was significantly increased when DAN-d1 was activated. (C) As in (B), but for the tail velocity. (D) The average relative importance of each attribute cluster, as provided by the random forest algorithm. Cluster 2 was clearly most important. (E-G) As in (A-C), but for DAN-f1. As no effects of DAN-f1 activation were seen, no random forest was applied. (H-L) As in (A-D), but for DAN-g1. (M-P) As in (A-D), but for a combination of DAN-f1 and DAN-g1. ns above a line indicates non-significance across the whole data set (KW), * and ns above brackets indicate pairwise significance or non-significance, respectively (MW). For the underlying source data and the results of the statistical tests, see Supplementary Data S1.

For DAN-d1, we found that the experimental genotype displayed an increased bending and decreased tail velocity compared to the controls that remained trends for the first stimulation (Fig. 9 – Figure supplement 1A-C) and were statistically significant for the second stimulation (Fig. 9A-D). The effect on bending seemed to be somewhat weaker than for *TH-Gal4* and expressed itself in a delayed peak rather than a general increase. The tail velocity, in contrast, was reduced to zero in both cases (compare Fig. 4D-G and 9A-D). However, using weaker light did not result in any significant differences between the experimental genotype and the controls, in contrast to the case of *TH-Gal4* (compare Fig. 5 – Figure supplement 1A-C and Fig. 9 – Figure supplement 2A-C). Thus, activating DAN-d1 individually has similar effects as activating all neurons covered by *TH-Gal4*, but two stimulations of strong light were required to fully induce these effects.

Next, we repeated the same experiments for DAN-f1 and DAN-g1. We found no effects on locomotion upon activating DAN-f1 (Fig. 9E-G, Fig. 9 – Figure supplement 1D-F, Fig. 9 – Figure supplement 2D-F). For DAN-g1, we found both significantly increased bending and decreased velocity, again only for strong light and clearer during the second activation, just like for DAN-d1 (Fig. 9H-L, Fig. 9 – Figure supplement 1G-I, Fig. 9 – Figure supplement 2G-I). Finally, we activated DAN-f1 and DAN-g1 simultaneously and found increased bending and decreased velocity comparable to the activation of DAN-g1 alone (Fig. 9M-P, Fig. 9 – Figure supplement 1K-M, Fig. 9 – Figure supplement 2K-M).

Taken together, our experiments demonstrated that two of the three tested DANs, when activated individually, induced qualitatively similar effects on bending and velocity that also the combination of almost all DANs induced.

## Discussion

In this study, we found that the optogenetic activation of the same individual DANs can have multiple effects in establishing associative memories, retrieving memories and acutely modulating innate locomotion (Fig. 10A). How can such diverse functions be integrated into the same neuronal circuit?

**Figure 10:**
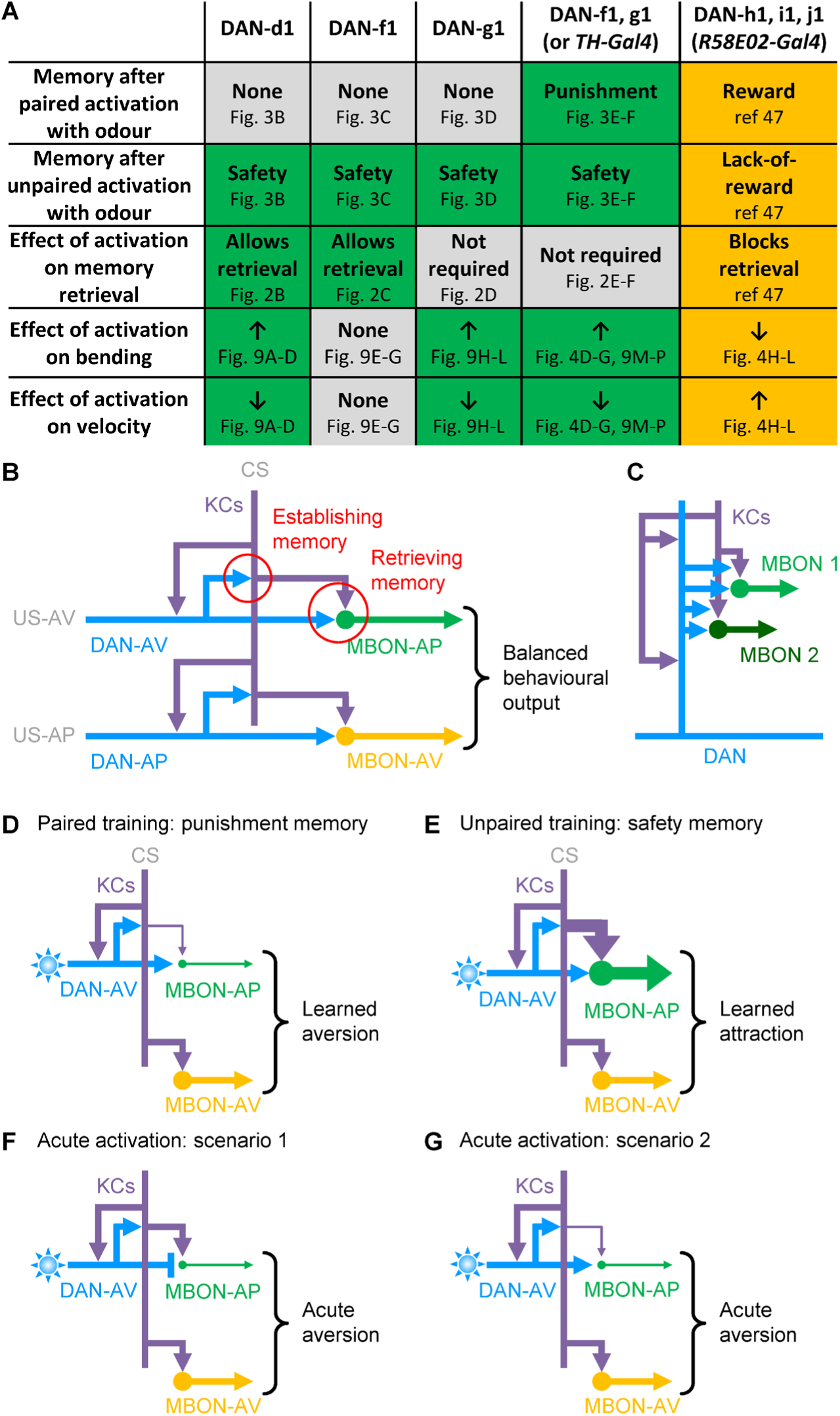
Summary and working hypothesis. (A) A table summarizing the findings of this study as well as of a related recent study^47^ for each DAN. (B) Aversive and appetitive memories are established via synapses between aversive US-signalling DANs (DAN-AV) and appetitive US-signalling DANs (DAN-AP) with KCs. Memories are stored in the strength of the synapses between the KCs and the respective MBONs, and memory retrieval is thought to be controlled via the DAN→MBON synapses. The balance of the MBON signals determines the resulting behaviour (appetitive or aversive). (C) The synapses on the DAN are organized in clusters with DAN→KC and DAN→MBON in the centre, surrounded by KC→DAN synapses. One sample cluster is schematically displayed. For more details, see Figure supplement 1. (D-G) For clarity, only DAN-AV is shown. Thickness of arrows indicate relative strength of synapses and MBON activity, respectively. (D) Paired training with DAN-AV activation depresses the KC→MBON-AP synapses, shifting the balance towards aversion. (E) Unpaired training, in turn, potentiates the KC→MBON-AP synapses, shifting the balance towards approach. (F) DAN-AV activation may directly inhibit the MBON-AP, inducing acute aversive locomotor changes. (G) Alternatively, DAN-AV activation may acutely depress signals from the KC to the MBON-AP.

### A working model for the establishment and retrieval of punishment and safety memories

All the DANs of the mushroom body share a similar circuitry (Fig. 10B). They receive various input outside of the mushroom body neuropil, both from post-sensory interneurons and feedback pathways from the mushroom body output^15,19,25^. A previous analysis of the inputs to the DANs of the DL1 cluster revealed that they integrate gustatory, somatosensory, thermosensory and olfactory input along with internal signals^25^. The two major downstream partners of each DAN are the KCs of the mushroom body as well as the MBONs innervating the same compartment as the DAN^6–7,16,19^ (Fig. 10 – Figure supplement 1A-C). The behavioural effects of the DANs are most likely executed via the MBONs, as they are the only obvious connection from the DANs to motor control^16^. Most evidence, both from larval and adult *Drosophila*, suggest that memories are established via the DAN→KC synapses (see below). We suggest that, in turn, the DAN→MBON synapses may gate the memory retrieval (Fig. 10B). Notably, these two kinds of synapse are not spatially segregated. Rather, DAN→KC, DAN→MBON and KC→DAN synapses form several dense clusters on the DAN, with the DAN→KC and DAN→MBON synapses more to the centre, and the KC→DAN synapses more on the outside of each cluster (Fig. 10 – Figure supplement 1D-E). In many cases, one cluster contains synapses with several different MBONs, and several clusters contain synapses with the same MBON (Fig. 10C). We had demonstrated the same clustered pattern already for DAN-i1^47^. The potential function of this arrangement remains speculative, but it may indicate that these clusters form functional units, each of them with all necessary synaptic connections for both memory formation and memory retrieval.

In the following, we use the d-compartment (lateral appendix) of the mushroom body as example. During training, the presence of an aversive US is signalled by DAN-d1 towards the mushroom body. The coincidence of dopaminergic input and activation of the KC by an odour during paired training is detected by the type I adenylate cyclase and results in presynaptic depression of the KC→MBON synapses (larvae^6–7^; adults^66–67^). This leads to less drive towards approach-promoting MBONs and hence to odour avoidance (Fig. 10D) (larvae^16^; adults^18,59,68–69^). For the case of the larval d-compartment, MBON-d1 was shown to be indeed approach-promoting, MBON-d2 and d3 did not show any behavioural effect, and MBON-p1 has not been tested so far^16^. Thus, the KC→MBON-d1 synapses are the best candidates for harbouring a memory established by DAN-d1.

Presenting an odour unpaired from DAN activation can lead to potentiation of the KC→MBON synapses^17^ (Fig. 10E). The details of this process are not well understood, but at least two scenarios seem plausible: (1) It has been shown that some DANs can confer opposite valence when activated or inhibited, respectively^70^. In our experiments, DANs may be inhibited after a previous US presentation via the feedback neurons from the MBONs to the DANs^16,71^. Alternatively, a decrease in the dopamine signal after activating the DAN could have an effect similar to an inhibition. In either case, this opposite-valence dopamine signal could be associated with the unpaired odour. (2) The same dopamine signal has been shown to have opposite effects on the internal signalling in KCs depending on their activity state^72^. Such an effect could lead to opposite modulations of the KC→MBON synapse, depending on whether the DAN signal is detected while the KC is active (during paired training) or not (during unpaired training). These opposite modulations would drive avoidance or approach towards the odour, respectively. In the case of DAN-d1, only the odour approach after unpaired training, indicating safety learning, was observed (Fig. 3B, 10E).

Previous studies in both adult and larval *Drosophila* have demonstrated that an odour that was paired with the end of an aversive US gains a positive learned valence through a process called relief learning^24,72–74^ (an analogous process has been observed for appetitive US, too^53,75^). The fact that both kinds of learning establish appetitive valences for stimuli paired with the absence (safety) or the end (relief) of the aversive US raises the question whether the same learning process may underlie both observations. As our experiments include three training cycles following each other immediately, the odour was presented several times after the virtual US in each of our unpaired training experiments. However, in a previous study, optogenetic activation of only DAN-f1 but not DAN-d1 supported relief learning^24^, whereas we observed similar levels of safety learning for both neurons (Fig. 3). In a recent study, we observed unpaired memory with an appetitive US even with only a single presentation of first the odour and then the US, and thus, with the odour never being presented after the US^76^. To clarify whether safety learning is a different process from relief learning, an analogous experiment with an aversive US would be required. In any case, it remains possible that these learning processes share the underlying circuit mechanisms.

During the test, the presence of the US is signalled by DAN-d1 and provides a gating signal. This signal is most likely received by the MBONs which are known to be necessary for the expression of learned behaviour (larvae^16,53^; adults^18,77–81^). Such a gating signal in the simplest case could be a logical ‘AND’ gate: for learned behaviour to occur, both the changed drive from the KC onto the MBON and the DAN gating signal to the MBON are required. The consequence is that any associative memory established by the modified KC→MBON-d1 synapses can only take effect behaviourally if DAN-d1 is activated also during the test, just as we observed (Fig. 2B).

### How the mushroom body circuit can account for the modulations of acute locomotion

So far, very little is known about how the mushroom body-innervating DANs could modulate innate locomotion. We consider two simple scenarios to explain our observations, a reduced velocity and increased bending upon activation of DAN-d1 (Fig. 9A-D): (1) the DAN-d1→MBON-d1 synapse acts inhibitory such that an activation of DAN-d1 silences the approach-promoting MBON-d1 (Fig. 10F); (2) the signal from the DAN-d1→KC synapse reduces the drive from the KCs onto the MBON-d1 (Fig. 10G). In both cases, the net output of the mushroom body would be tilted towards aversion.

Scenario 1 seems straight-forward, but two studies in adults caution against such a simple model: First, it was found that activating a DAN while pharmacological blocking output from the KCs activated the MBON of the respective compartment rather than inhibiting it^20^. So, we would need to assume that the “sign” of the DAN→MBON synapses differs either between developmental stages, or between different mushroom body compartments, or even changes in certain conditions. Different dopamine receptors can either excite or inhibit a neuron^82^, and many DANs likely release additional neurotransmitters^19,63,62^. Thus, it remains possible that activation of a DAN causes either activation or inhibition of the downstream MBON dependent on the exact composition of receptors.

Second, a recent study compared the innate valence induced by optogenetically activating DANs (of the PAM cluster) and MBONs and found that in the case of DANs valence was brought about largely by differences in speed in darkness or light, but in the case of MBONs by modulations of turns at the dark-light border^62^. This discrepancy makes a simple, direct inhibition unlikely.

Scenario 2 resembles the effects of the DAN signal in paired learning (Fig. 10D,G). However, the observed modulations of locomotion occurred within a few seconds, much faster than in our learning experiments. In addition, some activation of the KCs and therefore some drive of the MBONs would be required, otherwise an acute down-regulation of the KC→MBON synapse would be without any effect. To which extent the KCs were active during our experiments is not known.

Previous studies provide conflicting evidence for the involvement of the KCs: on the one hand, a recent study in adults found that KC activity was dispensable for the innate preference for PAM-DAN activation^62^. On the other hand, the same study found that knocking down the Dop2R receptor in the KCs reduced the innate preference for PAM-DAN activation^62^. In addition, a study in larvae found an increased locomotion when the Dop1R1 receptor was knocked-down in the KCs^83^.

Considering these discrepancies, it is very likely that both scenarios are too simplistic, by ignoring for example asymmetric body-side effects of DAN and/or MBON activities^47^, or cross-compartmental connections between DANs and MBONs^15^. It can also not be excluded that DANs exert their acute locomotion effects independent of the MBONs. Future research has to disentangle the neuronal mechanisms of how DAN activity modulates innate locomotion and valence.

### Each DAN induces a different combination of effects

Strikingly, each DAN in this study (as well as in a previous study regarding the appetitive domain) induced a different set of behavioural effects (Fig. 10A).

The optogenetic activation of DAN-d1 and DAN-f1 can have two effects with respect to memories: during training, the virtual US signal they confer can induce a safety memory. During testing, this memory is only retrieved when the same DAN is activated (Fig. 2B-C). This mimics many natural US such as tastants or mild mechanical disturbance^32–36^, or formic acid in adult flies^84^. We propose that this mechanism serves to ensure that memories only influence behaviour when it is adaptive to do so.

In contrast, the activation of DAN-g1, as well as of two different combinations of DANs (which both happen to include DAN-g1), establish memories that can be retrieved also without the respective DANs being activated (Fig. 2D-F). This mimics US like electrical shocks that induce memories that are retrieved in absence of the US^85–86^. It has been shown that memories in larvae can be specific for the kind of US and are retrieved independently^39^. It is conceivable that memories regarding relatively harmless types of US, like a bad taste, require the US to be present in order to be retrieved and guide behaviour. Memories regarding more severe US which threaten the body integrity of the animal, such as electrical shocks, are retrieved without the actual US being present. The activation of DAN-d1 or DAN-f1 on the one hand, and DAN-g1 on the other hand, may correspond to such harmless and dangerous US, respectively.

However, as we use optogenetic DAN activation as a virtual US rather than a natural US, our results do not allow for any further conclusions about the content of the established memory. Overall, our knowledge of what kind of information each DAN confers remains limited. Previous physiological studies demonstrated that DAN-d1, DAN-f1 and DAN-g1 respond differently to high concentrations of salt as well as to activation of mechanosensory, nociceptive and multisensory neurons in the ventral nerve cord^15,25^. A systematic overview of how different US are coded across the DANs is still missing. A recent study showed that DAN-g1 is required for aversive learning about high salt concentrations, but not about quinine^25^. As salt memories usually require the presence of salt during the test in order to be retrieved^32–33^, DAN-g1 also appears to be important for learning about such relatively harmless US.

In a previous study we found that the appetitive US-signalling DAN-i1 can establish memories of opposite valence when it was activated paired or unpaired with an odour^47^. This suggested that both kinds of memory rely on the activity of the same DANs. In the aversive domain, DAN-d1, DAN-f1 and DAN-g1 each establish only safety memory but not punishment memory, whereas activating combinations of DANs did establish both kinds of memory^15^ (Fig. 3). The reason for these discrepancies is not known yet. To our knowledge, no natural US is known that only induces safety, but no punishment memory, in larvae. This may suggest that all natural US tested so far activate combinations of DANs.

Most strikingly, DAN-d1 and DAN-g1 (but not DAN-f1) also modulate innate locomotion (Fig. 4, 9). Interestingly, recent studies in mammals suggested that the nigro-stratial ‘movement’ pathway also plays a role for carrying reward signals, in addition to the traditional mesolimbic ‘reward’ pathway^87^. Together, these results point towards a neuronal organization in which the same dopaminergic neurons can have multiple functions that are as diverse as instructing learning processes and modulating movements. Whether these multiple functions are rooted in different molecular mechanisms within the dopaminergic neurons or are brought about by different downstream neurons receiving the same dopaminergic signals remains to be resolved.

A recent study in adult flies focusing on the DANs of the PAM cluster found largely separated functions for different DAN subsets^60^: activating all PAM-DANs (*R58E02-Gal4*-DANs) induced appetitive memories, acute avoidance, modulations of locomotion and decreased feeding. Specific subsets of the adult PAM-DANs induced appetitive learning and acute preference but had very little effects on locomotion. Another subset increased feeding, but induced no memory, no acute preference and no effects on locomotion^60^.

In larvae, *R58E02-Gal4*-DANs also induce appetitive memories^27^, but we observed acute approach rather than aversion, consistent with another study in adults^62^ (Fig. 7C, Fig. 7 – Figure supplement 1A). For now, the reasons for these discrepancies remain unclear. As we did not test individual pPAM-DANs for acute preference or locomotion, it remains possible that in both larvae and adults some of the pPAM/PAM neurons promote approach and others avoidance, yet the net effect of activating all of them differs. Notably, the DL1 and pPAM clusters have very different fates during metamorphosis. The four DL1-DANs will be part of the PPL1 cluster in adults and either continue to provide input to the adult mushroom body or trans-differentiate to innervate other adult brain regions^88^. In total, adults have eight types of PPL1-DANs, with one or two neurons per celltype^71^. The four larval pPAM-DANs, in contrast, die during metamorphosis and are replaced by about 120 adult PAM-DANs^71,88^. It is possible that the larger number of PAM-DANs promotes more functional distinctions compared to a more overlapping-function organization across the few DL1/PPL1-DANs. Alternatively, it is possible that the larval pPAM-DANs are functionally equivalent to just one subset of the adult PAM-DANs, promoting appetitive learning and acute approach, whereas other PAM-DANs have different sets of functions. The larval equivalent of those neurons, if it exists, has not been identified yet.

Our results suggest a combinatorial-like neuronal organization, at least within the DL1 cluster, with each neuron having a unique yet overlapping set of functions. This raises a question: given that all mushroom body compartments share the same basic circuitry^19^ (Fig. 10B), how can such functional differences be explained? It is conceivable that they depend on the specific properties of the involved MBONs or its synapses with the KCs and the respective DANs. For example, if the synapses between the KCs and a given MBON are weak to begin with (for example the MBONs of the d- and f-compartment), they can be easily potentiated by unpaired training, but any depression by paired training will be minute. Such weak depression may need to happen in several compartments before it becomes behaviourally measurable.

Moreover, the impact of the gating signal by the DAN to the MBON may vary dependent on the circumstances: the memories established by DAN-d1 or DAN-f1 bypassed the retrieval control by the DANs if those memories were established with CsChrimson, but not with ChR2-XXL (Fig. 2 – Figure supplement 3)^15^. The memory established by DAN-g1 bypassed this control even when using ChR2-XXL (Fig. 2D). Little is known about the molecular mechanisms of memory retrieval in MBONs, and future research has to uncover which factors may determine its dependence on dopamine signals.

Our study revealed that individual DANs can induce surprisingly diverse effects, from establishing and retrieving memories to modulating innate locomotion. The mushroom body circuit – that is shared with adult flies and other insects – provides an excellent study case to advance our understanding of the complex functions of the dopaminergic system. Given the highly conserved roles of DANs in mediating “teaching” signals and modulating movement across the animal kingdom including vertebrates and humans, a deeper understanding of this circuit and its functions should provide new insights about the principles of DAN function in general.

## Materials & Methods

### Animals

Third-instar feeding-stage larvae (*Drosophila melanogaster*), aged 4-5 days after egg laying, were used throughout. Flies were maintained in mass culture at 25 °C, 60-70 % relative humidity and a 12/12 hour light/dark cycle. Two kinds of food medium were used:

Food medium #1 was used for all learning and olfactory experiments as well as for some of the locomotion experiments. Per 1 l water, 45 ml sugar beet molasses, 45 ml malt, 147.5 g cornmeal, 10 g soy flour, 18.75 g dry yeast, 6.25 g agar and 2.5 g methyl-4-hydroxybenzoate (Merck, Darmstadt, Germany) were added as described in detail previously^89^.

Food medium #2 was used for most of the locomotion experiments as well as the olfactory stimulation preference experiments. Per 1 l water, 100g glucose, 50 g cornmeal, 31.65 g wheat germ, 40 g dry yeast, 8 g agar, 2 ml propionic acid (Kanto-kagaku, Tokyo, Japan) and 5 ml 10% butyl p-hydroxybenzoate (Wako, Osaka, Japan) in ethanol were added as described in detail previously^90^.

We took a spoonful of food medium from a food vial, randomly selected the desired number of larvae, briefly rinsed them in tap water and started the experiment.

We used either wild-type larvae from the Canton Special genotype, or transgenic larvae to express light-gated ion channels in the respective DAN. Towards this end, one of two effector strains, *UAS-ChR2-XXL* (Bloomington Stock Center #58374), or *20xUAS-CsChrimson-mVenus* (Bloomington Stock Center #55134), was crossed to either *TH-Gal4* (Bloomington Stock Center #8848), *R58E02-Gal4* (Bloomington Stock #41347), or to one of four split-Gal4 driver strains: MB328B, MB054B, SS02180, or SS01716^15^ to obtain double-heterozygous offspring. As driver controls, the driver strains were crossed to a local copy of *w^11^*^18^ (Bloomington Stock Center #3605, 5905, 6326). As effector control, a strain carrying the landing sites used for the split-Gal4 (attP40/attP2), yet without a Gal4 domain inserted (“empty”)^91^, was crossed to the respective effector. When using *TH-Gal4*, we used *w^1118^* instead. As CsChrimson requires feeding of all-trans-retinal^56^, larvae that were crossed to *20xUAS-CsChrimson-mVenus* were raised in standard food medium that additionally contained 400 µM final concentration of all-trans-retinal (CAS: 116-31-4, Sigma-Aldrich). For the TH-RNAi experiments, a double effector strain was used, made from the *UAS-ChR2XXL* strain and *y[1]v[1];P{TRiP.JF01813}attP2* (Bloomington Stock #25796) to express a double-strand-RNA element of the *ple* gene^92^. All larvae used for optogenetic experiments were raised in constant darkness.

For the whole-body anatomy, we crossed each driver strain to a *y[1] w*;;MB247-mCherry-CAAX, UAS-mIFP-T2A-HO1* double-effector strain^28^ (called *MB247-mCherry-CAAX, UAS-mIFP* in the main text). As driver control, we crossed each driver strain to *y[1] w*;;MB247-mCherry-CAAX*^28^, as effector control, we crossed *y[1] w*;;MB247-mCherry-CAAX, UAS-mIFP-T2A-HO1* to *y[1] w\** (Bloomington Stock Center #1495).

### Experimental setup

For learning and preference experiments, larvae were trained and tested in Petri dishes of 9 cm inner diameter (order #82.1472, Sarstedt) filled with 1 % agarose (electrophoresis grade; CAS: 9012-36-6). For the experiments displayed in Fig. 2A and 3A, 5 mM quinine (purity: 92%, CAS: 6119-70-6, Sigma-Aldrich) was added into the agarose. As odour, we used *n*-amyl acetate diluted 1:20 in paraffin oil (AM; CAS: 628-63-7, Merck). For locomotion experiments, either Petri dishes of 15 cm inner diameter (order #82.1184.500, Sarstedt) filled with 1 % agarose were used, or dishes of 9 cm inner diameter (order #3-1491-01, AS ONE) filled with 2 % agar (CAS: 9002-18-0, Fujifilm Wako Pure Chemical Industries).

Experiments were performed inside a 43 x 43 x 73 cm surrounding box equipped with a custom-made light table featuring a 24 x 12 LED array (Solarox) and a 6 mm thick diffusion plate of frosted plexiglass on top to ensure uniform blue light for ChR2-XXL activation (470 nm; 120 µW/cm²; for Fig. 2 – Figure supplement 3C, Fig. 4 incl. Figure supplements, Fig. 7, Fig. 8 incl. Figure supplements, Fig. 9 and Fig. 9 – Figure supplement 1: 1000 µW/cm²) or red light for CsChrimson activation (630 nm; 430 µW/cm^2^). Petri dishes were placed directly on top of the diffusion plate. For additional details regarding the experimental setup, see^53^.

### Associative learning

Learning experiments followed established protocols^93^. Odour containers were prepared by adding 10 µl of odour substance into custom-made Teflon containers (5 mm inner diameter with a lid perforated with 7 holes of 0.5-mm diameter each). Petri dishes were covered with modified lids perforated in the centre by 15 holes of 1 mm diameter each to improve aeration.

For paired training with quinine as US, about 20 larvae were placed on a quinine-filled Petri dish equipped with two AM-filled odour containers. After 2.5 min, the animals were transferred to a dish with plain agarose and empty odour containers (EM) on which they spent another 2.5 min. This cycle (AM-/EM) was repeated two more times before the animals were transferred to a test Petri dish with AM on one and EM on the other side. This dish was either a plain agarose dish, or one with quinine added, as mentioned in the results section. After three minutes, the number of animals on either side as well as on a 1 cm-wide middle stripe were counted and the odour preference was calculated as:

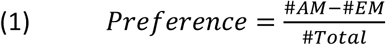

Thus, preference values are constrained between 1 and −1 with positive values indicating a preference for and negative values indicating avoidance of AM. Note that the training sequence in every other repetition of the experiment was reversed (EM/AM-)

For unpaired training, AM was presented on plain agarose Petri dishes and EM was present on quinine Petri dishes, such that the CS and US were experienced explicitly separated from each other (AM/EM- or EM-/AM). Otherwise, the training was performed analogously, and the final test was performed as described.

From each one paired and one unpaired trained cohort of animals we calculated an associative Memory score as:

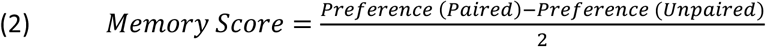

Thus, the values can range from 1 and −1 with negative values indicating that the larvae preferred the odour less after paired than unpaired training. Thus, negative scores indicate an aversive associative memory. Positive values, in turn, would indicate appetitive associative memory.

Learning experiments with ChR2-XXL or CsChrimson as effectors were performed analogously. However, all Petri dishes were filled with plain agarose, i.e. no tastants were added. Rather, for paired training, AM was paired with continuous blue (ChR2-XXL) or red light stimulation (CsChrimson) for 2.5 min, followed by 2.5 min of darkness without an odour. For unpaired training, animals received odour presentation and light stimulation separately from each other. The test was performed as described, either in darkness or with light stimulation as mentioned in the results section.

### Innate olfactory behaviour

Odour containers and Petri dishes were prepared as described above. Cohorts of about 20 larvae were collected from the vial, briefly washed in tap water and placed to a Petri dish with an AM container on one side and an empty container on the other side. After 3 minutes, the innate odour preference was determined according to equation (1).

### Locomotion experiments

Cohorts of about 8 larvae were placed on the middle of a Petri dish filled with agarose or agar. The dish was placed in the experimental chamber and video-recording was started after about 10 s. After 30 s of video-recording in darkness, light was switched on for 30 s followed by 60 s of darkness. In some experiments, 2 or 3 light stimulation cycles were applied. Blue or red light was used as mentioned along with the results.

Experiments involving feeding 3IY followed established protocols^57^. Before the experiment the larvae were transferred to a yeast paste with or without added 3-Iodo-L-tyrosine (3IY; stored at −20°C; CAS: 70-78-0, Sigma-Aldrich) at a concentration of 5 mg/ml. After 4 hours, cohorts of 8 larvae were collected from the yeast paste and the experiment was conducted as described.

### Optogenetic stimulation preference experiments

In order to test for the animals’ preference for having sets of DANs activated, we prepared Petri dishes filled with agar of which one half was illuminated by blue or red light, the other half was kept in darkness. Cohorts of about 10 larvae were placed on the dark half of the dish about 1.5 cm from the midline. Their behaviour was video-recorded for 3 minutes before the animals were counted and the preference was calculated as:

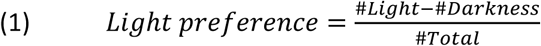

Thus, preference values are constrained between 1 and −1 with positive values indicating a preference for and negative values indicating avoidance of the light.

### Video analysis

Video analysis used the IMBA (Individual Maggot Behaviour Analyser) software and followed previously described procedures^51^. In brief, the positions of head, tail, as well as several spine and contour points of each larva were tracked over the course of the experiment. A combination of computational modelling and statistical approaches allowed keeping the identities of larvae across collisions and thus the analysis of individual animals tested in cohorts. Then, we discarded all tracks that were shorter than half of the time window used for the analysis. This prevented multiple tracks from the same individual animal from entering the analysis.

Larval behaviour was separated into head-casting (HC) phases and run phases. Lateral HCs are a very prominent feature during larval chemotaxis. We quantified these head movements based on the angular speed of the head vector through the front third of the animal. HCs were detected when the angular speed exceeded ± 35°/s. Runs were determined as periods in which the animals were not performing HCs, omitting 2 s before and after each HC event to prevent behavioural effects of the

HC. We noticed that the rhythmic velocity changes of the tail point provided a reliable indicator for the occurrence of one peristaltic forward movement. We therefore defined each local maximum in the tail forward velocity as one step, and the period between two maxima, i.e. one complete peristaltic cycle, as an inter-step (IS) period. Since the algorithm we employed to detect local-maxima also produced false maxima during intervals of little or no forward motion, we discarded maxima whenever the tail velocity in forward direction was less than 0.6 mm/s.

After the detection of runs, steps and head casts, we calculated several behavioural attributes.

We summarize all attributes in this study in table 1. Whenever the average behaviour of a group of larvae is displayed over time, the mean and the 95 % confidence intervals of the respective attribute are calculated per 1 s time bin with a 5 s convolution filter. Whenever data are displayed in box plots to compare across experimental conditions, the mean of each attribute is calculated for each individual larva across the indicated time period. As we were interested in the difference in a given attribute X before and during light stimulation, we calculated Δ-values as the difference in X during the last 15 s of light stimulation minus X during the 15 s just before light stimulation (called baseline in the main text).

We used random forest models which are based on classification and regression trees^94–95^. The random forest algorithm works by building N number of decision trees and then predicting the class label of an instance using a majority voting. Each tree is trained on a random subspace of the data, only using a sample of the instances. To evaluate the random forest classification, we used five-fold cross validation, which was repeated three times. Five-fold cross validation means that the test set for the model is taken from five different, disjunct sets of the data set. The data set is then shuffled and the procedure is repeated three times. Cross validation is used to make sure that the model is not based merely on the selected data but can explain the full data set.

For the applications of the random forest in this study, we pre-selected 17 behavioural attributes to be included (Table 1). The pre-selection was based on our previous study^51^ and was warranted to avoid using largely redundant attributes and to exclude non-behavioural attributes such as the size of the larvae. The random forest was always applied to pairwise compare two experimental conditions, typically the experimental groups with one of the genetic controls. In order to provide a good estimate of the importance of each attribute, the complete random forest model (including the cross validation) was performed 10 times for any given comparison. In the typical case, this means 10 repetitions of the comparison between the experimental genotype with the driver control and 10 repetitions of the comparison with the effector control. In each case, the relative importance rank of each attribute was noted.

### Whole-body anatomy

Experimental procedures followed previous work^28^. In brief, third-instar larvae were first bleached (4% sodium hypochlorite, order #9062.1, Roth) for 10 min at room temperature. After washing (3 × 5 min in distilled water), they were fixed in 4% PFA in 0.1 mol/L PB at pH 9, with gentle shaking overnight at 4°C. Fixed samples were briefly rinsed 3 × with 0.1 mol/L PB containing 0.2% Triton X-100 (PBT), then washed 2 × 60 min and left overnight at 4°C in PBT. Dehydration the following day used a graded ethanol (EtOH) series (60 min in 10% and 25% EtOH, followed by 30 min in 50%, 60%, 80%, and 2 × 100% EtOH). Samples were then cleared by replacing EtOH with ethyl cinnamate (ECi; ethyl 3-phenyl-2-propenoate; order #112372-100G, Sigma-Aldrich). After 30-60 min, the ECi was refreshed once, and the samples stored at room temperature in black boxes in a desiccator.

The samples were placed in ECi-cleared phytagel blocks (1 × 1 × 1 cm, P8169, Sigma-Aldrich) before imaging on an Ultra-Blaze microscope (Miltenyi Biotec). The microscope was equipped with an FIU-15 white light laser (NKT Photonics) for excitation through triple-sheet optics to illuminate samples from one side. The following excitation and emission filters were used: autofluorescence: exc. 480/20, em. 515/30, mcherry: exc. 590/20, em. 625/30, mIFP: exc. 680/20, em. 720/30.

Tiled image stacks were acquired with either a 12× objective with zoom set to 1 or 4x objective with zoom set to 1.66 (both Miltenyi Biotec) with a format of 2048 × 2048. Using ImSpector software (version 7.3.6, Miltenyi Biotec), a 10% overlap of tiles was set for stitching.

Image files were processed with Imaris software (version 10.1, Bitplane), including file conversion, stitching, further processing, and rendering. Either the ortho slicer, oblique slicer or clipping plane tool was used to restrict volumes in the z direction to improve the representation of structures that were otherwise covered in the context of the whole body. All inserts were generated using the snapshot function in Imaris. Whole-body 2D maximum-intensity projections were generated in Fiji^96^. Supplementary Movie 1 was produced in Imaris with the key frame animation tool; Adobe Premiere Pro 2020 (version 14.9.0, Adobe) was used for cutting and labelling.

### Dendrogram analysis

To visualize neuron morphology and synapse locations, we employed neuron dendrograms^97^ that are linearized, topologically correct 2D representations of 3D neuron reconstructions with synapse annotations (based on ^19^). Synapse locations and the distances between them are approximately correct, with slight adjustments to avoid overlap and cluttering of the synapse symbols^97^.

We formally analysed the spatial organization of synapses based on geodesic distances along the neuron’s branches (“cable length distances”). The geodesic distance between two synapses was determined by finding the shortest path between them on the neuron, and then computing the cumulated Euclidean distance, in the 3D coordinate space of the neuron, between points along that path.

In particular, we analysed spatially local clusters of synapses and the distances to the centroid of the cluster. To this end, we clustered the synapses on each neuron based on geodesic distances and using hierarchical clustering with Ward’s criterion^98^ (implemented in the base R package “stats”). The maximum number of clusters per neuron was set to k=10. For analysis, we kept only clusters with at least 5 presynaptic and 5 postsynaptic KC connections.

The centroid of a cluster was defined to be the most central synapse, computed by partitioning around medoids clustering (PAM with k = 1 cluster)^99^ based on geodesic distances.

## Quantification and statistical analysis

As behavioural data are often non-normally distributed, we used two-tailed non-parametric tests throughout. Our results are displayed as box plots which allow for a simple visualization of the data distribution independent of the sample size. The median is presented as the middle line and 25 %/75 % and 10 %/90 % as box boundaries and whiskers, respectively, as these descriptors are less sensitive to outliers than the mean. Values were compared across multiple groups with Kruskal-Wallis tests (KW). In case of significance, subsequent pair-wise comparisons used Mann-Whitney U-tests (MW). To test whether values of a given group differ from zero, we used Wilcoxon Signed-Rank tests (WSR). When multiple comparisons were performed within one analysis, a Bonferroni-Holm correction was applied to keep the experiment-wide error rate below 5 %^100^. The detailed results of all statistical tests are reported in the Supplementary Data S1 along with the source data. Experimenters were blind to genotype. Sample sizes (biological replications) were chosen based on previous studies revealing moderate to mild effect sizes^47,51^ and are displayed in the figures.

## Acknowledgements

This study received institutional support by the Otto von Guericke Universität Magdeburg, the Wissenschaftsgemeinschaft Gottfried Wilhelm Leibniz (WGL), the Leibniz Institute for Neurobiology (LIN), the Hokkaido University, and the RWTH University Aachen. This study received grant support from the Deutsche Forschungsgemeinschaft (DFG) (GE 1091/4-1 and FOR 2705 *Mushroom body* [to B. Gerber]), the CRCNS program of the German Federal Ministry of Education and Research (BMBF) “DrosoExpect” (01GQ2103A [to B. Gerber]), the Japan Society for the Promotion of Science (JSPS) (Kakenhi 22K20673), the Goho Life Sciences International Fund, and the German-Israeli Foundation for Science (GIF) (G-2502-418.13/2018). Experimental contributions of J. Chen, R. Khondaker, T. Meyer, S. Schuller, M. Thane, L. Warzog, B. Fischer, technical assistance by M. Dombach, M. Paisios, M. Thane and F. Unterstab, as well as discussions with C. Eschbach, B. Gerber, C. König, N. Mancini, N. Tanaka, A. Thum and A. Yarali are gratefully acknowledged. We thank C. Eschbach, C. König, A. Thum, D. Weber and M. Zlatic for sharing fly strains and results prior to publication.

## Author contributions

Conceptualization: Michael Schleyer, Naoko Toshima.

Formal analysis: Michael Schleyer, Arman Behrad, Franziska Behnke.

Investigation: Naoko Toshima, Arman Behrad, Franziska Behnke, Gauri Kaushik, Aliće Weiglein,

Martin Strauch, Juliane Thoener, Oliver Kobler, Maia Lisandra M. Wang, Ayaka Fukushima, Markus Dörr.

Data curation: Michael Schleyer, Naoko Toshima, Arman Behrad, Aliće Weiglein, Franziska Behnke.

Writing - original draft: Michael Schleyer, Naoko Toshima, Aliće Weiglein, Franziska Behnke.

Writing - review & editing: Naoko Toshima, Aliće Weiglein, Arman Behrad, Franziska Behnke, Gauri Kaushik, Martin Strauch, Juliane Thoener, Oliver Kobler, Maia Lisandra M. Wang, Ayaka Fukushima, Markus Dörr.

Visualization: Michael Schleyer, Martin Strauch, Juliane Thoener, Oliver Kobler.

Supervision: Michael Schleyer, Juliane Thoener.

Project administration: Michael Schleyer.

Funding acquisition: Michael Schleyer.

## Data availability

Source data to all behavioural experiments are included as supplementary material (Supplementary Data S1).

---

**Figure 2 - Figure supplement 1:**
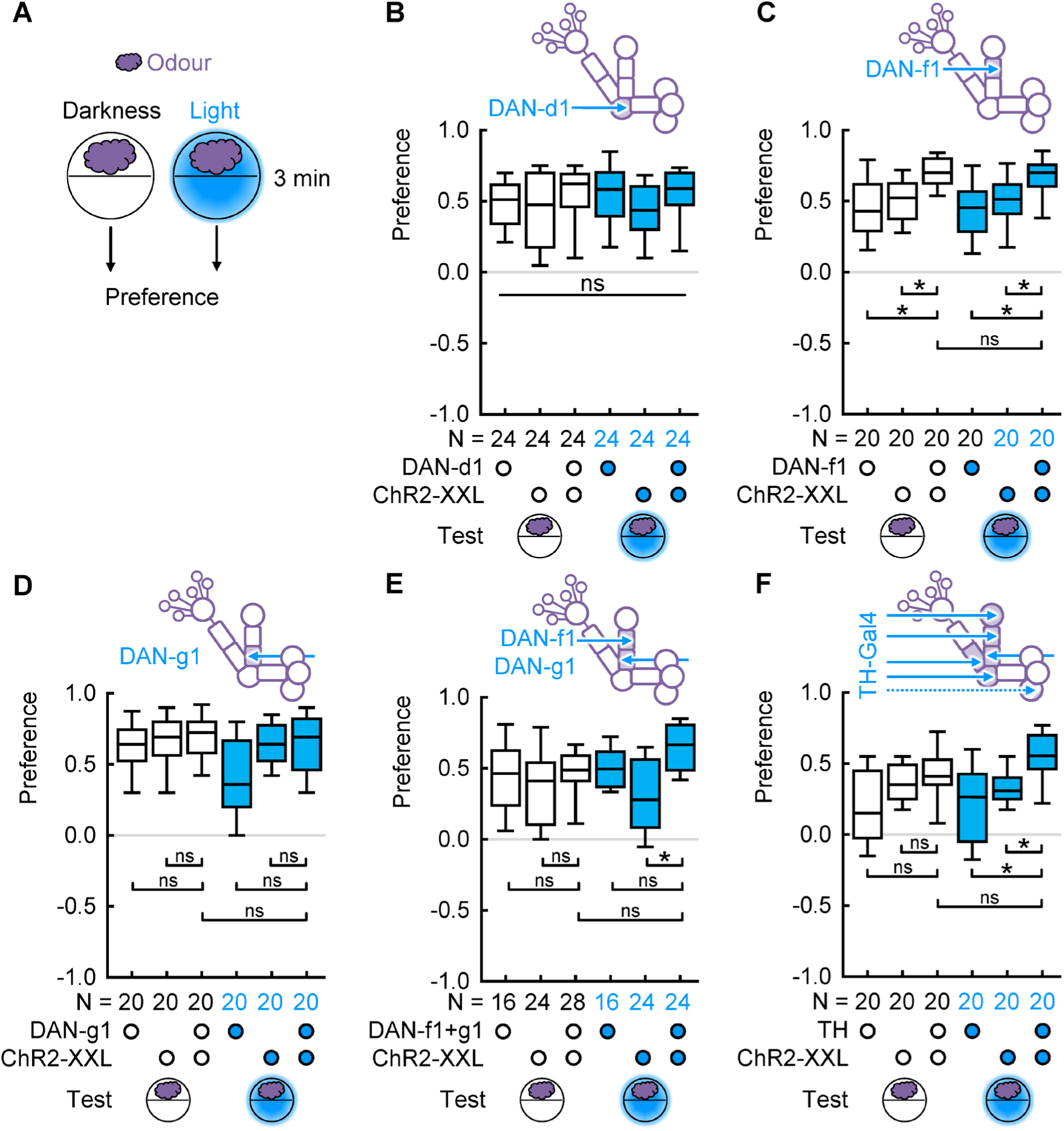
Innate odour preferences. (A) Groups of about 20 larvae were tested for three minutes for their odour preference, either in darkness or blue light. (B) Innate odour preferences of the genotypes used in Fig. 2B, tested in darkness or blue light. No differences were found. (C) Innate odour preferences of the genotypes used in Fig. 2C. The experimental genotypes displayed a higher odour preference than the controls, independent of the testing condition. Note that this cannot explain the difference in memory score between testing conditions found in Fig. 2C. (D) Innate odour preferences of the genotypes used in Fig. 2D. No differences were found. (E) Innate odour preferences of the genotypes used in Fig. 2E. When tested in light, the experimental genotype displayed a higher odour preference than the effector. Note that this cannot explain the difference in memory score between genotypes tested in darkness as seen in Fig. 2E. (F) Innate odour preferences of the genotypes used in Fig. 2F. When tested in light, the experimental genotype displayed a higher odour preference than the controls. Note that this cannot explain the difference in memory score between genotypes tested in darkness as seen in Fig. 2F. ns above a line indicates non-significance across the whole data set (KW), * and ns above brackets indicate pairwise significance or non-significance, respectively (MW). For the underlying source data and the results of the statistical tests, see Supplementary Data S1.

**Figure 2 - supplement 2:**
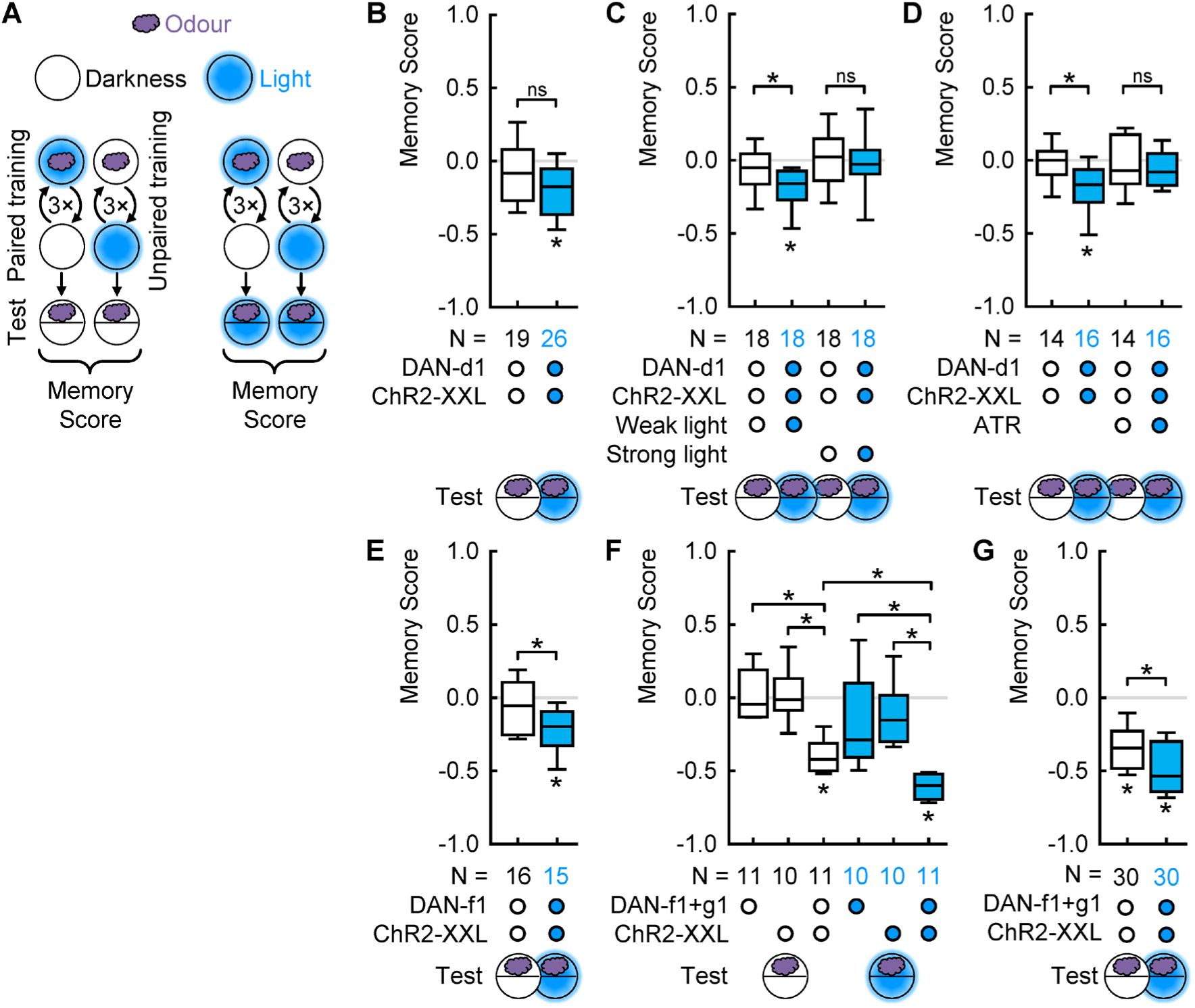
Replications of the experiments shown in Fig. 2. (A) Larvae were trained with odour presented either paired or unpaired with blue light activation and tested either in darkness or in blue light. (B) Larvae of the experimental genotype, expressing ChR2-XXL in DAN-d1, showed associative memory only when tested in presence but not absence of the blue light. (C) Larvae of the experimental genotype showed associative memory only when trained with the standard weak light and tested in presence of light (of the same strength). When they were trained and tested with a higher light intensity, no memories were observed. (D) Larvae of the experimental genotype showed associative memory only when raised in standard food and tested in presence of light. When they were raised in food with all-trans-retinal added, which has been shown to increase the light-sensitivity of ChR2-XXL^42^, no memories were observed. (E) Larvae expressing ChR2-XXL in DAN-f1 showed associative memory only when tested in presence but not absence of the blue light. (F) Only the larvae of the experimental genotype, expressing ChR2-XXL in DAN-f1 and DAN-g1, showed associative memory independent of the presence or absence of the US. The genetic controls displayed no significant associative memory. (G) Larvae of the experimental genotype showed associative memory only when tested in presence but not absence of the blue light. * and ns above brackets indicate pairwise significance or non-significance, respectively (MW), * below boxes indicate memory scores significantly different from zero (WSR). For the underlying source data and the results of the statistical tests, see Supplementary Data S1.

**Figure 2 - supplement 3:**
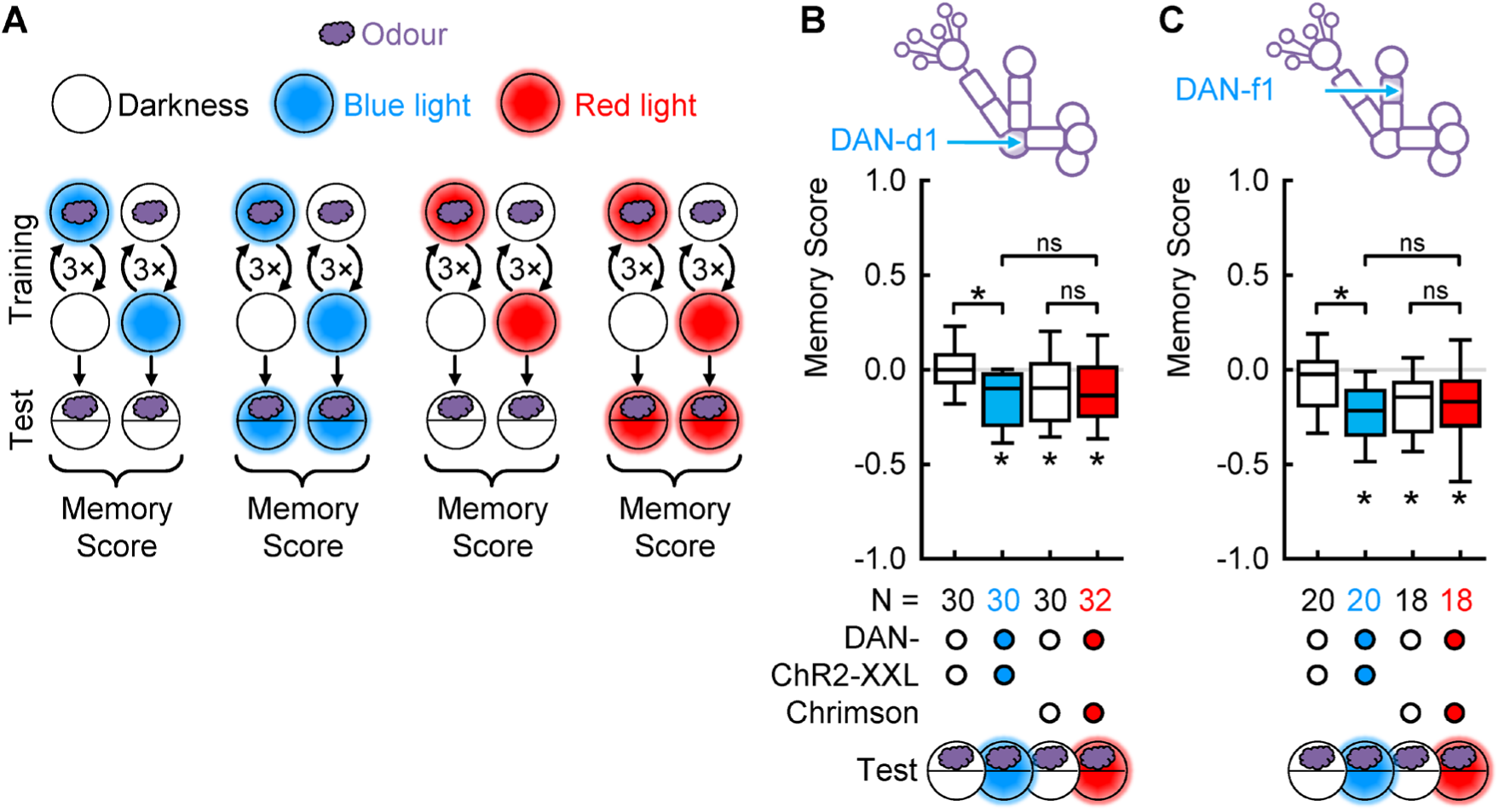
DAN activation with CsChrimson establishes memories that are retrieved independent of DAN activation during the test. (A) Larvae were trained with the odour presented either paired or unpaired with blue (*UAS-ChR2-XXL*) or red (*UAS-CsChrimson*) light activation and tested either in darkness or the respective light. (B) In a replication of the experiment shown in Fig. 2B, larvae displayed an associative memory only if DAN-d1 was activated also during the test (left). However, if instead of ChR2-XXL the red-shifted channel CsChrimson was used as effector, associative memory scores were observed independent of the activation of DAN-d1 during the test. (C) As in (B), but for DAN-f1. * and ns above brackets indicate pairwise significance or non-significance, respectively (MW), * below boxes indicate memory scores significantly different from zero (WSR). For the underlying source data and the results of the statistical tests, see Supplementary Data S1.

**Figure 4 - supplement 1:**
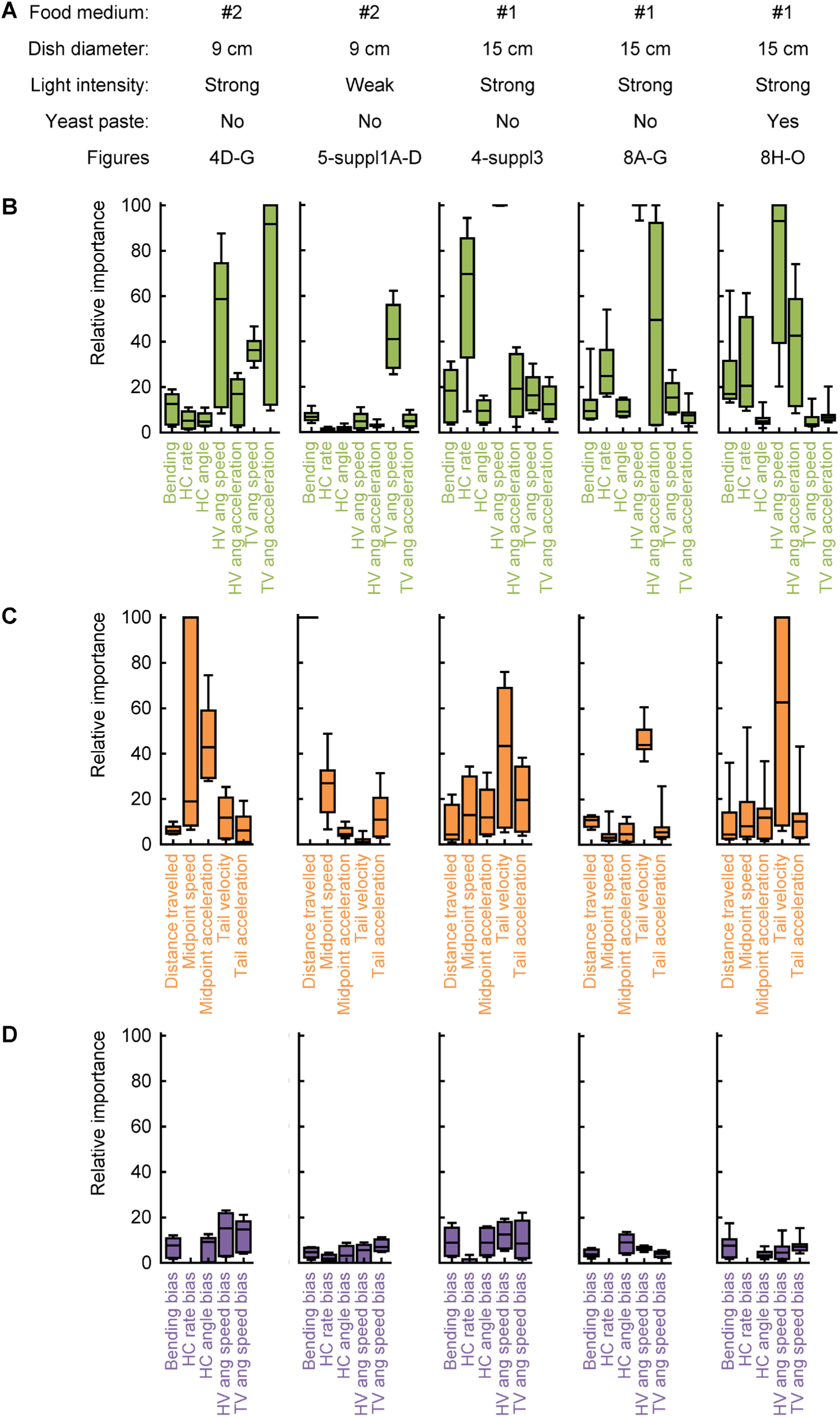
Random forest analysis of five independent repetitions of the same locomotion experiment. (A) Summary of the experimental conditions of five independent replications of the locomotion experiment with *TH-Gal4* and *ChR2-XXL*. For the right-most experiment, the larvae were placed on a yeast paste for 4 hours before the experiment (see Materials & Methods for details). (B-D) Each column shows the relative importance based on the random forest algorithm of a given experimental replication. (B), (C), (D) show the attributes of cluster 1, 2 and 3, respectively.

**Figure 4 - supplement 2:**
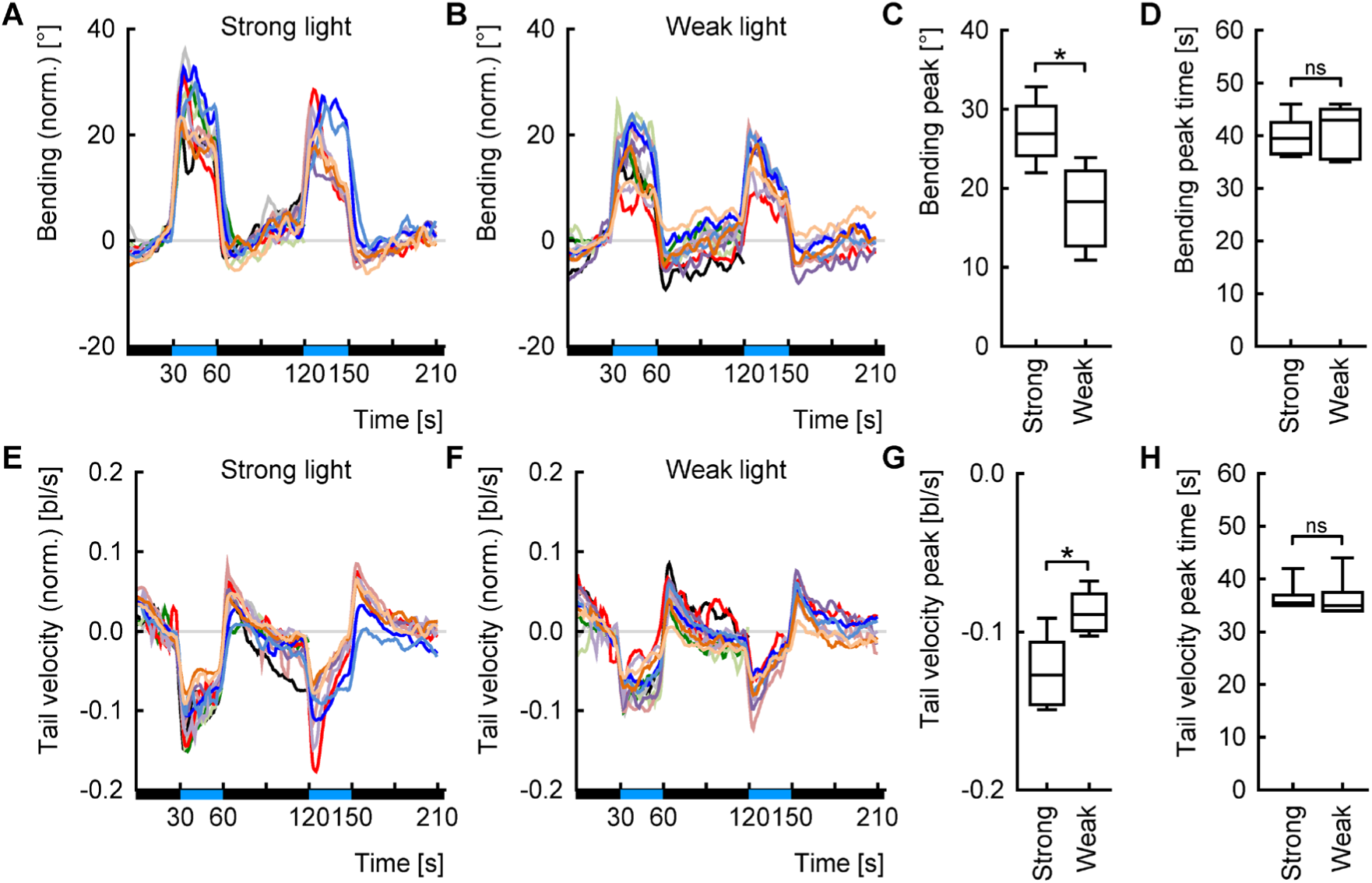
Light responses of genetic controls. (A) Bending over time. Black and blue bars at the x-axis indicate the light regimen. Each line represents the average bending of one genetic control of one of this study’s experiments using strong blue light (specifically, the experiments shown in Fig. 4, 9). The curve of each control was normalised to its bending during the time period between 15 and 30 s to allow comparisons of the shape of the curves independently from the baseline behaviour. Please note that some experiments included only one cycle of light stimulations (these curves end at time point 120 s), others included two cycles. (B) As in (A), but for experiments using weak blue light (shown in Fig. 5 – Figure supplement 1, Fig. 9 – Figure supplement 2). (C) The highest value during the first stimulation of each curve was determined. This peak value was higher for strong than for weak light. (D) The time until the peak of the first stimulation of each curve. No differences were seen between the light intensities. (E-H) As for (A-D), but for the tail velocity. As “peak” value, the lowest tail velocity of each curve during the first stimulation was used. * and ns above brackets indicate pairwise significance or non-significance, respectively (MW). For the underlying source data and the results of the statistical tests, see Supplementary Data S1.

**Figure 4 - supplement 3:**
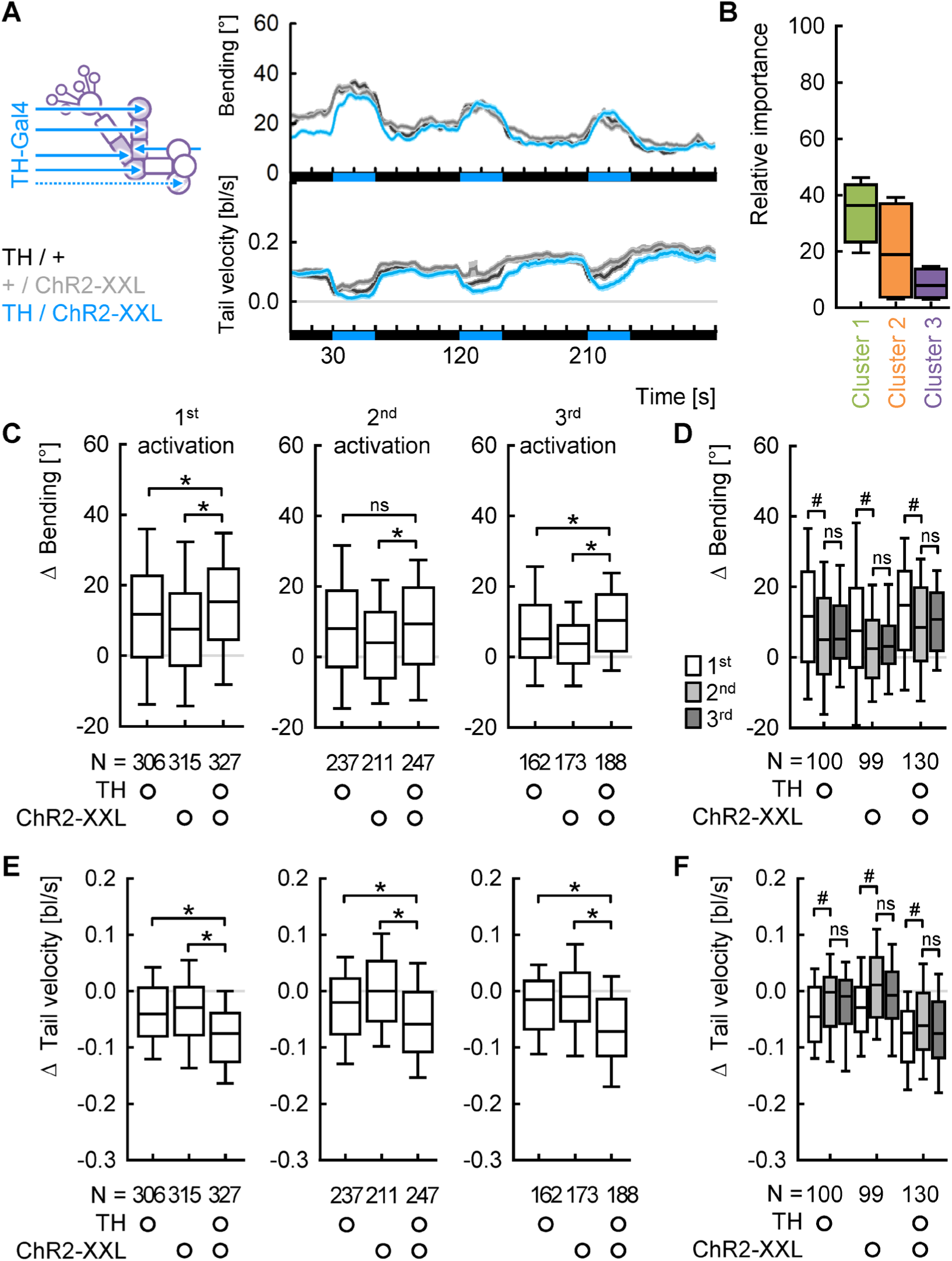
Replication of the experiment shown in Fig. 4D-G with three stimulation cycles. (A) Average bending and tail velocity over time. Larvae of all genotypes increased bending and decreased tail velocity upon light stimulation. Both effects were enhanced in the experimental genotype. The black and blue bars at the bottom indicate the light regimen. (B) The average relative importance of all behavioural attributes within each cluster, as provided by the random forest algorithm. (C) A Δ bending value was calculated in each individual to quantify the change in bending upon light stimulation. This change was significantly increased in the experimental genotype for each light stimulation. (D) As in (C), but only including individuals that were recorded during all three light stimulations. Differences across light stimulations are compared within individuals. The change in bending was reduced for the second and third stimulation in all genotypes. (E-G) As in (C-D), but for the tail velocity. * and ns above brackets indicate pairwise significance or non-significance, respectively (MW). In (D) and (F), # and ns above brackets indicate pairwise within-animal significance or non-significance, respectively (WSR). For the underlying source data and the results of the statistical tests, see Supplementary Data S1.

**Figure 5 - supplement 1:**
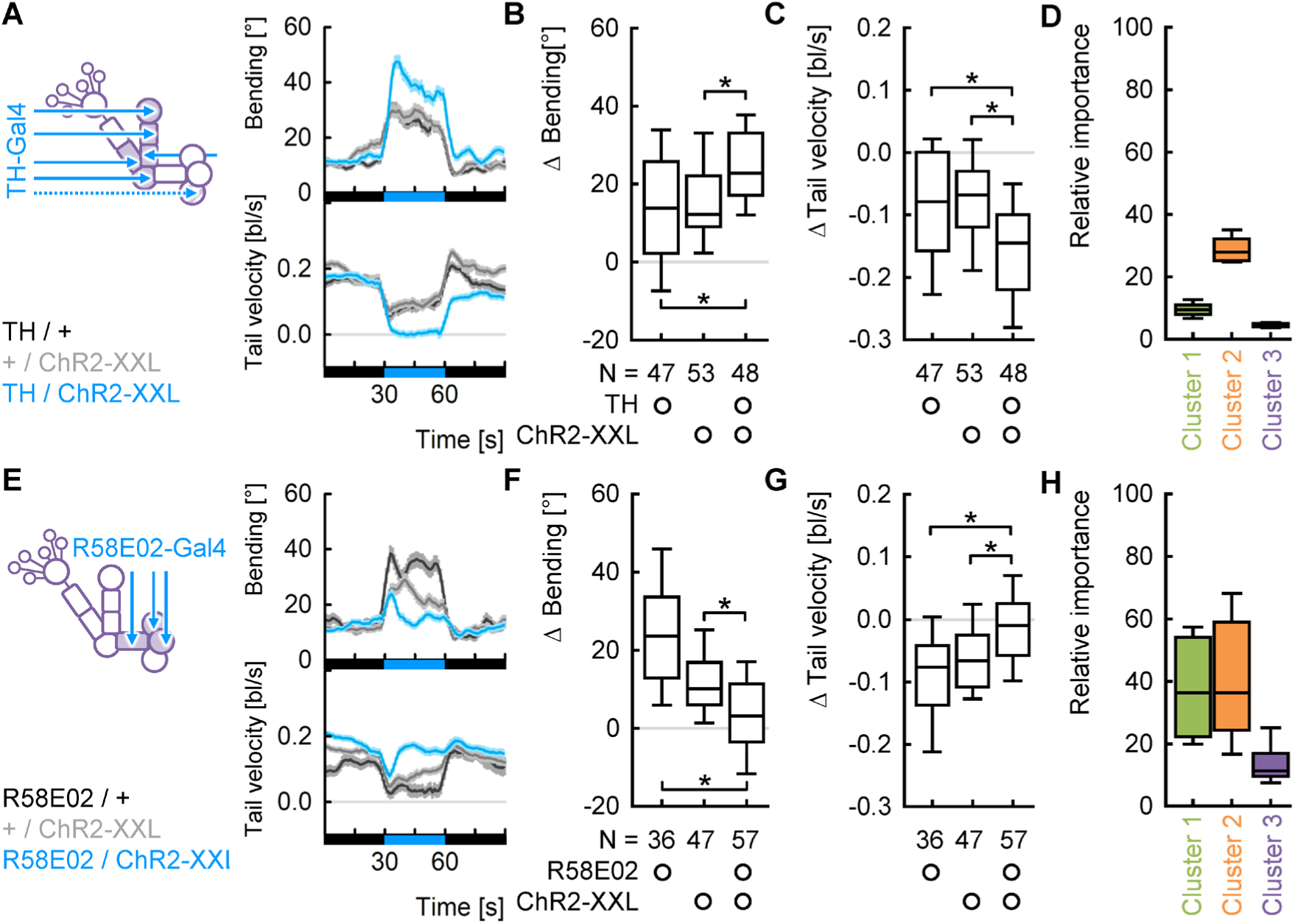
DAN activations modulate locomotion independently of light intensity. Replication of the experiments and analyses of the experiments displayed in Fig. 4D-L, but using weak blue light during the stimulations. (A,E) Average bending and tail velocity over time. (B,F) Change in bending upon light stimulation. (C,G) Change in tail velocity upon light stimulation. (D,H) Average relative importance of the behavioural attributes within each cluster, as provided by the random forest algorithm. For *TH-Gal4*, cluster 1 was less important than in the case of strong light, but the change in bending was nevertheless significant. * and ns above brackets indicate pairwise significance or non-significance, respectively (MW). For the underlying source data and the results of the statistical tests, see Supplementary Data S1.

**Figure 5 - supplement 2:**
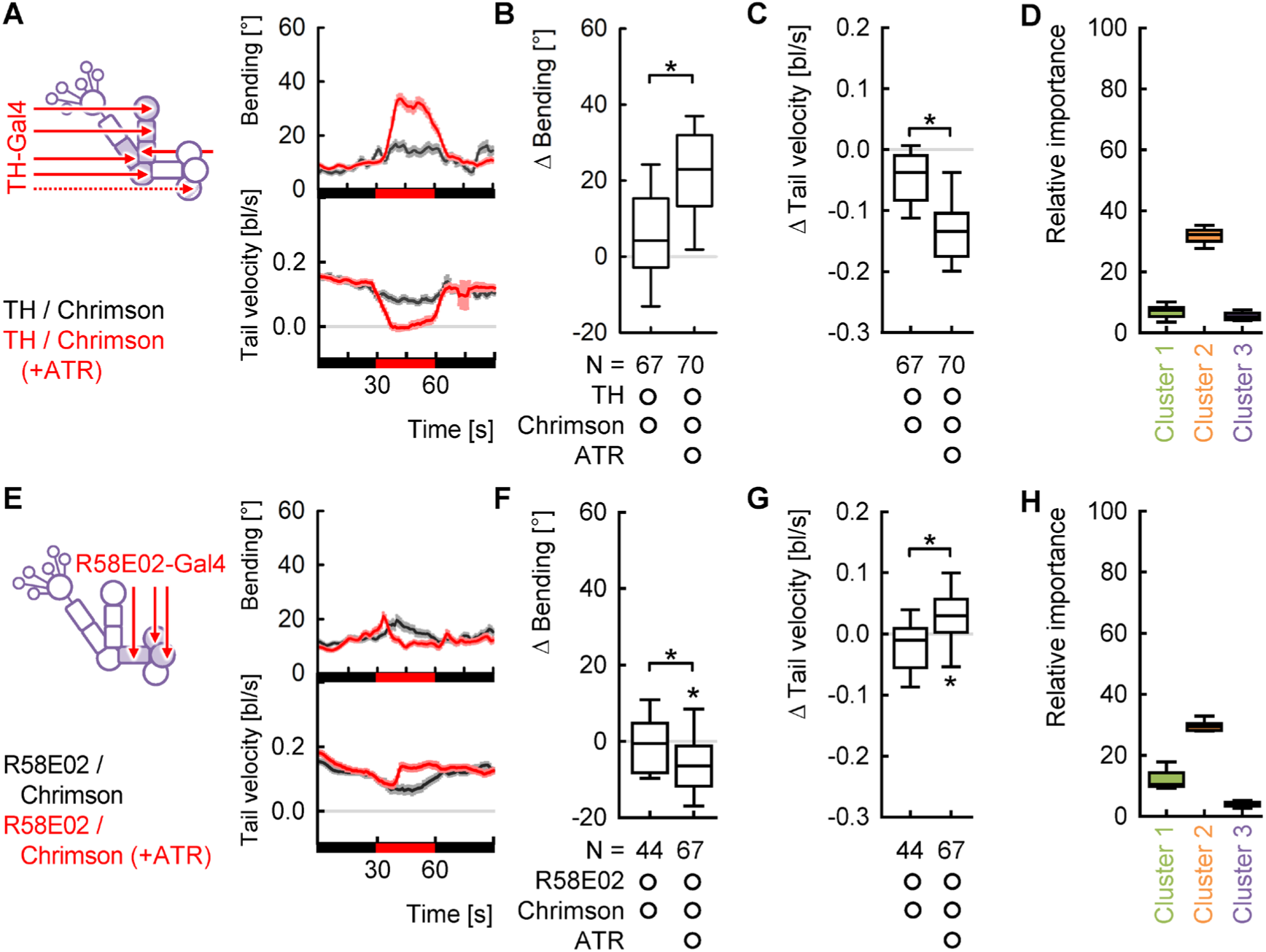
DAN activations modulate locomotion independently of effector. Replication of the experiments and analyses of the experiments displayed in Fig. 4D-L, but using CsChrimson as effector and red light stimulation. The experimental groups were larvae expressing CsChrimson in *TH-Gal4*-DANs (A-D) or *R58E02-Gal4*-DANs (E-H), fed with all-trans-retinal. The control groups were of the same genotype in standard food without all-trans-retinal. Note that these control animals behaved comparably to an effector control (Figure supplement 3), indicating that the DANs were not activated in the controls. (A,E) Average bending and tail velocity over time. (B,F) Change in bending upon light stimulation. (C,G) Change in tail velocity upon light stimulation. (D,H) Average relative importance of the behavioural attributes within each cluster, as provided by the random forest algorithm. Cluster 1 was less important than in the case of blue light, but the changes in bending was nevertheless significant. Strikingly, activating *R58E02-Gal4*-DANs decreased bending below baseline and increased velocity above baseline rather than just reverting the behaviour back to baseline. * and ns above brackets indicate pairwise significance or non-significance, respectively (MW), * below boxes indicate memory scores significantly different from zero (WSR). For the underlying source data and the results of the statistical tests, see Supplementary Data S1.

**Figure 5 - supplement 3:**
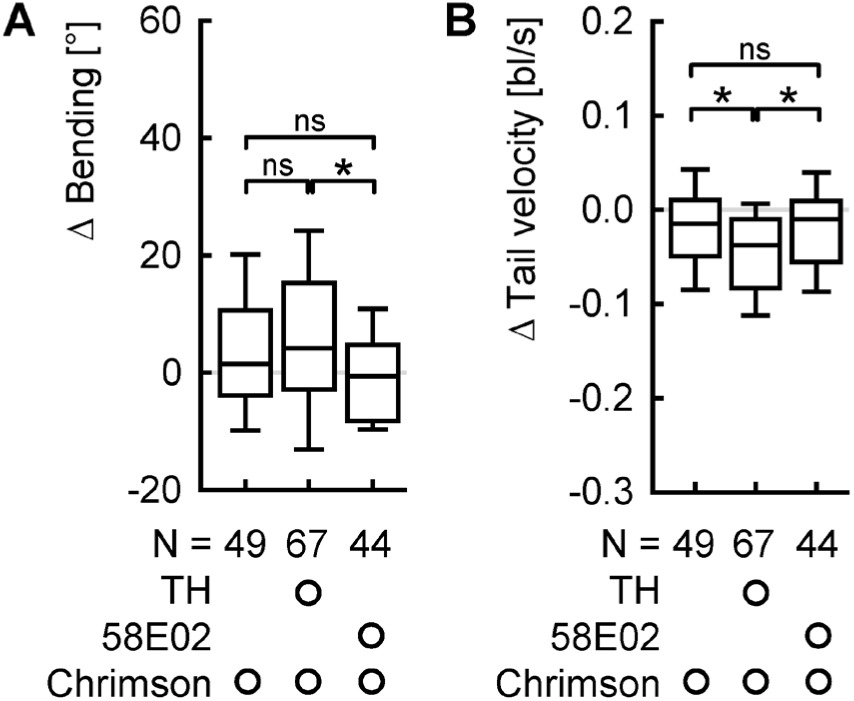
Comparison of controls for the experiments shown in Figure supplement 3. The change in (A) bending and (B) tail velocity of larvae expressing CsChrimson either under the control of *TH-Gal4* or *R58E02-Gal4* were compared with the heterozygous effector control. None of the genotypes were raised with all-trans retinal. Overall, the controls showed almost no reaction to the light. * and ns above brackets indicate pairwise significance or non-significance, respectively (MW). For the underlying source data and the results of the statistical tests, see Supplementary Data S1.

**Figure 7 - supplement 1:**
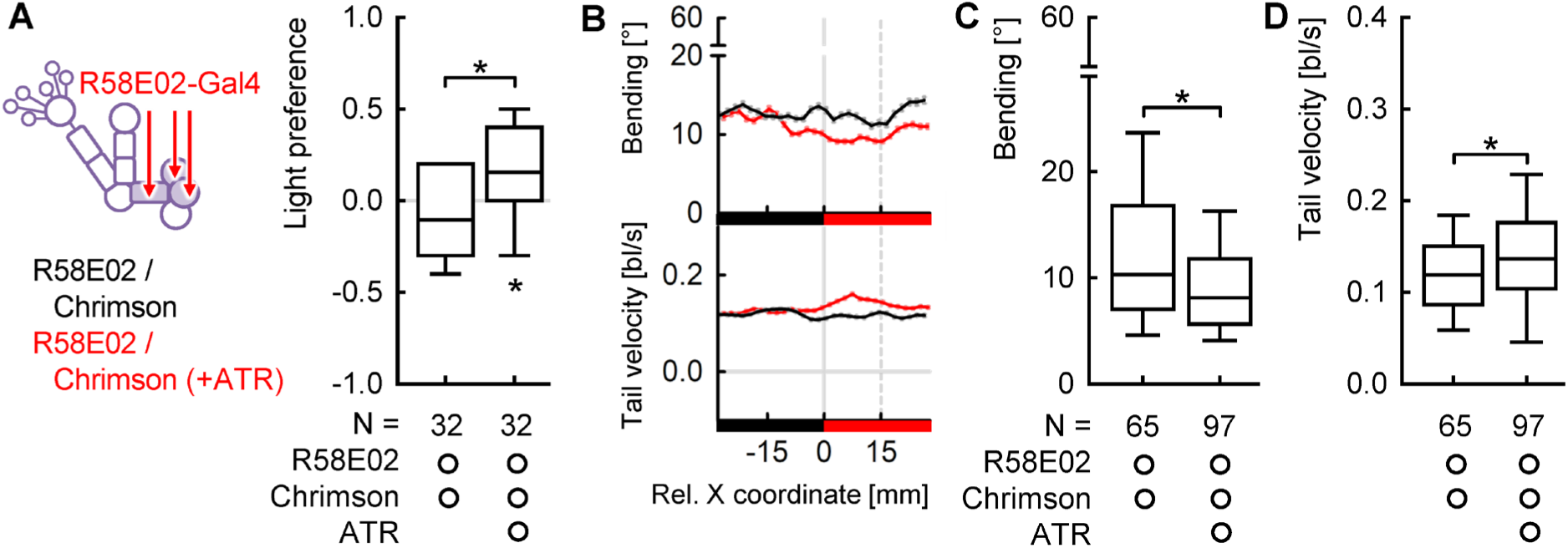
Independent repetition of the experiment of the experiment shown in Figure 7C with CsChrimson as effector. (A) The light preference after 3 minutes. Larvae raised on food with ATR showed a preference for the light whereas control larvae raised without ATR did not. This indicates that the larvae preferred to have the neurons covered by *R58E02-Gal4* activated. (B) Average bending and tail velocity of *R58E02>CsChrimson* raised with or without ATR over the X coordinate of the Petri dish, with 0 indicating the midline. The black and red bars at the bottom indicate which half was illuminated. (C) The average bending on the illuminated side of the dish. We limited the analysis to an area up to 15 mm from the midline to capture the behaviour in the choice zone. (D) As in (C), but for the tail velocity. Overall, the effects on bending and tail velocity were qualitatively comparable to the experiments with ChR2-XXL, but with a reduced magnitude (please note the truncated y axis). * above brackets indicate pairwise significance (MW), * below boxes indicate significant differences from zero (WSR). For the underlying source data and the results of the statistical tests, see Supplementary Data S1.

**Figure 8 - supplement 1:**
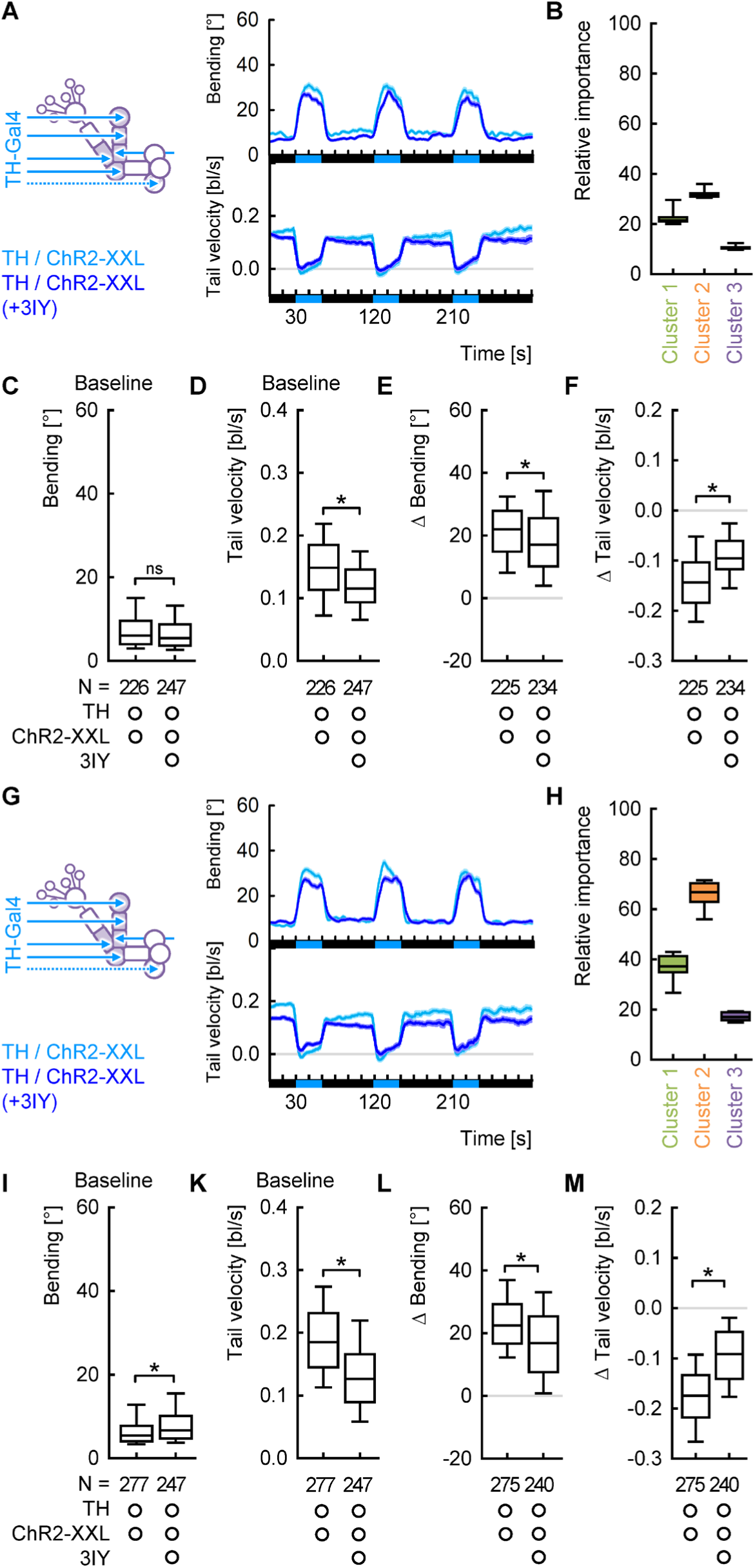
Two independent replications of the experiments shown in Figure 8H-O. (A,G) Average bending and tail velocity over time. The black and blue bars at the bottom indicate the light regimen. (B,H) The average relative importance of all behavioural attributes within each cluster, as provided by the random forest algorithm. (C,I) The animals’ average bending during the baseline period (15 to 30 s) in darkness. (D,K) As for (C,I), but for the tail velocity. Feeding 3IY reduced the tail velocity. (E,L) The change in bending upon the first light stimulation. This change was significantly decreased in animals fed with 3IY. (F,M) As in (E,L), but for the tail velocity. * and ns above brackets indicate pairwise significance or non-significance, respectively (MW). For the underlying source data and the results of the statistical tests, see Supplementary Data S1.

**Figure 9 - supplement 1:**
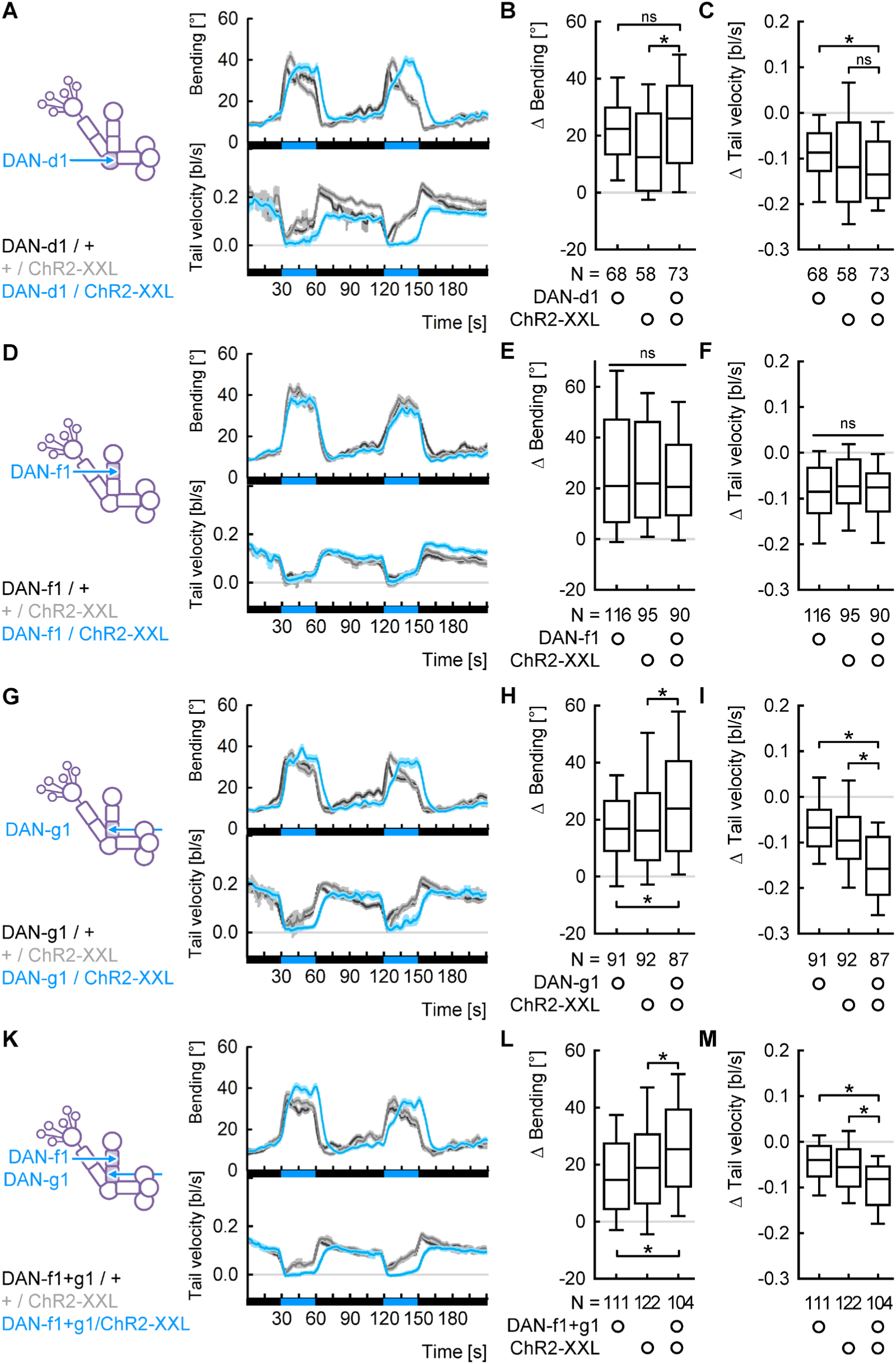
Individual DANs are sufficient to modulate bending and tail velocity. (A) Average bending and tail velocity over time. Notably, the difference between controls and experimental genotype (*DAN-d1>ChR2-XXL*) was more pronounced at the second stimulation. Partially, this was due to a more transient light response of the controls. The black and blue bars at the bottom indicate the light regimen. (B) The change in bending upon the first light stimulation. (C) As in (B), but for the tail velocity. (D-F) As for (A-C), but for DAN-f1. (G-I) As for (A-C) but for DAN-g1. (K-M) As for (A-C) but for a combination of DAN-f1 and DAN-g1. ns above a line indicates non-significance across the whole data set (KW), * and ns above brackets indicate pairwise significance or non-significance, respectively (MW). For the underlying source data and the results of the statistical tests, see Supplementary Data S1.

**Figure 9 - supplement 2:**
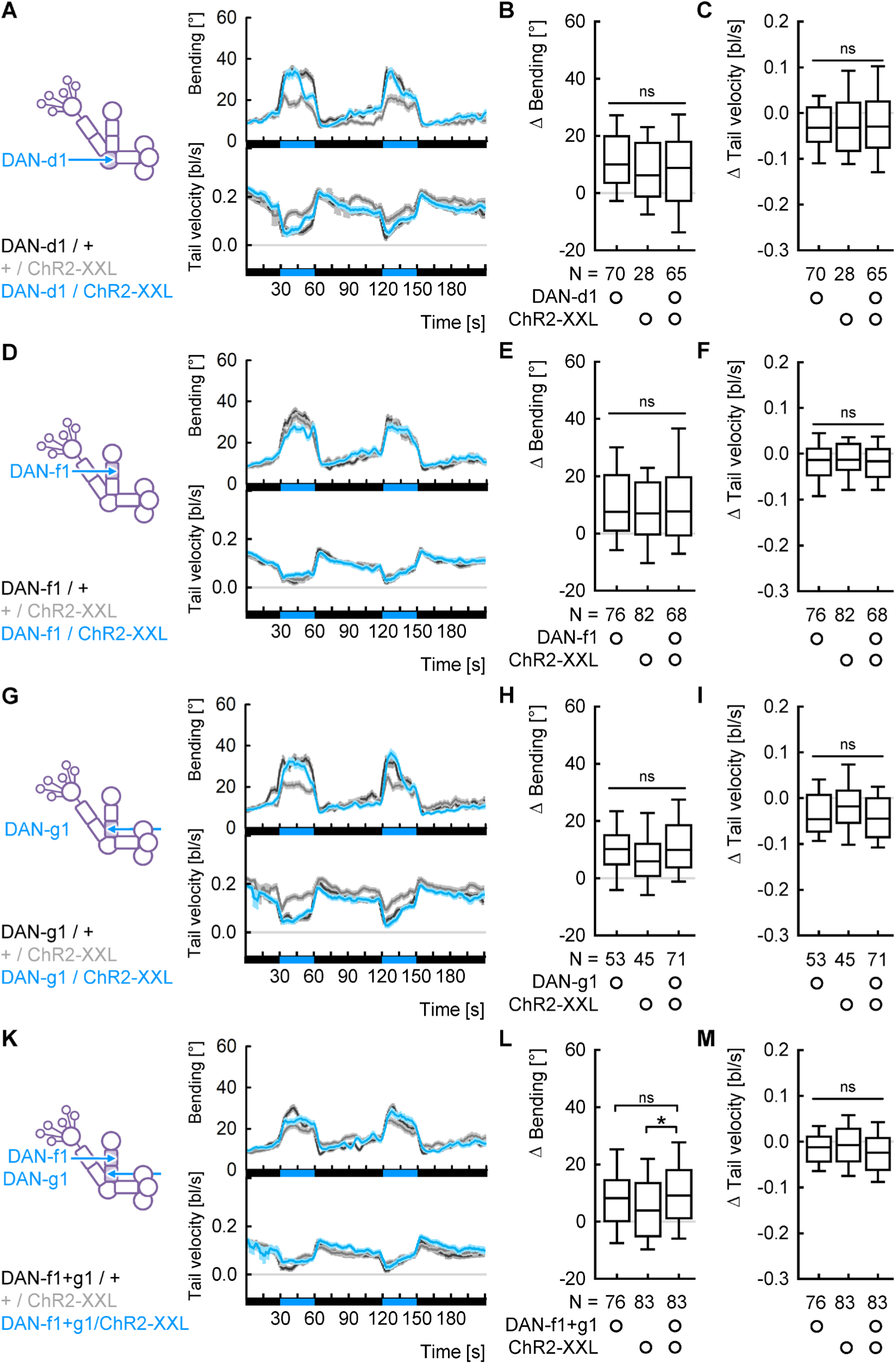
Activation of individual DANs with weak light is not sufficient to modulate locomotion. (A) Average bending and tail velocity over time. The black and blue bars at the bottom indicate the light regimen. (B) The change in bending upon the second light stimulation. There was no difference between the genotypes. (C) As in (B), but for the tail velocity. (D-F) As for (A-C), but for DAN-f1. (G-I) As for (A-C) but for DAN-g1. (K-M) As for (A-C) but for a combination of DAN-f1 and DAN-g1. ns above a line indicates non-significance across the whole data set (KW), * and ns above brackets indicate pairwise significance or non-significance, respectively (MW). For the underlying source data and the results of the statistical tests, see Supplementary Data S1.

**Figure 10 - supplement 1:**
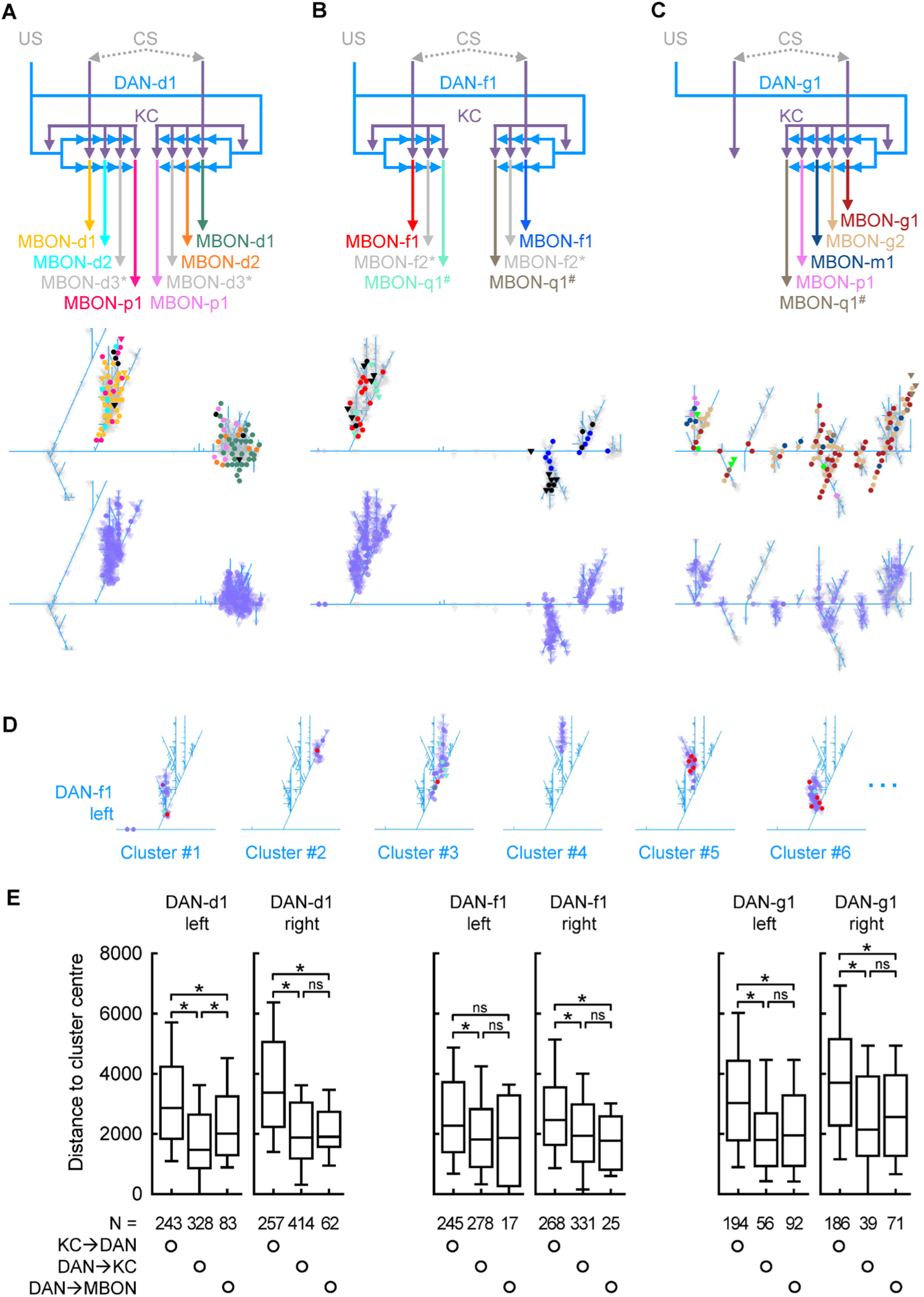
Synaptic organization of the DL1-cluster DANs. (A) Top: CS information is signalled towards the mushroom body Kenyon cells (KCs). DAN-d1 and most KCs on both hemispheres establish mutual synapses, and both DAN-d1 and KCs provide output to 4 MBONs. For simplicity, only synapses with the left DAN-d1 are shown. Middle: Truncated dendrogram of the left DAN-d1 based on EM microscopy of stage 1 larvae^19,53^. The positions of all synapses with MBONs are indicated in the colours shown in the top sketch. Bottom: The positions of all synapses with KCs are indicated in purple. (B-C) As in (A), but for DAN-f1 and DAN-g1, respectively. Note that DAN-g1 only innervates the contralateral mushroom body. *: this MBON has no synapses with the respective DAN in stage 1, but innervates the compartment in stage 3; ^#^: this MBON establishes synapses with the respective DAN in stage 1 but is not found in stage 3^19,53^. (D) The synapses are organized in clusters. Displayed are 6 sample clusters of the left DAN-f1. (E) Distances of all synapses within each cluster from the cluster centre. For all 6 DANs, KC→DAN synapses are further away from the centre than the other synapse types. * and ns above brackets indicate pairwise significance or non-significance, respectively (MW). For the underlying source data and the results of the statistical tests, see Supplementary Data 1. For high-resolution dendrograms of both the left and right DANs, see Supplementary Material S2.

## Supplementary Information

**Supplementary Data S1**: Raw data and results of statistical tests. This Excel file provides all raw data underlying the figures. Each tab shows the data of one figure or supplementary figure, with the results of all statistical tests to the left.

**Supplementary Material S1**: Full body expression of all used driver strains and genetic controls. Each page shows the three channels (autofluorescence, mCherry, mIFP) for the experimental genotype (driver strain crossed to *MB247-mCherry-CAAX, UAS-mIFP*) and a driver control lacking the UAS construct for one driver strain. On the first page, in addition the effector control crossed to *yw* is displayed. Next to the whole body, the insert shows a volume-restricted magnification of the CNS region. Blue arrowheads indicate DAN cell bodies. Scale bars represent 250 µm for the full body and 50 µm for the inserts.

**Supplementary Material S2**: High-resolution dendrograms. Each page shows the dendrogram of one DAN in high resolution. For clarity, the synapses with KCs and other synapses are displayed separate. Even pages highlight only synapses with MBONs, odd pages synapses with KCs. All other synapses are represented with transparent grey symbols. Circles display synapses from the given DAN to other neurons, triangles synapses from other neurons to that DAN. The colour code is given on each page.

**Supplementary Movie 1**: An animation through the data set used in Fig. 1E. *TH-Gal4* was crossed to *MB247-mCherry-CAAX, UAS-mIFP*, and both the mCherry-CAAX (magenta) and the mIFP (yellow) signals are shown. The movie starts with the larva’s head towards the front left. The second half shows the animal from top. At the end, the z-layers covered by the proventriculus and the melanin-producing cells were removed to allow the cells of the CNS to become visible. Grid spacing: 200 µm.

**Supplementary Movie 2**: A sample video of the experiment presented in Fig. 4 with *TH/ChR2-XXL* larvae. The video is played in double speed. Blue light (not visible in the video) was switched on at timestamp 15 s (30 s in real time) and off at timestamp 30 s (60 s in real time). Larval heads are marked in green, tails in red.

**Supplementary Movie 3**: A sample video of the experiment presented in Fig. 4 with *R58E02/ChR2-XXL* larvae. The video is played in double speed. Blue light (not visible in the video) was switched on at timestamp 15 s (30 s in real time) and off at timestamp 30 s (60 s in real time).

**Supplementary Movie 4**: A sample video of the experiment presented in Fig. 7 with *TH/ChR2-XXL* larvae. The video is played in double speed. The right half of the dish was illuminated with blue light (not visible in the video) throughout the experiment.

**Supplementary Movie 5**: A sample video of the experiment presented in Fig. 7 with *R58E02/ChR2-XXL* larvae. The video is played in double speed. The right half of the dish was illuminated with blue light (not visible in the video) throughout the experiment.

