## Supplementary Material S1 for "Individual dopaminergic neurons induce unique, yet overlapping combinations of behavioural modulations including safety learning, memory retrieval and acute locomotion"

Full body expression of all used driver strains and genetic controls. Each page shows the three channels (autofluorescence, mCherry, mIFP) for the experimental genotype (driver strain crossed to *MB247-mCherry-CAAX, UAS-mIFP*) and a driver control lacking the UAS construct for one driver strain. On the first page, in addition the effector control crossed to *yw* is displayed. Next to the whole body, the insert shows a volume-restricted magnification of the CNS region. Blue arrowheads indicate DAN cell bodies. Scale bars represent 250 µm for the full body and 50 µm for the inserts.

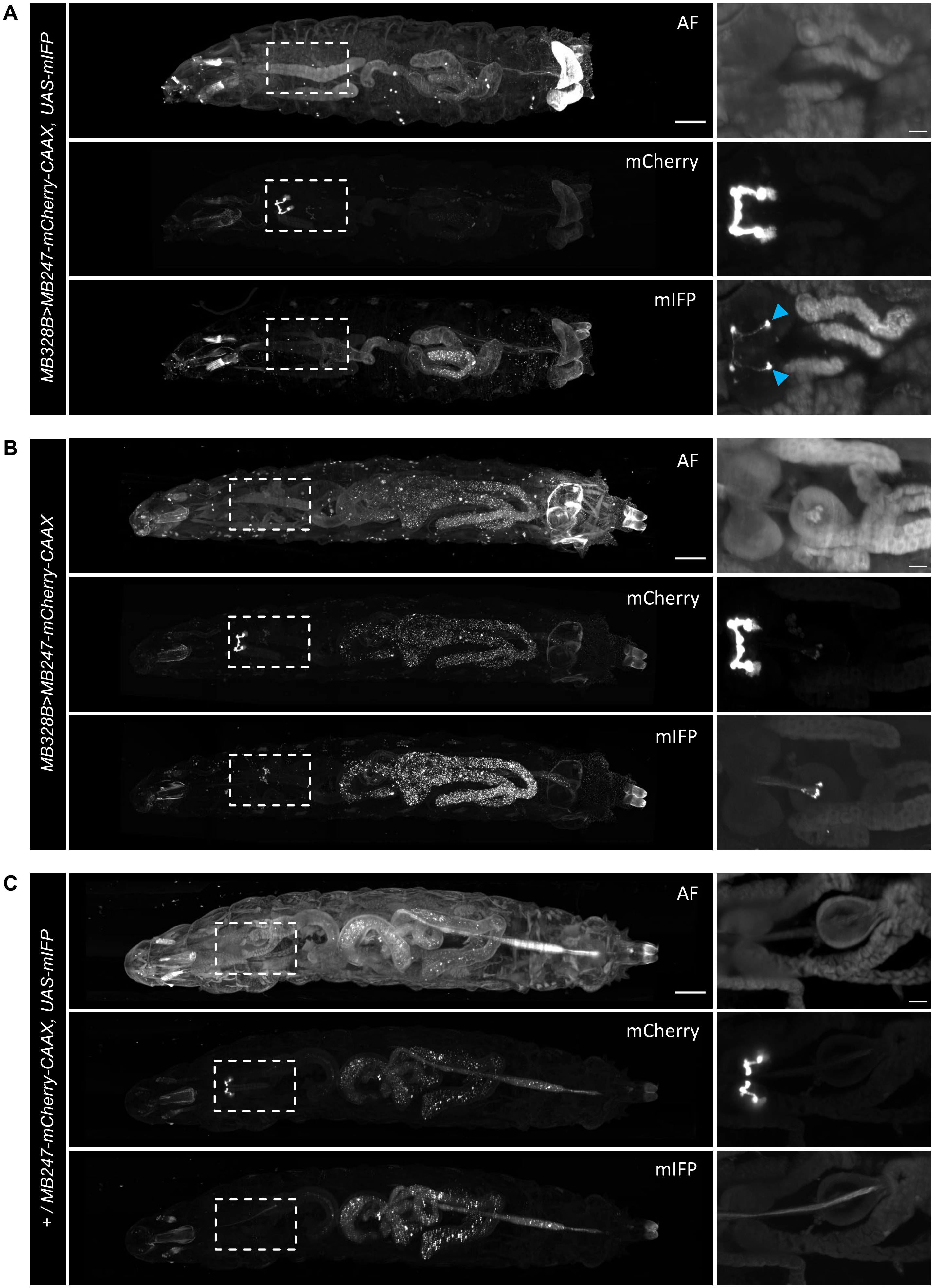
A
MB328B>MB247-mCherry-CAAX, UAS-mIFP
AF
mCherry
mIFP
B
MB328B>MB247-mCherry-CAAX
AF
mCherry
mIFP
C
+ / MB247-mCherry-CAAX, UAS-mIFP
AF
mCherry
mIFP

D

E

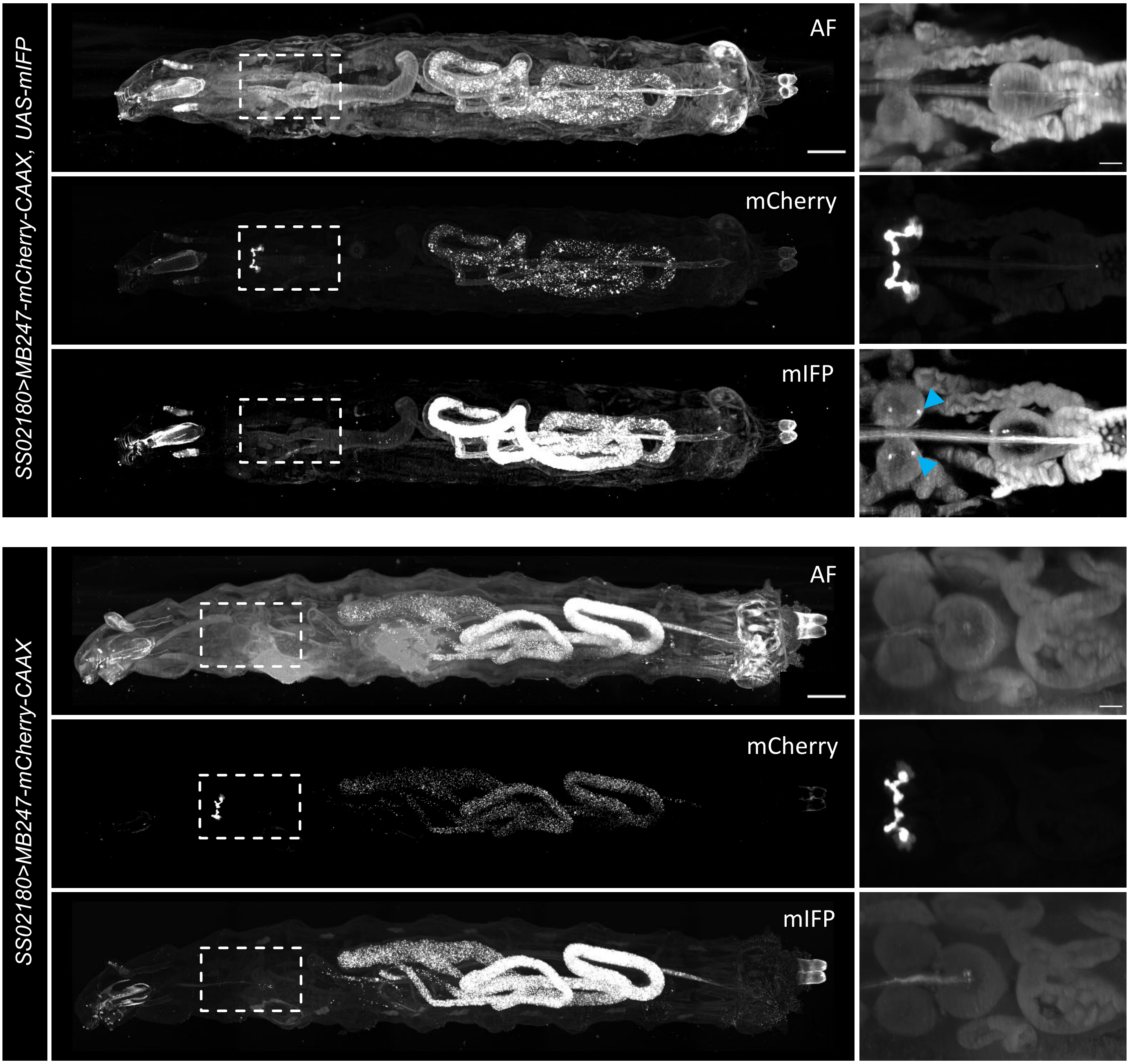

SS02180>MB247-mCherry-CAAX, UAS-mIFP
AF
mCherry
mIFP
SS02180>MB247-mCherry-CAAX
AF
mCherry
mIFP

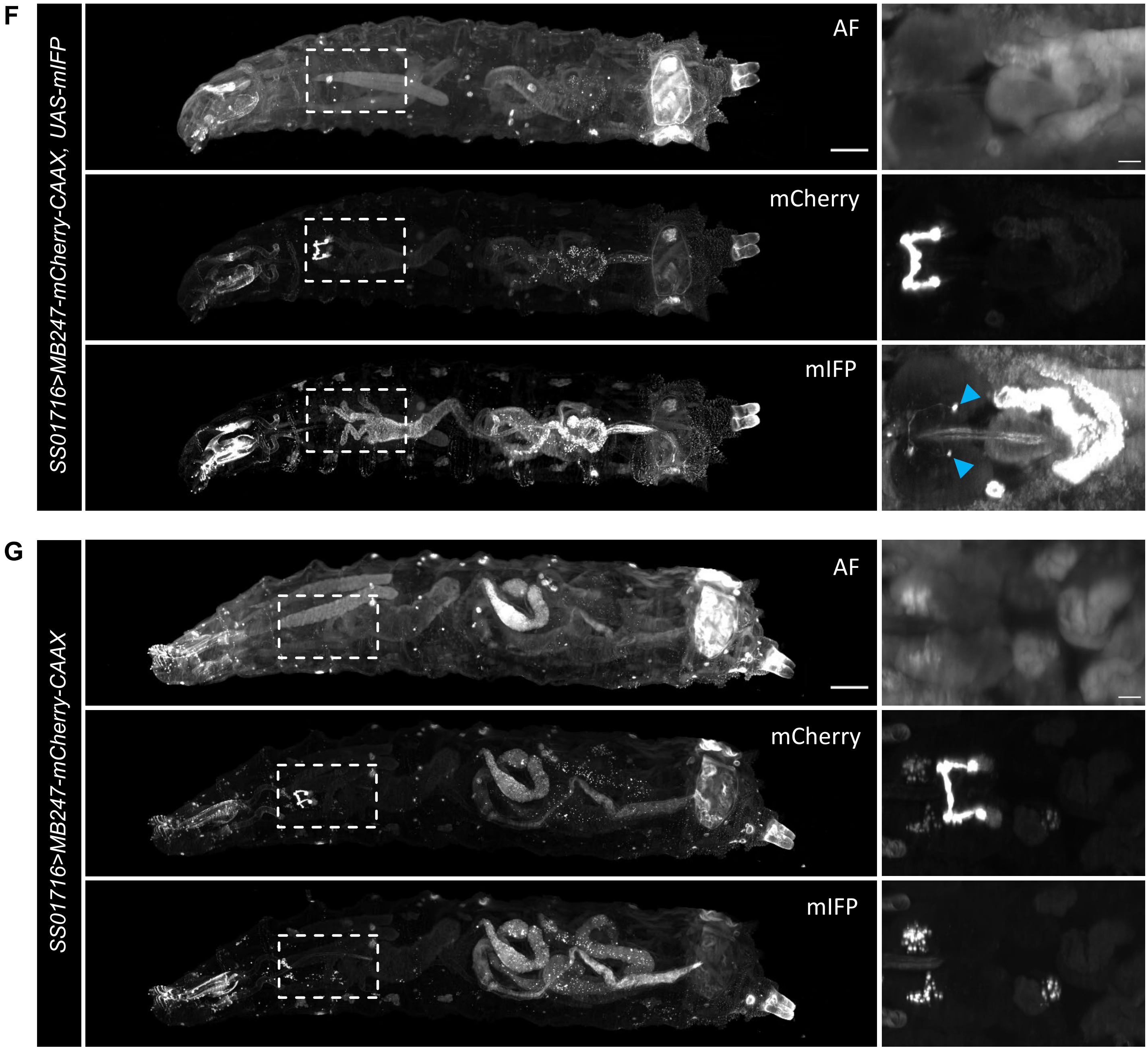
F
SS01716>MB247-mCherry-CAAX, UAS-mIFP
AF
mCherry
mIFP
G
SS01716>MB247-mCherry-CAAX
AF
mCherry
mIFP

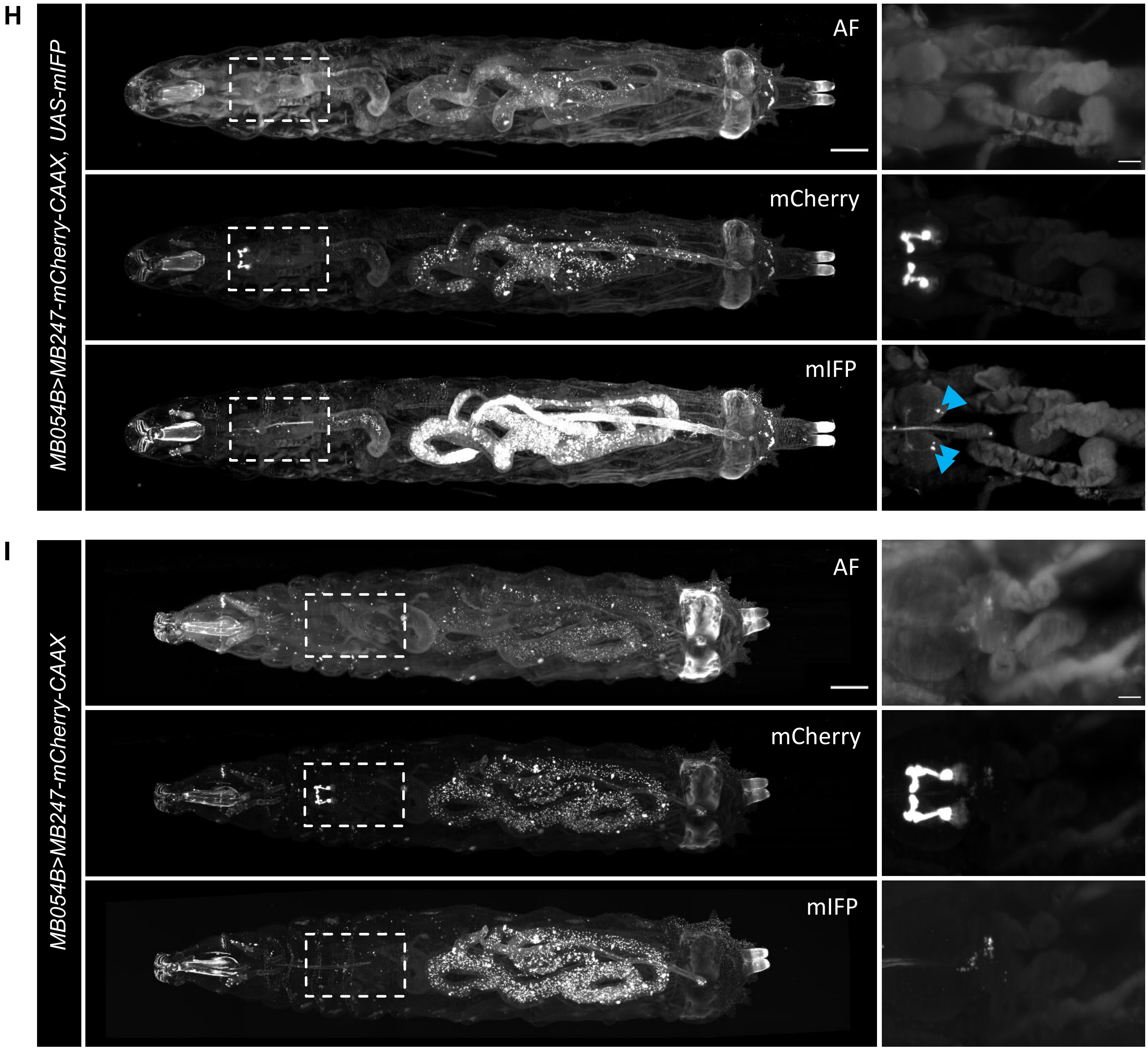
H
MB054B>MB247-mCherry-CAAX, UAS-mIFP
AF
mCherry
mIFP
I
MB054B>MB247-mCherry-CAAX
AF
mCherry
mIFP

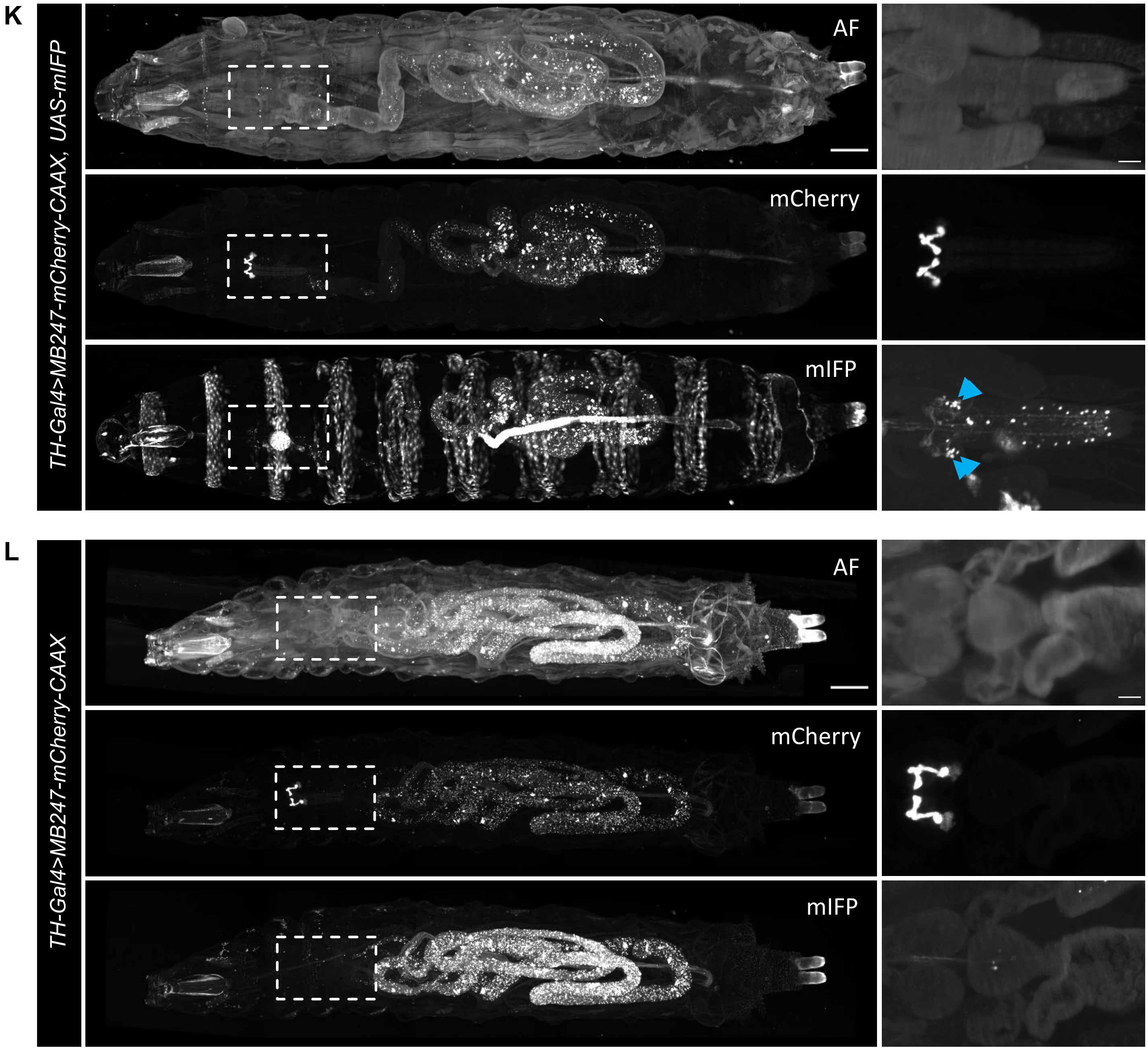
K
TH-Gal4>MB247-mCherry-CAAX, UAS-mIFP
AF
mCherry
mIFP
L
TH-Gal4>MB247-mCherry-CAAX
AF
mCherry
mIFP

M

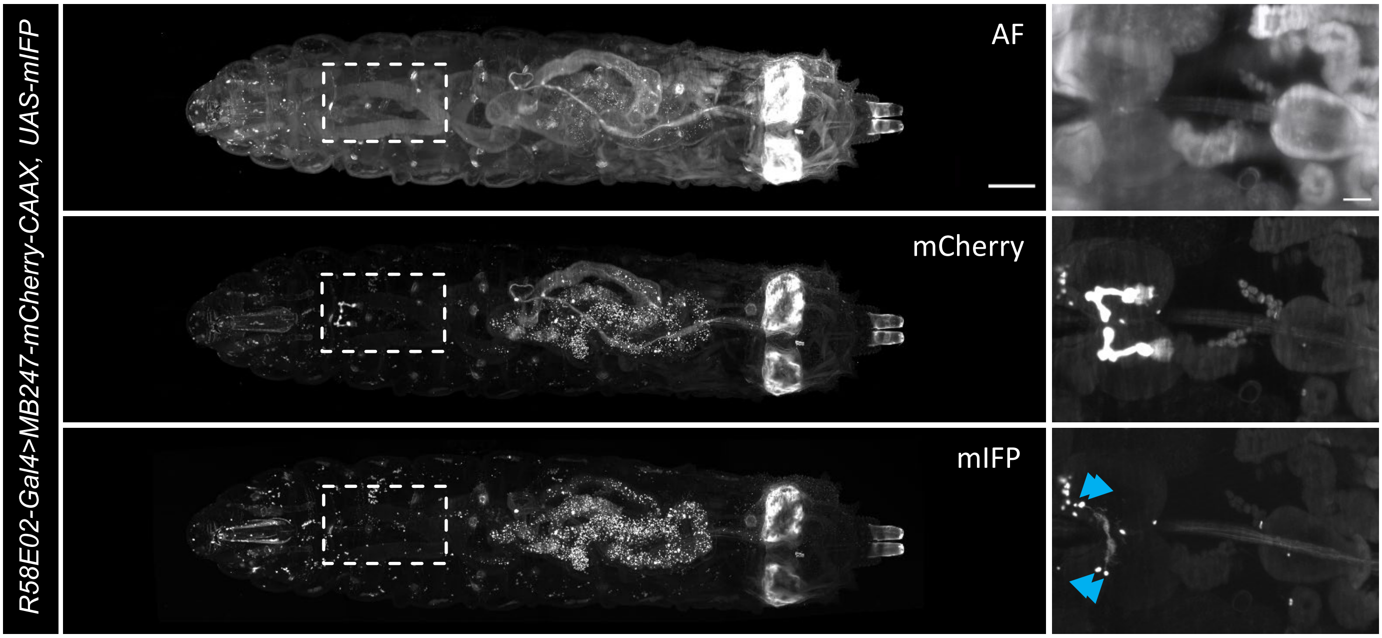
R58E02-Gal4>MB247-mCherry-CAAX, UAS-mIFP
AF
mCherry
mIFP

N

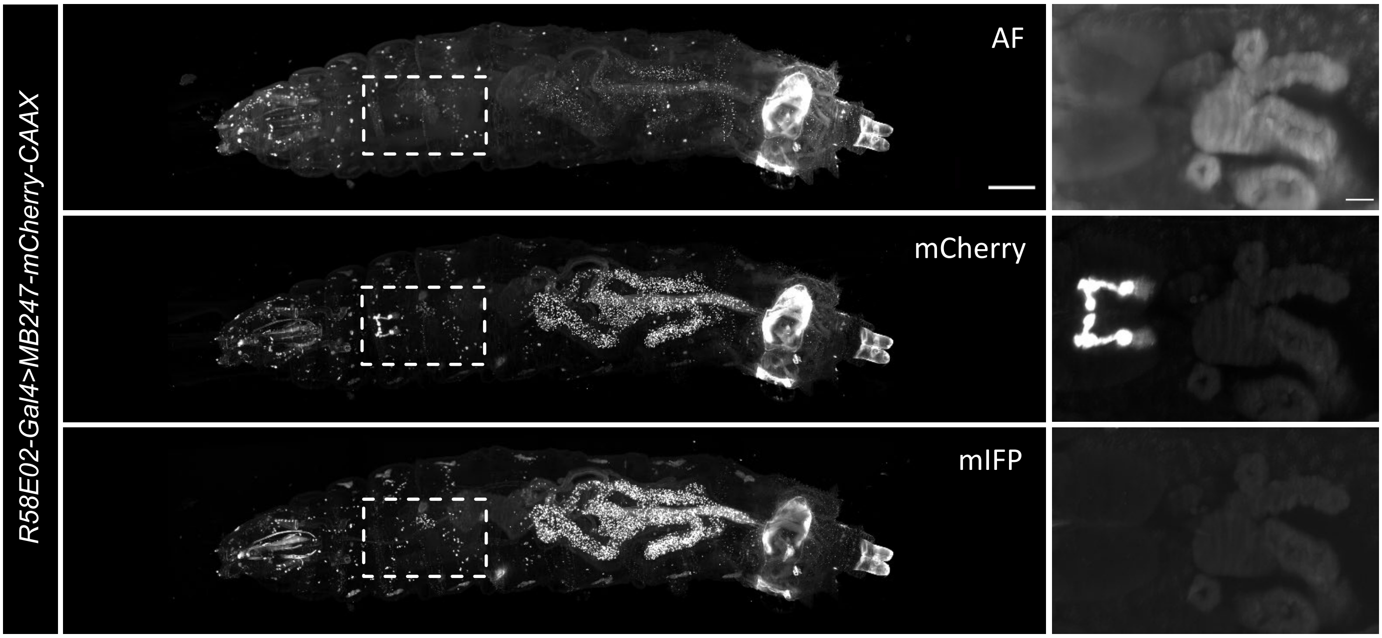
R58E02-Gal4>MB247-mCherry-CAAX
AF
mCherry
mIFP
