## Supplementary Material S2 for "Individual dopaminergic neurons induce unique, yet overlapping combinations of behavioural modulations including safety learning, memory retrieval and acute locomotion"

High-resolution dendrograms. Each page shows the dendrogram of one DAN in high resolution. For clarity, the synapses with KCs and other synapses are displayed separate. Even pages highlight only synapses with MBONs, odd pages synapses with KCs. All other synapses are represented with transparent grey symbols. Circles display synapses from the given DAN to other neurons, triangles synapses from other neurons to that DAN. The colour code is given on each page.

### DAN−d1 left

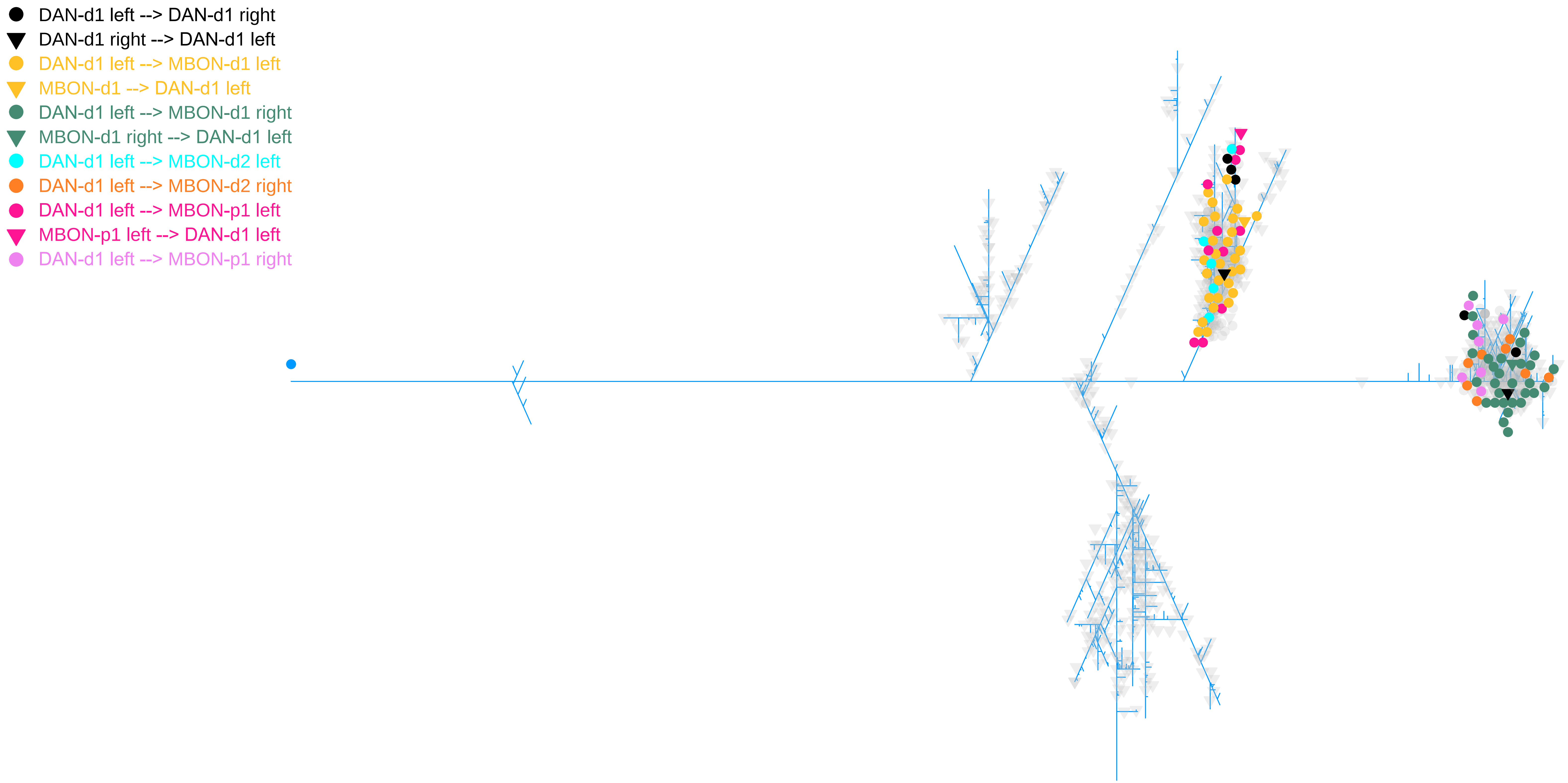
DAN-d1 left --> DAN-d1 right
DAN-d1 right --> DAN-d1 left
DAN-d1 left --> MBON-d1 left
MBON-d1 --> DAN-d1 left
DAN-d1 left --> MBON-d1 right
MBON-d1 right --> DAN-d1 left
DAN-d1 left --> MBON-d2 left
DAN-d1 left --> MBON-d2 right
DAN-d1 left --> MBON-p1 left
MBON-p1 left --> DAN-d1 left
DAN-d1 left --> MBON-p1 right

#### DAN−d1 left

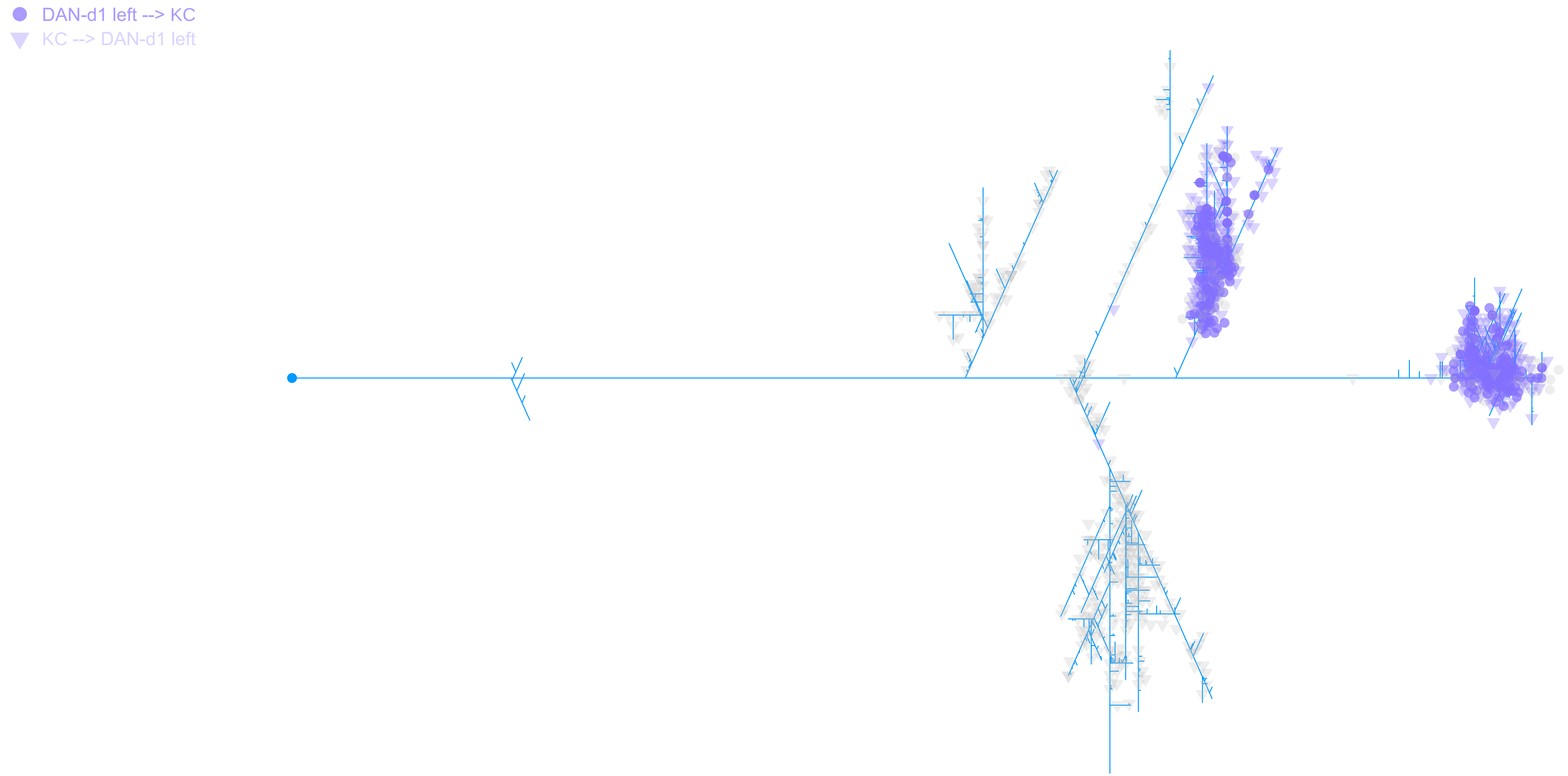

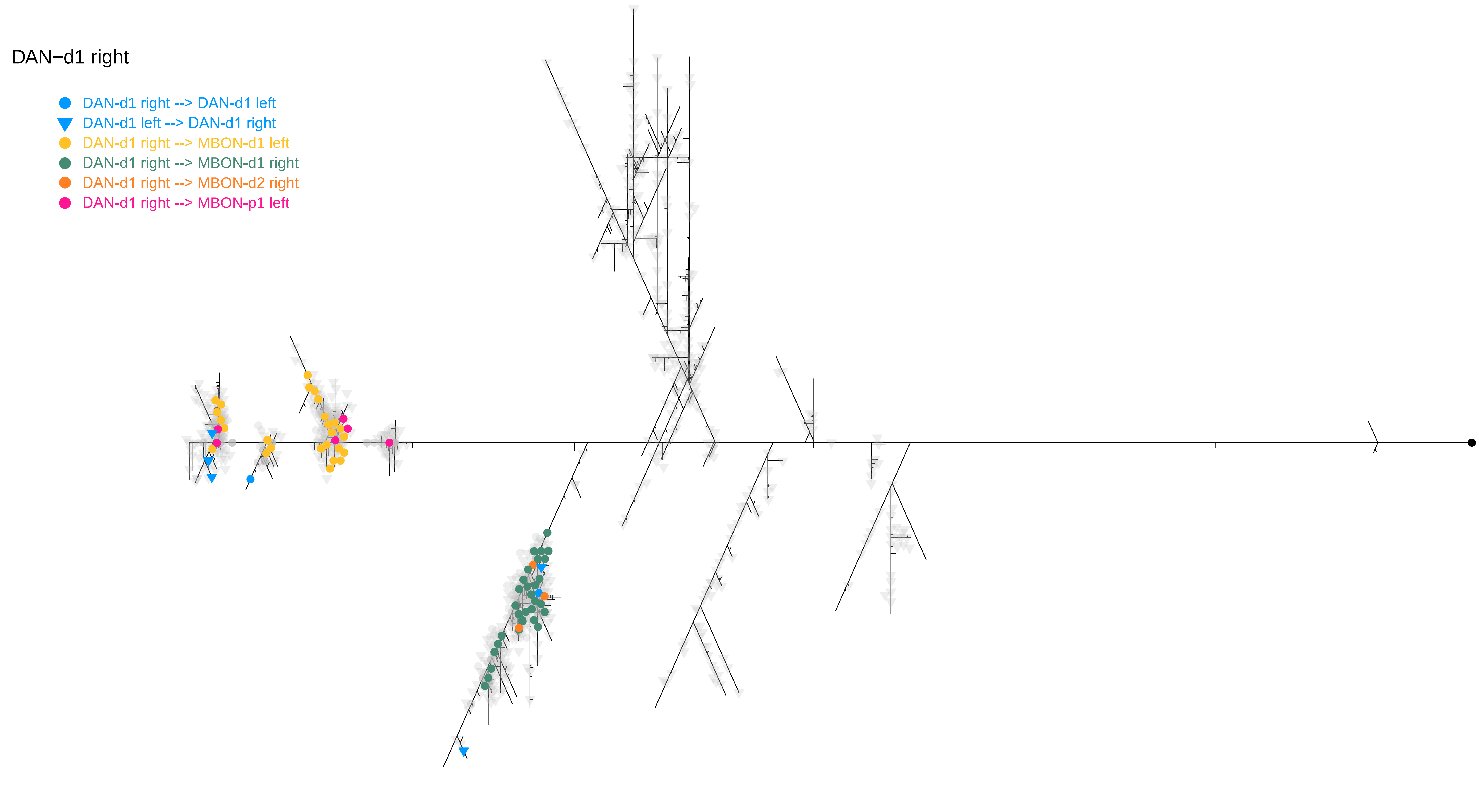

DAN−d1 right
DAN-d1 right --> DAN-d1 left
DAN-d1 left --> DAN-d1 right
DAN-d1 right --> MBON-d1 left
DAN-d1 right --> MBON-d1 right
DAN-d1 right --> MBON-d2 right
DAN-d1 right --> MBON-p1 left

DAN−d1 right

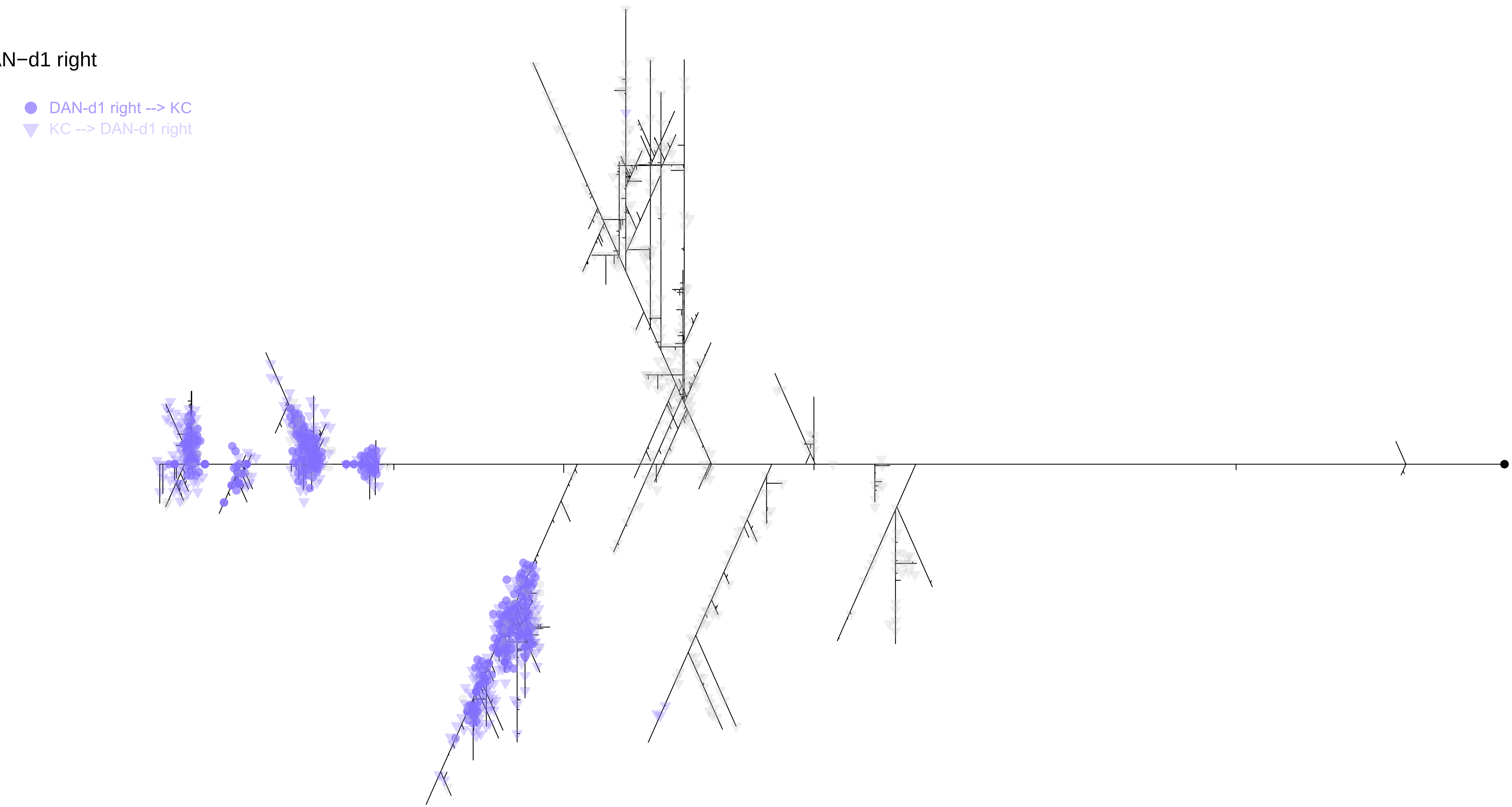

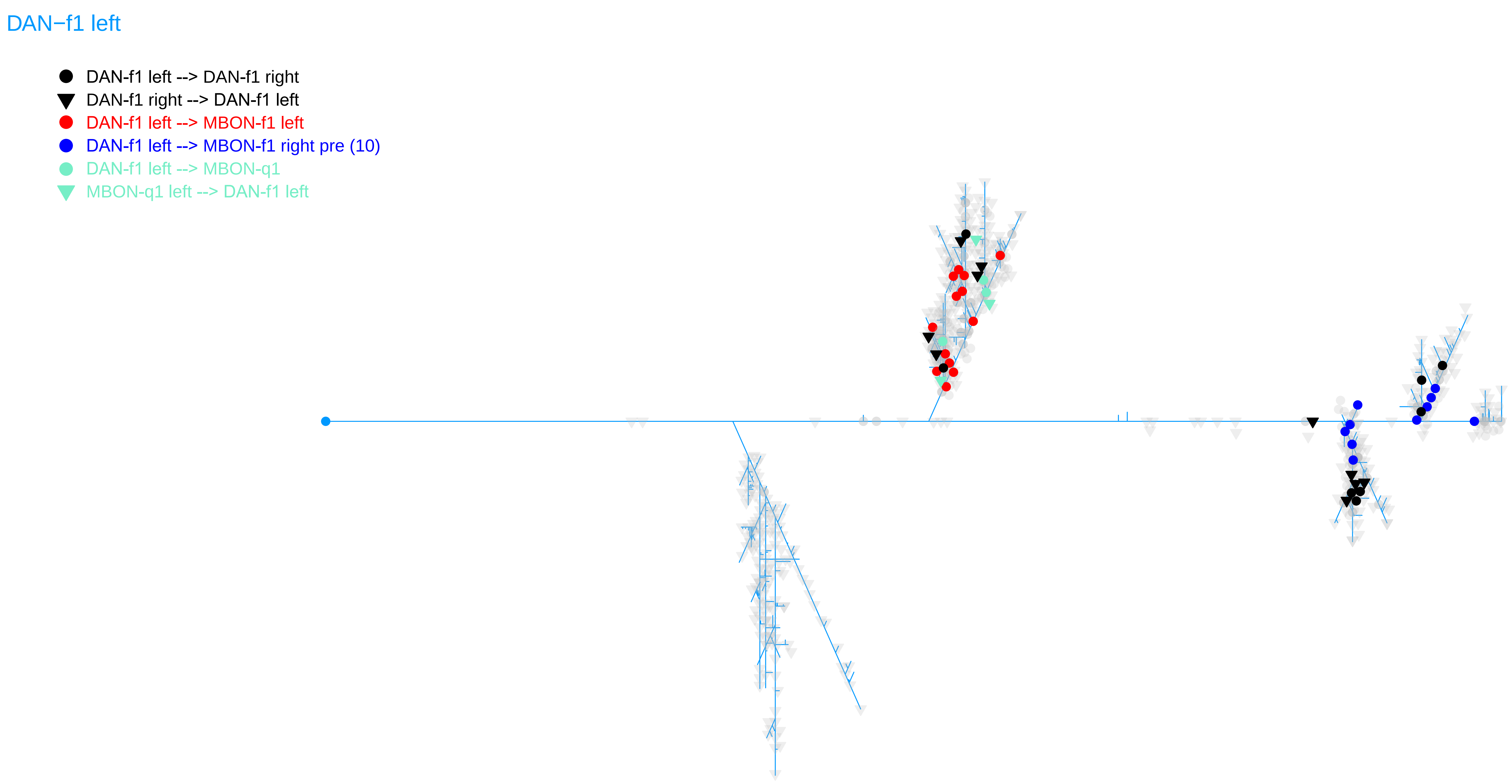
DAN−f1 left
DAN-f1 left --> DAN-f1 right
DAN-f1 right --> DAN-f1 left
DAN-f1 left --> MBON-f1 left
DAN-f1 left --> MBON-f1 right pre (10)
DAN-f1 left --> MBON-q1
MBON-q1 left --> DAN-f1 left

DAN−f1 left

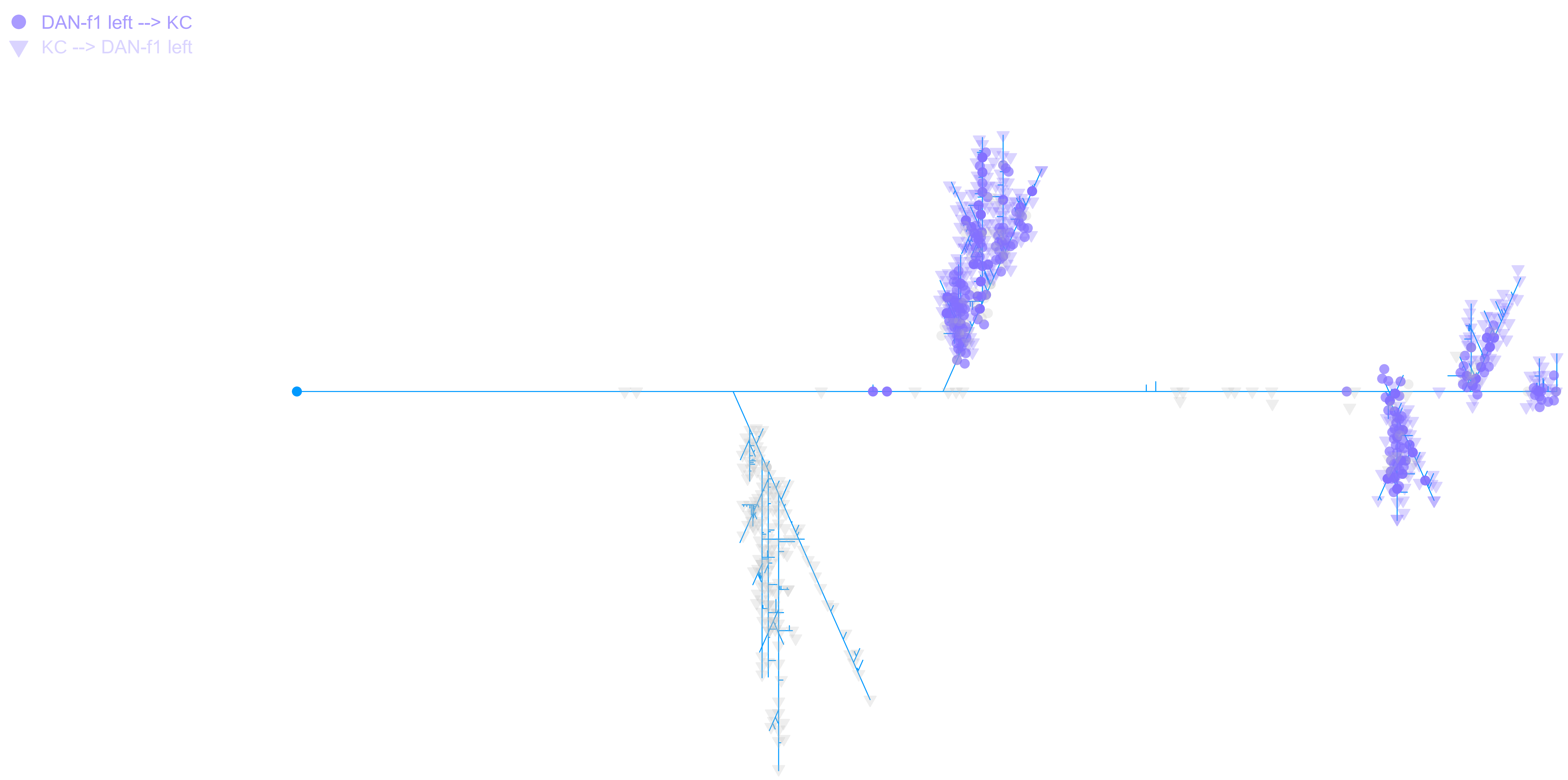

DAN−f1 right

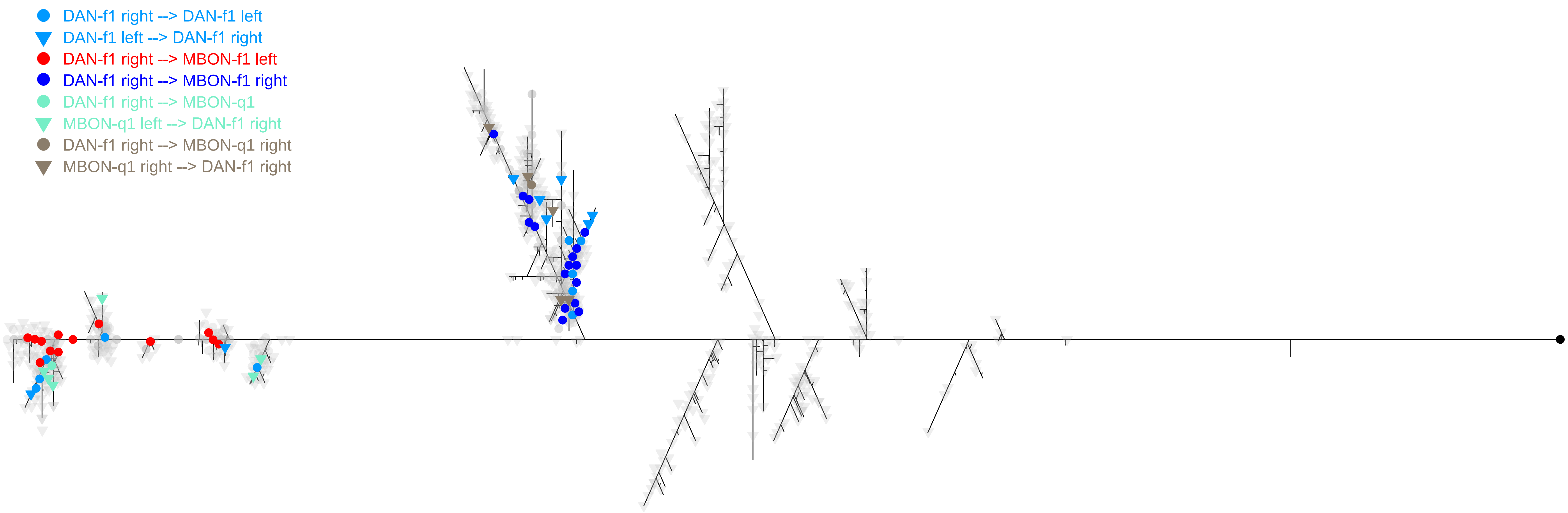

DAN-f1 right --> DAN-f1 left
DAN-f1 left --> DAN-f1 right
DAN-f1 right --> MBON-f1 left
DAN-f1 right --> MBON-f1 right
DAN-f1 right --> MBON-q1
MBON-q1 left --> DAN-f1 right
DAN-f1 right --> MBON-q1 right
MBON-q1 right --> DAN-f1 right

DAN−f1 right

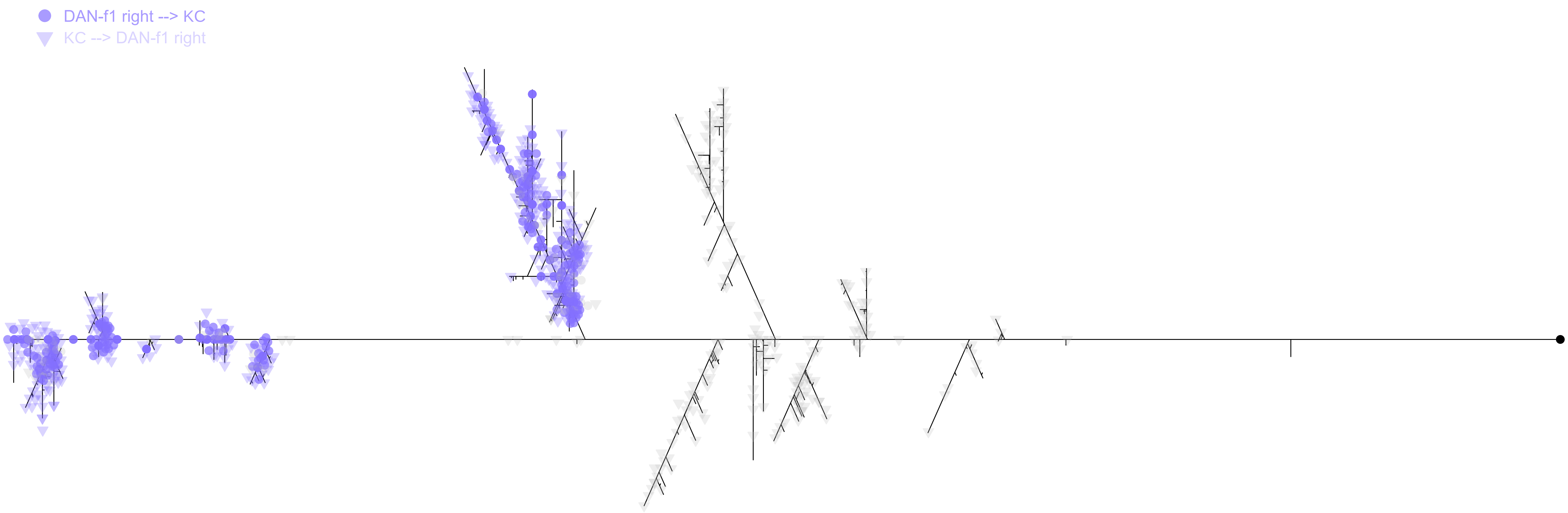

DAN−g1 left

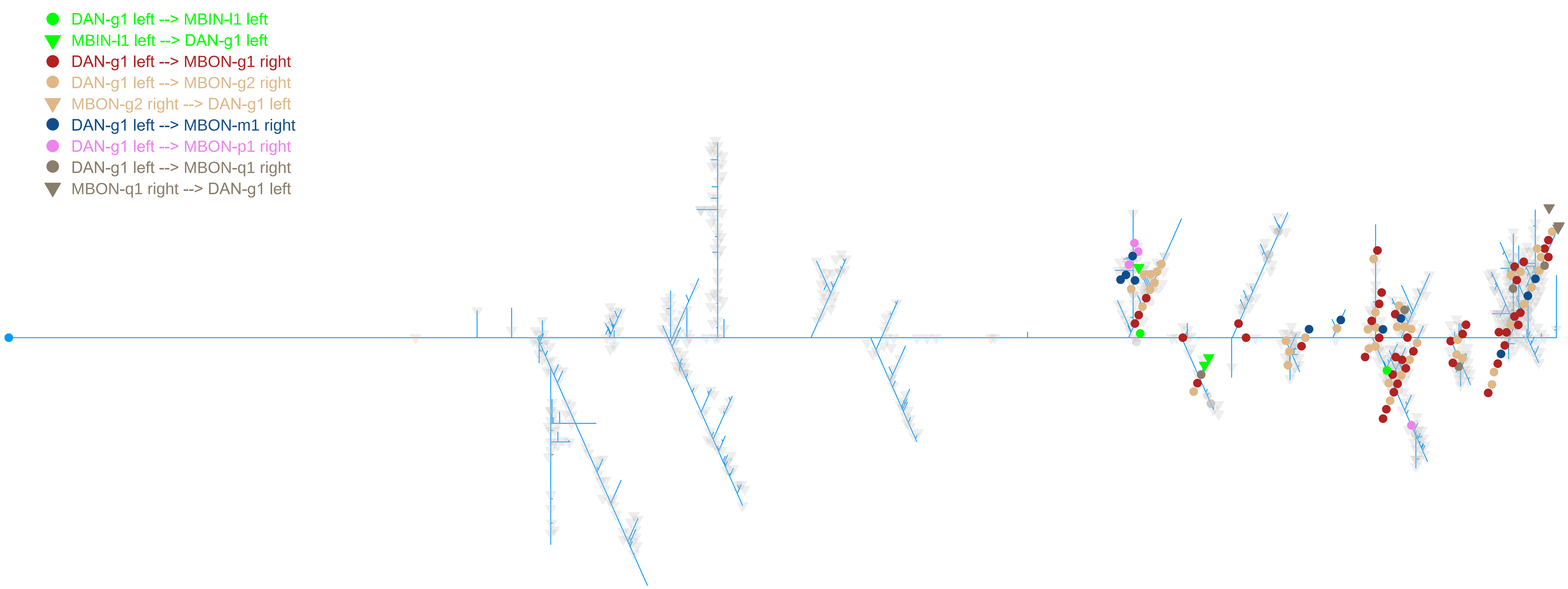
DAN-g1 left --> MBIN-l1 left
MBIN-l1 left --> DAN-g1 left
DAN-g1 left --> MBON-g1 right
DAN-g1 left --> MBON-g2 right
MBON-g2 right --> DAN-g1 left
DAN-g1 left --> MBON-m1 right
DAN-g1 left --> MBON-p1 right
DAN-g1 left --> MBON-q1 right
MBON-q1 right --> DAN-g1 left

DAN−g1 left

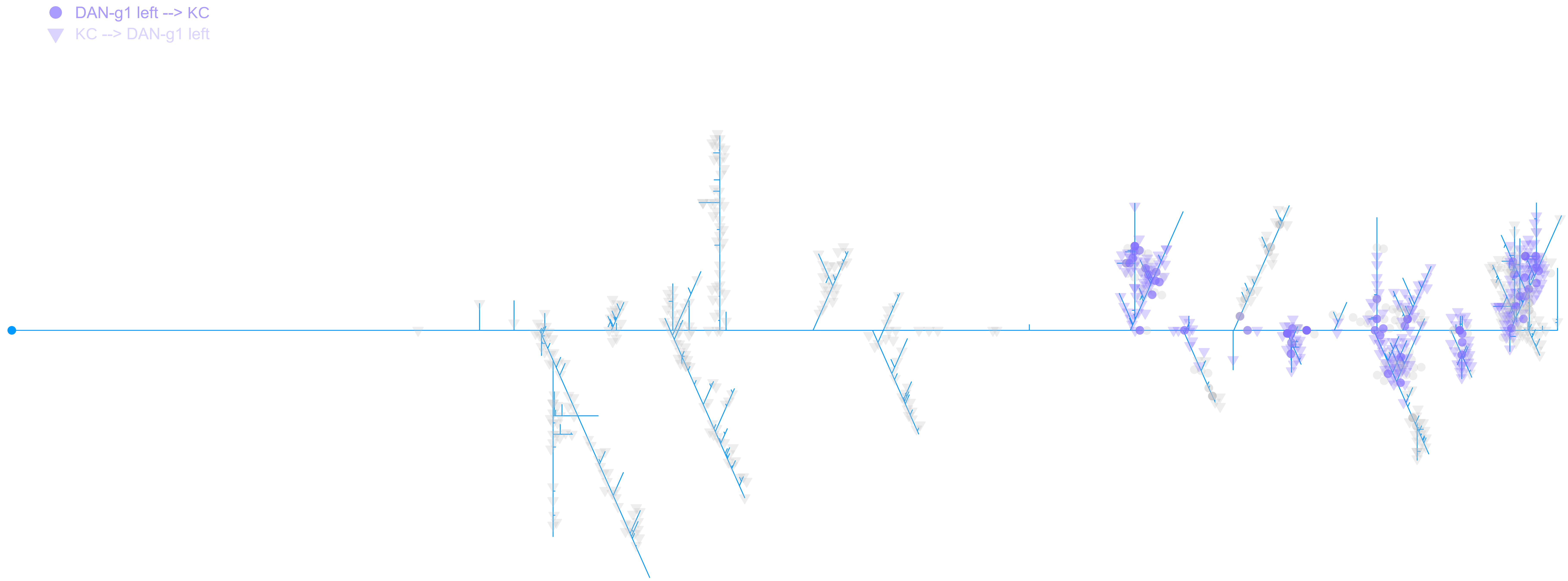

DAN−g1 right

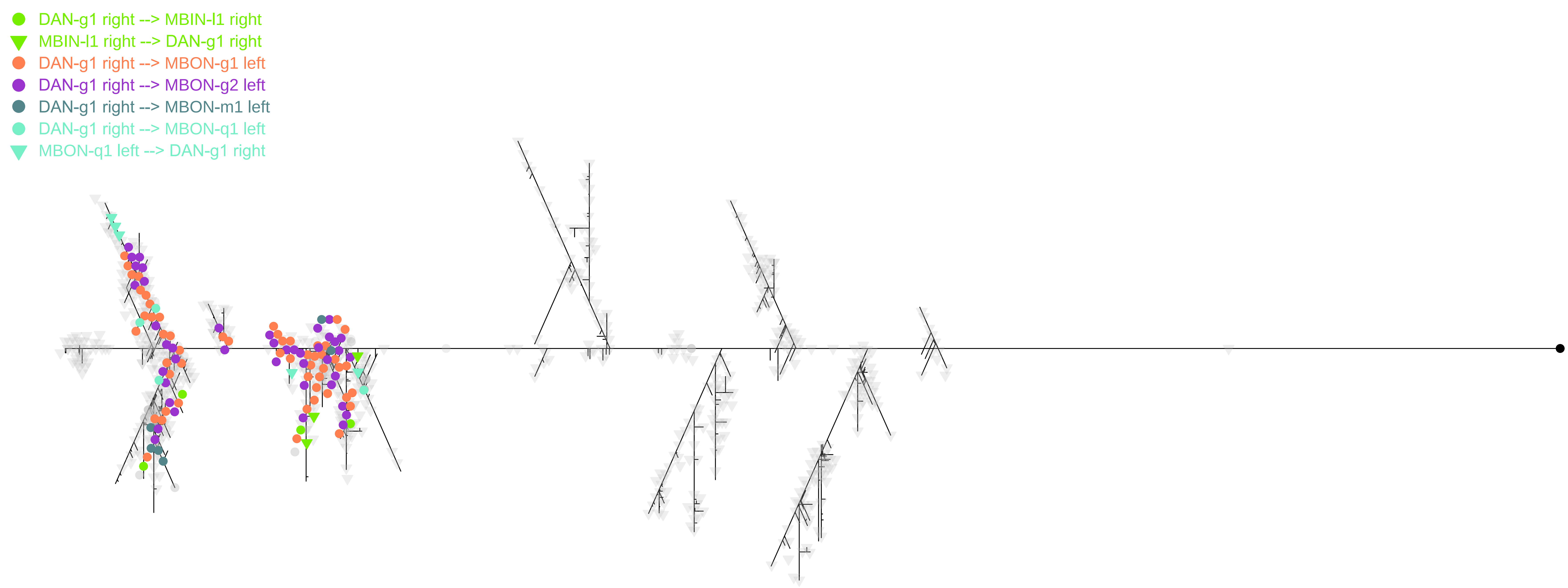
DAN-g1 right --> MBIN-l1 right
MBIN-l1 right --> DAN-g1 right
DAN-g1 right --> MBON-g1 left
DAN-g1 right --> MBON-g2 left
DAN-g1 right --> MBON-m1 left
DAN-g1 right --> MBON-q1 left
MBON-q1 left --> DAN-g1 right

DAN−g1 right

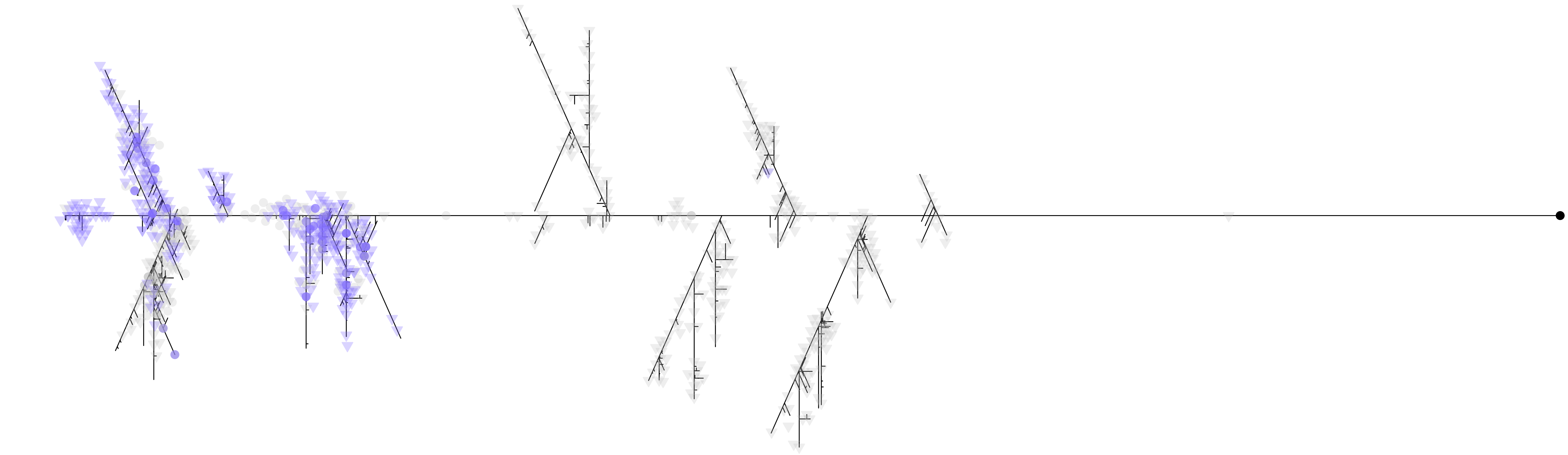
